# Comparison Between Lotka-Volterra and Multivariate Autoregressive Models of Ecological Interaction Systems

**DOI:** 10.1101/2021.10.07.463461

**Authors:** Daniel V. Olivença, Jacob D. Davis, Eberhard O. Voit

## Abstract

1. Lotka-Volterra (LV) and Multivariate Autoregressive (MAR) models are computational frameworks with different mathematical structures that have both been proposed for the same purpose of extracting governing features of dynamic interactions among coexisting populations of different species from observed time series data.
2. We systematically compare the feasibility of the two modeling approaches, using four synthetically generated datasets and seven ecological datasets from the literature.
3. The overarching result is that LV models outperform MAR models in most cases and are generally superior for representing cases where the dependent variables deviate greatly from their steady states. A large dynamic range is particularly prevalent when the populations are highly abundant, change considerably over time, and exhibit a large signal-to-noise ratio. By contrast, MAR models are better suited for analyses of populations with low abundances and for investigations where the quantification of noise is important.
4. We conclude that the choice of either one or the other modeling framework should be guided by the specific goals of the analysis and the dynamic features of the data.

**Availability of algorithms used:** https://github.com/LBSA-VoitLab/Comparison-Between-LV-and-MAR-Models-of-Ecological-Interaction-Systems

## 1. Introduction

The growth of populations has been a topic of human interest since prehistoric times: Babylonian clay tablets documented exponential growth in cuneiform lettering as early as about 4,000 years ago (Sachs & Goetze, 1945; Savageau, 1979). The quantitative representation and analysis of population dynamics is also one of the early roots of mathematical modeling in biology. In the mid-1920s, Alfred Lotka (Lotka, 1925) studied periodic increases and decreases in the populations of lynx and hare in Canada, while Vito Volterra (Volterra, 1926) independently analyzed fish catches and the competition among populations in the Adriatic Sea. Since these early days, Lotka-Volterra (LV) models have become a mainstay—and typical default—in computational ecology (May, 2001).

With the discovery of complex microbiomes and their surprisingly strong effects on human health and the environment, the interest in interactions among different species has received renewed attention (Gavin, Pokrovskii, Prentice, & Sobolev, 2006; Stein et al., 2013; Shenhav et al., 2019). As an example, we recently inferred the temporally changing interactions among bacterial communities in different lake environments with over 12,000 Operational Taxonomic Units (OTUs) (Dam et al., 2016; Dam et al., 2020). We chose as our computational framework an LV model, which we augmented with LV equations for environmental variables that affected the OTUs (see also (Stein et al., 2013)). Our rationale for this choice was a combination of (1) the successful history of LV models, (2) their mathematical simplicity and tractability and (3) the important fact that parameter values (and thus signs and strengths of interactions) can be obtained from time series data of OTU abundances with methods of linear regression (Voit & Chou, 2010).

Multivariate Autoregressive (MAR) models were proposed a few decades ago as a viable alternative to LV models. Originally proposed for problems in economics (Sims, 1980), Ives suggested their use for predicting responses of populations to environmental changes (Ives, 1995). His specific motivation was to establish techniques for studying how population abundances change in response to long-term environmental trends and for partitioning different factors driving changes in mean population densities in response to these trends. Since this early work, MAR models have been chosen to represent the interaction dynamics between biotic and abiotic drivers, infer the intra- and interspecific effects of species abundances on population growth rates, identify environmental drivers of community dynamics, predict the fate of communities submitted to environmental changes and extract measures of community stability and resilience. The latter was initially applied to lake and marine systems and later in terrestrial ecology (Certain, Barraquand, & Gårdmark, 2018).

Thus, two modeling frameworks with entirely different structures have been proposed for essentially the same purpose of extracting key features of dynamic interactions among coexisting populations of different species from observed time series data. Both methods have had successes, but a direct comparison of the two approaches has not been reported. Such a comparison is the subject of this article.

LV are ODE models, whereas MAR are statistical models. The former were designed to elucidate the long-term dynamics of interacting populations, whereas the latter were conceived to also describe the stochastic structure of a dataset. Our focus for their comparison is the ability of each model structure to produce an acceptable fit to the available data and to capture the process dynamics underlying the observed trends in population abundances.

We use four versions of MAR: MAR without any data transformation, MAR with log transformation, MAR upon data smoothing and MAR with log transformation upon data smoothing. Log transformation is necessary for comparing the general mathematical interpretation of a MAR model with a typical ecological interpretation, where they can be viewed as multispecies competition models with Gompertz density dependence (Ives, 1995; Certain et al., 2018) (Section 1.3 of the *Supplements*). Data smoothing is explored to assess if the advantages of LV models are in fact due to this preparatory step. It is clear that data smoothing will impede the ability of MARs to describe stochastic structures in the data, but this aspect is not the focus of this study.

We begin with a description and comparison of the main features of LV models and MAR models, subsequently analyze small synthetic systems, which offer the advantage of simplicity and full knowledge of all model features, and then assess several real-world systems. It is quite evident that it is impossible to compare distinct mathematical approaches with absolute objectivity and without bias (Rykiel, 1996), and it sometimes happens that inferior choices of models in specific cases outperform otherwise superior choices. We will attempt to counteract these vagaries by selecting case studies we consider representative and by stating positive and negative facts and features as objectively as possible.

## 2. Materials and Methods

### 2.1. Lotka-Volterra models

Lotka-Volterra (LV) models (Lotka, 1925; Volterra, 1926) are systems of first-order ordinary differential equations (ODEs) of the format

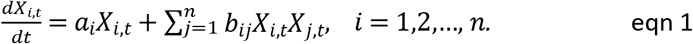

The left side of equation 1 represents the change in species *X*_*i*_ with respect to time. The equation with only the first term on the right side, *a*_*i*_*X*_*i*,*t*_, yields exponential growth, while the sum captures interactions between pairs of populations. Most of these terms represent interactions between different species, such as predation, competition for the same resources or cooperation, but one term in each equation, *b*_*ii*_*X*_*i*,*t*_*X*_*i*,*t*_, accounts for interactions among the members of the same species and is sometimes interpreted as “crowding effect.” Background and further details regarding these models are presented in *Supplements* Section 1.1. Because ODEs are natural representations of dynamic processes, the index *t* is usually omitted.

### 2.2. Estimation of LV Parameters Based on Slopes of Time Courses

Any of the numerous generic parameter estimation approaches for systems of nonlinear ODEs may be used to estimate the parameter values of LV systems; reviews include (Mendes & Kell, 1998; Wedelin & Gennemark, 2007; Chou & Voit, 2009). Here, we use a combination of smoothing, slope estimation, and parameter inference, for which we use the recently introduced Algebraic Lotka-Volterra Inference (ALVI) method (Voit *et al*., 2021). We begin by smoothing the raw time series data in order to reduce noise in the data as well as in their slopes, where the effects of noise are known be exacerbated (Knowles & Renka, 2014). Many options are available, but smoothing splines and local regression methods are particularly useful (Cleveland, 1981); they are reviewed in *Supplements* Section 1.2.1. Splines have degrees of freedom and we will refer to a spline with, say, 8 degrees of freedom as “8DF-spline”.

The estimation of slopes allows us to convert the inference problem from one involving ODEs into one exclusively using algebraic functions. This conversion is accomplished by substituting the left side of equation 1 with estimated slopes that correspond to values to the dependent variables on the right side, which leaves the parameters as the only unknowns (Voit & Savageau, 1982; Varah, 1982; Voit & Almeida, 2004; see also *Supplements* Sections 1.2.2. and 1.2.3.).

After the differentials are replaced with estimated slopes, two options permit the inference of the parameter values of LV-models. We can apply simple multivariate linear regression (ALVI-LR), where we either use all datapoints or iterate the regression several times with subsets of points, which is a natural approach of creating ensembles of solutions. As an alternative, if *n* is the number of variables, one may use *n*+1 of the datapoints and slopes, which results in a system of linear equations that can be solved with simple algebraic matrix inversion (ALVI-MI). For a thorough description of the ALVI method please see (Voit et. al., 2021) and an example in *Supplements* Section 1.2.4.

### 2.3. Multivariate Autoregressive (MAR) models

In contrast to the ODEs of the LV format, Multivariate Autoregressive (MAR) models are discrete recursive linear models. They have the general format

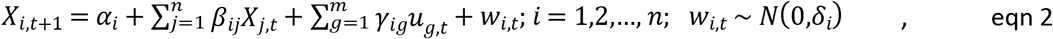

where *u_g_*_,t_ are environmental variables and *w_i_*_,t_ represents normally distributed noise. This set of equations, for different *i*, is usual represented in the matrix form

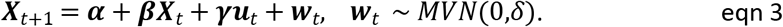

Explained in words, the “state” of the system at time *t*+1, expressed by the vector *_t_*_+1_, depends exclusively on the state of system one time unit earlier, ***X****_t_*, as well as on external inputs and stochastic environmental effects. Furthermore, *α* is the vector of intersects and ***β*** is one row of the population interaction matrix. The term ***γ****u*_*t*_ describes how cofactors affect the dependent variables. Specifically, ***u****_t_* is a vector of external variables and ***γ*** is the vector of weights associated with these external variables. Finally, the term *w*_*t*_ is a vector representing stochastic noise affecting the dependent variables. Background and further details regarding these models are presented in *Supplements* Section 1.3.

### 2.4. Parameter estimation for MAR

Software packages for the estimation of MAR model parameters greatly facilitate the use of these models. An example is the package MARSS, which uses an expectation maximization algorithm (Holmes, Ward, & Wills, 2012; Holmes, Ward, & Scheuerell, 2020). Some details of MARSS are discussed in in *Supplements* Section 1.4. and in the next section.

### 2.5. Structural Similarities between Modeling Formats

Although LV and MAR models have both been proposed for characterizing the interactions among populations within a mixed community, they are distinctly different in structure and appearance. Nonetheless, they also exhibit fundamental mathematical similarities, which are sketched below; a detailed analysis is presented in *Supplements* Section 1.5.

To assess these similarities, we focus on models without environmental factors, and thus on

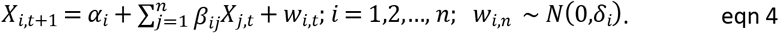

We also suppose that the MAR variables represent the logarithms of abundances, as proposed in (Dennis & Taper, 1994; Ives, 1995; Certain *et al*., 2018). Borrowing the principles of solving ODEs with Euler’s method, we discretize the LV model, which yields

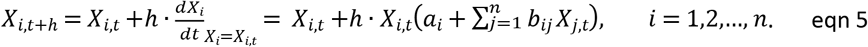

If the dynamics remains close to the steady state, then *X_i,t+1_* − *X_i,t_* ≈ 0 for any given *t*. Substituting this approximation into equations 4 and 5 yields

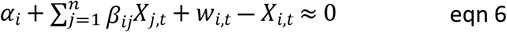

and

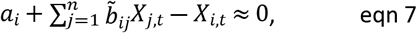

respectively. Ignoring the noise term in the MAR model, the two sets of near-steady-state equations 6 and 7 are the same if *α_i_ = a_i_*, *β_ij_ = b_ij_* for all *i ≠ j* and *β_ij_ = b_ij_* + 1 for *i* = *j*. Thus, the MAR and LV models are mathematically equivalent at the steady state and similar close to it.

## 3. Results

The comparison between LV and MAR models may be executed in two ways. A purely mathematical approach was sketched in Section 2.5. An alternative approach focuses on practical considerations and actual results of inferences from data. It is described in the following.

For simplicity, we omit environmental inputs (*c*_*i*_*X*_*i*_*U*_*t*_ and *γ*_*i*_*u*_*t*_, respectively) and begin by testing several synthetic datasets with different dynamics. We suppose that these data are moderately sparse and noisy, to mimic reality. In particular, we test whether the LV inference from synthetic LV data returns the correct interaction parameters and whether the MAR inference from synthetic MAR data does the same. Subsequently, we test to what degree LV inferences from MAR data yield reasonable results and *vice versa*. Finally, we apply the inference methods to real data. As the main metric, we compare the sums of squared errors and use a Wilcoxson rank test to assess the significance of the difference.

### 3.1. Case study 1: Synthetic LV data

The first case consists of data that were generated with a four-variable LV model and superimposed with synthetic, normally distributed noise (for details, see *Supplements* Section 2). We also generate a smaller noisy dataset, which however comes with replicates. The specific question we address is whether the LV and MAR inference methods return the true dynamics and parameter values.

The fits for the noisy and replicate LV datasets are presented in Figures 1 a, b, along with parameter estimates. These generally possess the correct sign and could, if deemed beneficial, serve as the starting point for an additional, refining optimization, for instance with a steepest-decent method. The inferred and true values are quite similar for the replicate dataset. By contrast, the parameter values inferred for the noisy dataset do not exactly recoup the true values. In fact, these inferences yield slightly better fits to the noisy data (SSE=17.981) than the “true” values (SSE=18.197), due to the noise. Because we usually obtain better results through ALVI-MI, we display those results here and ALVI-LR results in the *Supplements*.

**Figure 1:**
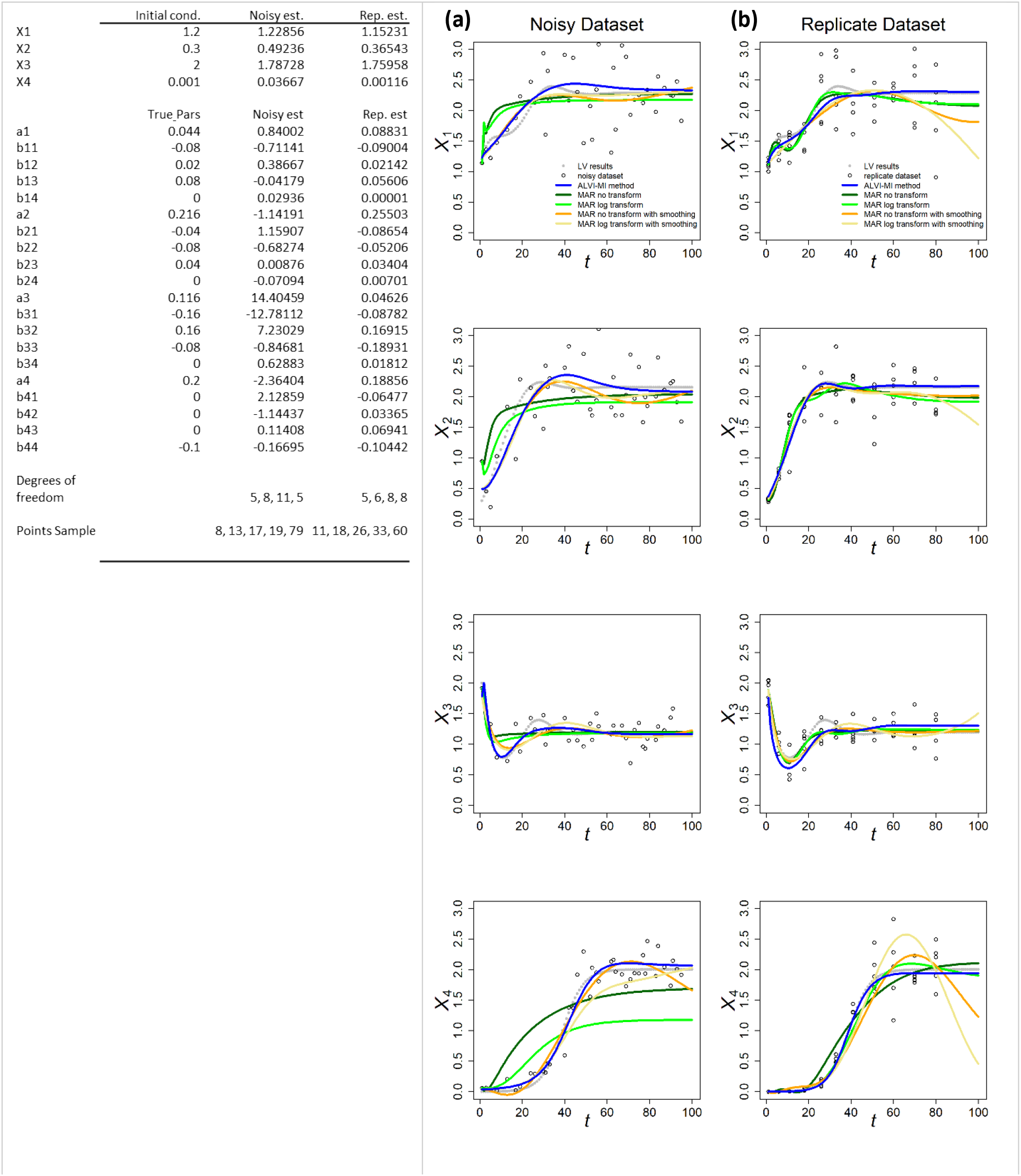
ALVI-MI and MARSS methods applied to noisy (a) and replicate (b) LV datasets. Original synthetic data are shown as gray dots and data with added noise as black circles. ALVI results are presented in blue. True parameters and ALVI-MI estimates are presented in the Table. MAR estimates are presented in green, orange and yellow. Data and parameter estimates for MAR can be seen in Table S1. SSEs for all fits are presented in Table 1 toward the end of the article.

**Table 1.** Sum of squared errors (SSE) of data fits for all experiments with ALVI-LR, ALVI-MI and four variants of the MAR methods. We also include SSEs for the estimates obtained by Mühlbauer *et al*. (2020) for LV data presented in Figure 3. Bold values identify the lowest SSE score for each example. Examples used in the Wilcoxson rank test are marked with asterisks.

|  | Shown in | ALVI-LR | ALVI-MI | MAR | MAR logTrans | MAR with smoothing | MAR with log smoothing | Mühlbauer *et al.* | Test |
| --- | --- | --- | --- | --- | --- | --- | --- | --- | --- |
| Noisy LV dataset | Fig. 1a | 9.009 | **4.362** | 37.31 | 53.31 | 7.364 | 8.260 |  | * |
| Replicate LV dataset | Fig. 1b | 11.41 | **2.953** | 6.247 | 5.439 | 13.87 | 42.83 |  | * |
| Noisy MAR dataset | Fig. 2a | 2,727 | 1,142 | **908.3** | 37,447 | 1,191 | 1,604 |  | * |
| Replicate MAR dataset | Fig. 2b | 2,518 | 1,462 | 933.1 | **177.967** | 1,773 | 1,482 |  | * |
| SynthData1 | Fig. S5a | **3.29E-07** | 9.04E-07 | 10.49 | 4.626 |  |  |  |  |
| SynthData2 | Fig. S5b | 8.68E-06 | **8.57E-06** | 3.029 | 5.876 |  |  |  |  |
| SynthData3 | Fig. S5c | **0.002** | 0.018 | 5.111 | 4.378 |  |  |  |  |
| SynthData4 | Fig. S5d | **5.65E-04** | 2.10E-02 | 2.47E+04 | 3.99E+35 |  |  |  |  |
| SynthData5 | Fig. S5e | **7.048** | 18.94 | 16.59 | 15.70 |  |  |  |  |
| SynthData6 | Fig. S5f | 71.27 | **28.36** | 7,379,825 | 174.2 |  |  |  |  |
| Mühlbauer et al. 1 | Fig. 3a | 2,588 | **2,516** | 18,160 | 51,738 | 17,877 | 16,399 | 3271 | * |
| Mühlbauer et al. 2 | Fig. 3b | 6,204 | **4,781** | 13,016 | 22,162 | 9,293 | 12,447 | 40604 | * |
| Mühlbauer et al. 3 | Fig. 3c | 24,183 | **2,191** | 4,619 | 10,146 | 4,780 | 8,795 | 578.1 | * |
| Mühlbauer et al. 4 | Fig. 3d | 17,700,169 | **1,118,350** | 2,199,174 | 17,77,559 | 9,904,281 | 3,973,876 | 7934136 | * |
| Mühlbauer et al. 5 | Fig. 3e | 12,114,811 | **2,251,804** | 4,245,588 | 4,450,871 | 5,596,187 | 4,554,089 | 2893764 | * |
| Holmes et al. 1 | Fig. 4a | 168,134,153 | 153,703,251 | 217,478,452 | 166,186,993 | **143,728,190** | 212,424,910 |  | * |
| Holmes et al. 2 | Fig. 4b | 32,326,913 | **5,556,585** | 10,267,681 | 10,947,294 | 11,136,306 | 14,342,449 |  | * |

Figure 1 also displays the MARSS estimates with and without log-transformation of the data and with or without data smoothing. With respect to *X*_1_, *X*_2_ and *X*_3_, these estimates are of adequate quality. They present good fits, although not as good as the LV inference, which is probably not surprising as the data were generated with an LV system. MARSS did not perform well for the “detached” variable *X*_4_, especially for the noisy dataset.

Because MARSS yields parameter values for a discrete recursive system, they are not directly comparable to the true parameters of a LV system; nonetheless, their numerical values are recorded for completeness in Tables S1.4 and S1.5.

For MARSS inferences from the replicate dataset we had to average points with the same time value. MARSS did not converge for all parameters but it still presented a relatively good fit. Additional details are presented in *Supplements* Section 2.

The ALVI-MI method also works well for more complicated dynamics, as demonstrated in *Supplements* Section 2 and Figure S5.

### 3.2. Case study 2: Synthetic MAR data

Here we reverse the set-up of Case Study 1 by creating synthetic data with an MAR model and test whether either method can infer results corresponding to the original system.

As a representative example, we use a four-variable MAR system to create 31 synthetic datapoints. We create a *noisy MAR* dataset by adding noise in the form of a random normal variable of mean zero and a standard deviation of 20% of each variable’s mean to the original 31 points sample. We also create a *replicate MAR* dataset by choosing 12 points and generating 5 replicates by multiplying their values by a random normal of mean 1 and standard deviation of 0.2. The initial conditions, parameters, dynamics, ALVI and MAR fits are presented in Figure 2. All fits to the synthetic MAR data are satisfactory. The LV parameters are not directly comparable to the MAR parameters.

**Figure 2:**
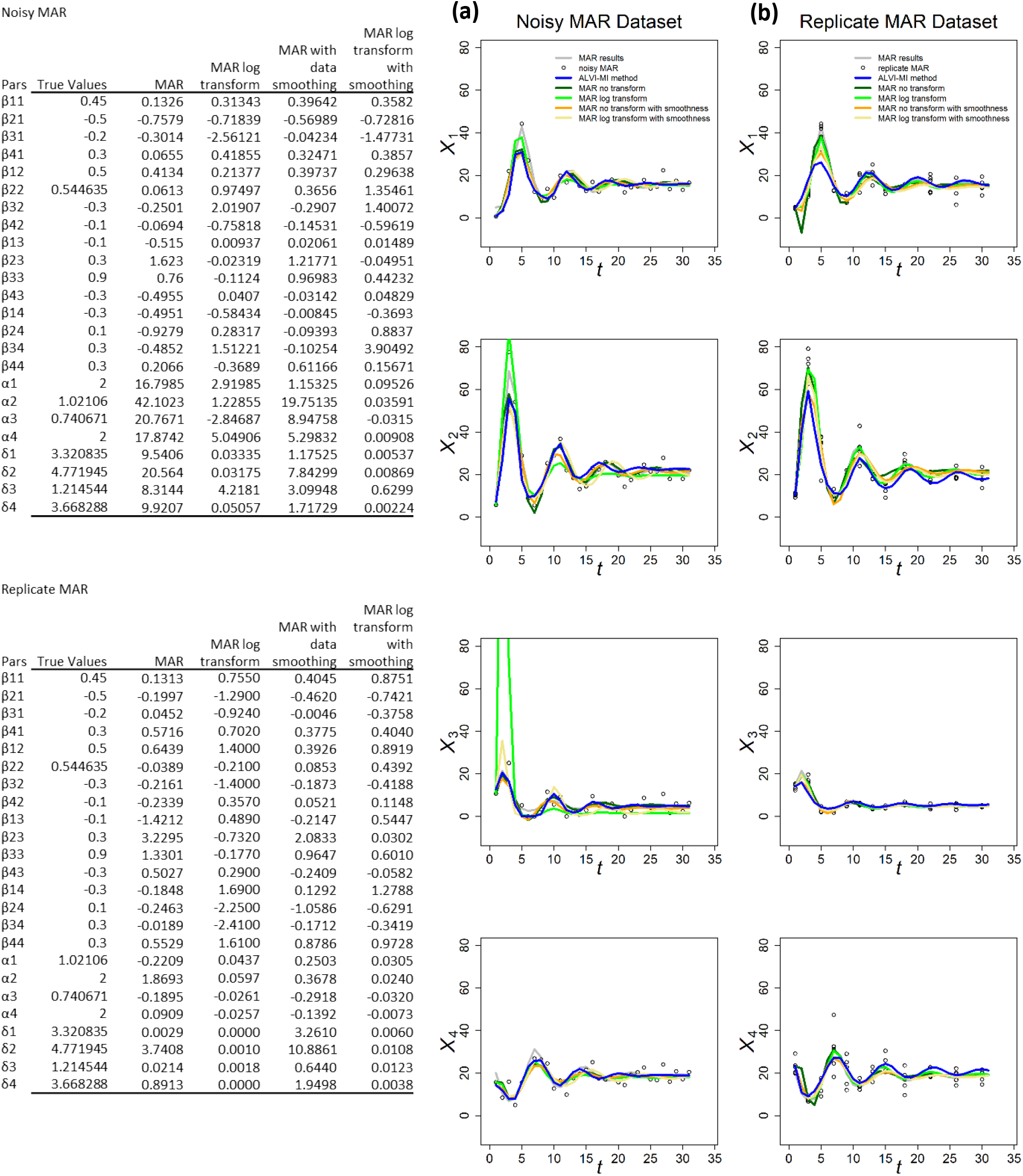
MARSS and ALVI-MI methods applied to noisy (a) and replicate (b) MAR datasets. Original synthetic data are shown as gray dots, data with added noise as black circles. MAR estimates are presented in green, orange and yellow. ALVI results are in blue. The variables of the noisy dataset were smoothed with 15DF-splines and the ALVI-MI solution was calculated with spline points at times 2, 6, 15, 18 and 26. The variables of the replicate dataset were smoothed with 15DF-splines and the ALVI-MI solution was calculated with spline points corresponding to times 1, 3, 11, 13 and 15. Data and parameter estimates for ALVI can be seen in *Supplements* Table S5. SSEs for these fits are presented in Table 1 toward the end of the article.

In most cases, the different MAR models had difficulties retrieving the true parameters of the system, and sometimes even the correct sign (Figure 2). This is probably due to the small number of datapoints: Certain *et al*. (Certain et al., 2018) suggest that the length of the time series should be at least 5 times greater than the number of *a priori* nonzero elements in the matrix B in order to recover interaction signs correctly. Our sample has 31 observations and should have at least 80. For *X*_1_ and *X*_4_, all models show a similar fit, but not for *X*_2_ and *X*_3_, where MAR models with log transformation show a considerable deviation from the data.

### 3.3. Case study 3: Experimental data from the literature, previously used for inferences with LV and MAR models

#### 3.3.1. Published LV inferences

Data from Georgy Gause’s 1930s experiments and others were recently compiled in the R package gauseR (Mühlbauer *et al*. 2020). In the accompanying paper, the authors present five examples to test their method for estimating LV model parameters. We use the exact same examples to demonstrate to what degree LV and MAR methods are compatible with these real-world data and compare our results to those presented by Mühlbauer and colleagues. For more information regarding the original experimental data, see (Gause, 1934), (Huffaker, Shea, & Herman, 1963), (McLaren & Peterson, 1994) and (Mühlbauer et al., 2020). The results are presented in Figure 3, with data as symbols and various estimates as lines. SSEs of the different estimates for these and other test examples are presented in Table 1.

**Figure 3:**
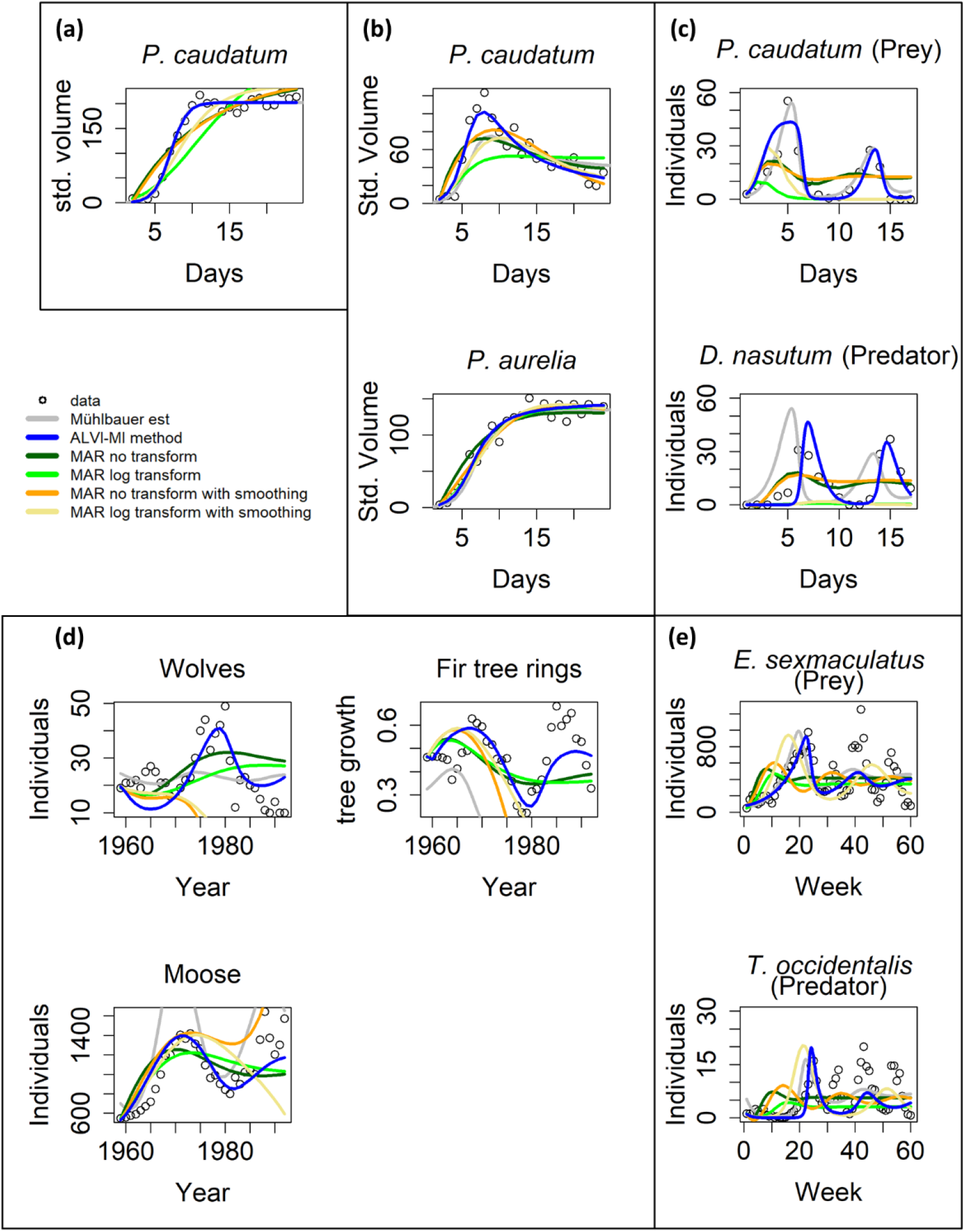
Model inferences associated with Gause’s data (Gause, 1934). Circles show observations, gray lines are the estimates from Mühlbauer *et al*. (Mühlbauer et al., 2020). ALVI-MI method estimates are presented as blue lines and MAR estimates as green, orange and yellow lines. See text and *Supplements* Tables S4.1 and S4.3 for further details.

The MAR method never outperforms the other methods considered. To be fair, these examples had been used to test actual data for compatibility with the LV structure, which may explain the superior performance of the LV model. Nonetheless, these are the types of data the MAR method is supposed to capture.

For the case in Figure 3a, ALVI-MI yields the same results as found in (Mühlbauer et al., 2020). In contrast, the MAR estimates are poor, with a very high estimate for the noise (Table S4.3), especially if one does not use log-abundances; this problem occurs in all cases presented in Figure 3. The data in Figure 3a are close to a logistic function, similar to *X*_4_ in the previous noisy dataset, where MAR also did not perform well.

Figure 3b shows data from a competition experiment between *Paramecium caudatum* and *Paramecium aurelia* that were co-cultured. Estimates for *P. aurelia* are similar for all methods but ALVI-MI exhibits clear superiority for *P. caudatum*.

The data in Figure 3c are complicated. Mühlbauer and colleagues noted that aditional quantities of bacteria were introduced to avoid species extinction. Furthermore, many datapoints in this dataset are zero, which causes problems for the parameter estimators. As a remedy, we changed the zeros to 10^-5^, but our initial estimates still produced poor fits. However, if we use the estimated trajectories from Mühlbauer *et al*. as “data,” quasi as a diagnostic measure, ALVI-MI captures the parameters that reproduce the fit of Mühlbauer *et al*.. This finding suggests that the initial poor fit is not a problem of LV adequacy. Instead, we hypothesize that the problem was caused by insufficient datapoints or almost-linear dependence, which affects the matrix inversion. To test this hypothesis, we used the first splines as data to create a second set of splines that has more datapoints to create the subsample to be used on ALVI-MI. We were able to achieve the presented fit, which is still somewhat inferior to the one by Mühlbauer *et al.,* but a considerable improvement over our initial fits.

When calculating splines for this dataset, it is difficult to choose degrees of freedom that capture both maxima. High degrees of freedom capture the global maxima but overshoot the local maxima. Low degrees of freedom capture the local but undershoot the global maxima. We suspect this to be the cause of ALVI’s initial poor performance. Still, ALVI yields better fits than MAR.

The data in Figure 3d are also complicated, in this case due to two aspects. First, they show a stark difference in absolute numbers, with the abundance values for moose being several magnitudes higher than the numbers of tree rings. As a potential remedy, we normalized the fitting error for each dependent variable by dividing it by its mean to balance the SSE. The result is shown in Figure 3d. The LV models perform better than MAR, and MAR with log-abundances produces a better noise estimate than with the untransformed data.

The second issue is the fact that, around 1980, the wolves were exposed to a disease introduced by dogs that caused a precipitous drop in the population between 1981 and 1982 (Park Service, 2021). Typical mathematical models are not equipped to simulate such a black swan event, and the totality of results from the various methods suggests that neither LV nor MAR may be good models for this system, because none of the fits, by Mühlbauer *et al*., ALVI, and MAR, are entirely satisfactory. Nonetheless, our ALVI results present a decent fit for moose and fir tree rings. To improve the fit to the wolf data, we divided the data into two groups, from 1959 to 1980 and from 1983 to the end of the series and estimated parameters for the two intervals. The results are presented in Figure S7 in orange lines. The fit is greatly improved, although still not perfect.

Figure 3e describes yet another complicated example. According to the inference, the ALVI-MI estimates fit the first peak well but the oscillations die down, in contrast to the data. Estimates from Mühlbauer *et al*. produce even poorer estimates, suggesting that the data may not be compliant with the LV structure. As in the previous example, MAR models do not capture the dynamics, although MAR with log-abundances produces good noise estimates. Surprisingly, MAR with smoothing presented very poor fits to these data.

We repeated the analysis using ALVI-LR instead of ALVI-MI. The results were by and large similar and slightly inferior; they are shown in *Supplements* Figure S6 and Table S4.2.

One should note that Mühlbauer *et al*. used a steepest-descent method, while our method did not. Therefore, our results could possibly be further improved by adding a refinement cycle of steepest-descent optimization. The main problem of these algorithms, getting stuck in local minima, would presumably not be an issue, since the ALVI results are already close to the optimum.

#### 3.3.2. Published MAR inferences

Here we use two datasets presented in the MAR inference package MARSS. The first dataset, “gray whales,” consists of 24 annual abundance estimates of eastern North Pacific gray whales during recovery from intensive commercial whaling prior to 1900 (Gerber, Demaster, & Kareiva, 1999). It is thus to be expected that the whales are initially far from the carrying capacity of the system. The second case consists of data for wolf and moose populations on Isle Royale in Lake Superior between 1960 to 2011; this dataset was used by Holmes and colleagues (Holmes et al., 2020) to demonstrate usage of the MARSS R package.

Figure 4a show fits to the gray whale data (Gerber *et al*., 1999). ALVI-MI noticeably outperforms MAR, even though the data came from a MAR demonstration. In particular, the MAR results suggest that the whales are close to regaining their carrying capacity, which seems to contradict the trend in the data. The SSEs can be seen in Table 1. It is unclear why the MAR method without transformation does not perform better. As it stands, the estimates are inadequate (with the highest SSE) and have a very high variance for the error. An LV model with one variable is a logistic function, and the LV fit represents initial quasi-exponential growth that starts to slow down after a while. This behavior nicely reflects the fact that the whales were recovering from very small numbers due to overfishing but are apparently still much below the carrying capacity.

**Figure 4:**
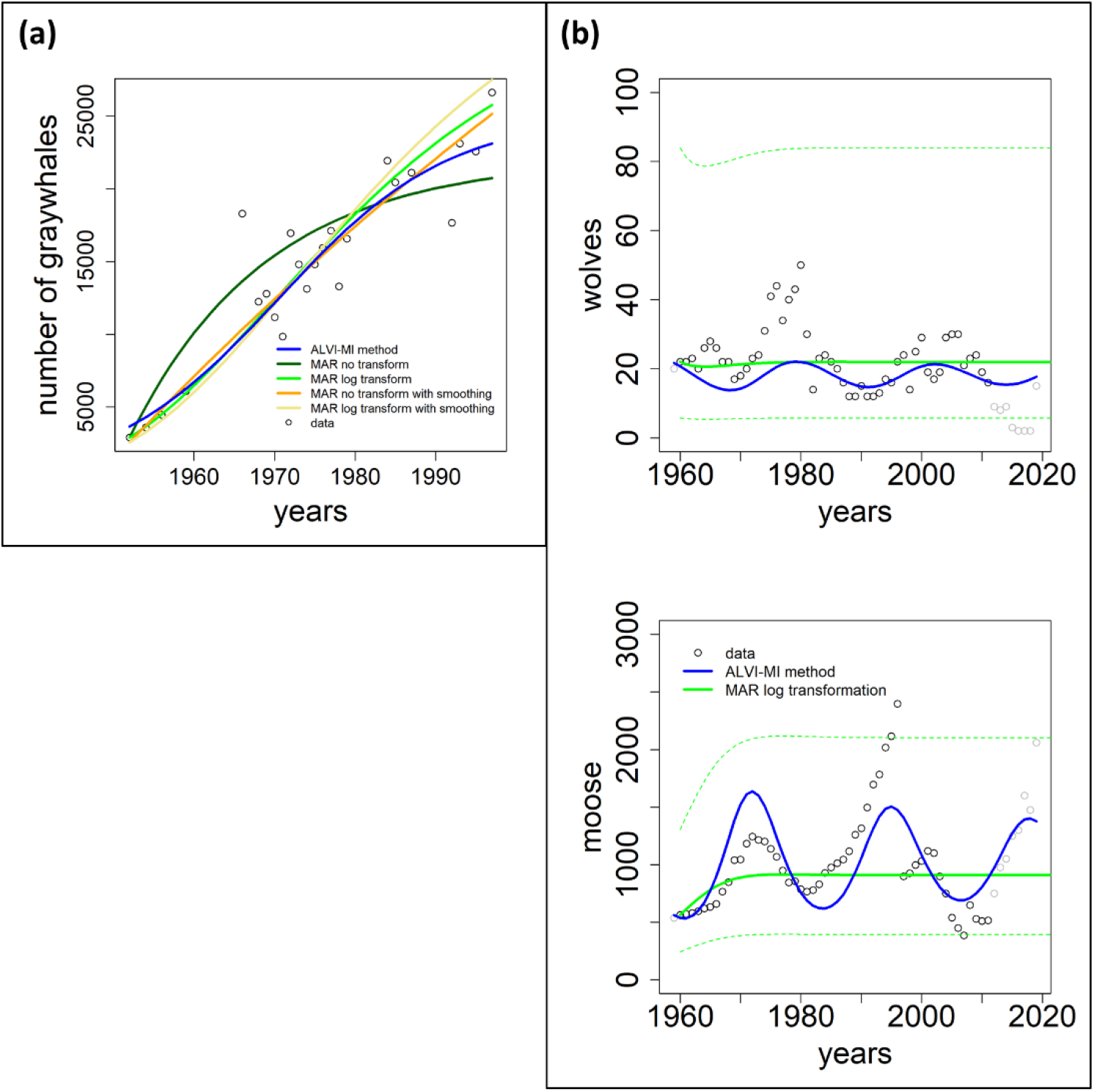
Two datasets of wildlife observations. Column a: Abundance data of gray whales (Gerber et al., 1999). The plot shows results from ALVI-MI in blue; MAR estimates are displayed with green, orange and yellow lines. Column b: Wolves and moose on Isle Royale (Vucetich, 2021). The original data used for parameter estimation are displayed with black circles, data not used by the estimation processes are shown in gray, ALVI-MI results are in blue, MAR estimates using log-abundances are displayed with green lines. The dashed lines indicate confidence intervals for the MAR estimates. Values of the estimates can be seen in Table S6.

Figure 4b returns to the Isle Royale dataset from Vucetich (Vucetich, 2021), which we already used in the context of examples from the collection of Mühlbauer and colleagues (Mühlbauer et al., 2020); *cf*. Figure 3. Holmes *et al*. (Holmes et al., 2020) used only the data of wolves and moose for a MAR analysis but extended them over a longer time horizon. Specifically, eight datapoints were added since the former usage of this dataset by Holmes and colleagues, from 2012 to 2020 (gray symbols in Figure 4b).

The results of the MAR model are identical with those published in (Holmes et al., 2020), with the same log transformation and z-scoring of the data, and the same parameter values were inferred. The result consists of acceptable estimates, although we found a slightly better fit without the z-scoring. Still, for a direct comparison, we opted to present the example exactly as Holmes *et al*. did. Interestingly, these fits miss all oscillatory behavior seen in the data. The ALVI results do show oscillations but clearly suffer from the disruption in the wolf population in 1981 and 1982, as discussed before. Because we used in this example only MAR with log transformation, we display the confidence intervals for the MAR model as dashed green lines. Very few datapoints are outside the confidence intervals.

### 3.4. Comparative summary of the performance of LV and MAR in the presented examples

Inspection of Table 1 renders is evident that ALVI clearly performs better than MAR. In a few cases, the ALVI-LR solution gives a better SSE than ALVI-MI, but the difference between the two is not substantial. ALVI-LR appears to be superior when the data are noise-free.

We used a one-sided Wilcoxson rank test to see if the differences in performance are significant. The results and alternative hypotheses for these tests are presented in Table 2.

**Table 2.** Test results of one-sided Wilcoxson rank.

| Null hypothesis $H_0$: | Alternative hypothesis $H_1$: | $p$-value |
| --- | --- | --- |
| SSE values of ALVI-MI are equal or higher than corresponding MAR values | SSE values of ALVI-MI are less than corresponding MAR values | **0.0093** |
| SSE values of ALVI-MI are equal or higher than corresponding MAR values for log transformed variables | SSE values of ALVI-MI are less than corresponding MAR values for log transformed variables | **0.0024** |
| SSE values of ALVI-MI are equal or higher than corresponding MAR values for smoothed data | SSE values of ALVI-MI are less than corresponding MAR values for smoothed data | **0.0269** |
| SSE values of ALVI-MI are equal or higher than corresponding MAR values for log transformed variables and smoothed data | SSE values of ALVI-MI are less than corresponding MAR values for log transformed variables and smoothed data | **0.0005** |
| SSE values of the ALVI-MI are equal or higher than SSE values obtained with the ALVI-LR | SSE values of ALVI-MI are less than SSE values obtained with the ALVI-LR | **0.0005** |
| SSE values of MAR with data smoothing are equal or higher than MAR values without | SSE values of MAR with data smoothing are less than MAR values without | 0.7676 |
| SSE values of MAR with log transformation and data smoothing are equal or higher than those for MAR with log transformation values | SSE values of MAR with log transformation and data smoothing are less than those for MAR with log transformation values | 0.7935 |

The data supports that the ALVI-MI method produces smaller SSE’s than the other methods considered. Also, in the last two tests, the data do not show evidence that data smoothing reduces the SSE’s in MAR.

Comparing the results ALVI-LR and ALVI-MI with respect to the absolute value of the difference between true and estimated parameters, we obtained mixed results (Table 3). Indeed, a one-sided Wilcoxson rank test with the alternative hypothesis that the absolute errors in parameter values associated with ALVI-LR were smaller than those associated with ALVI-MI did not yield a significant *p*-value (0.2783).

**Table 3.** Absolute differences between true and estimated parameters for ALVI-LR and ALVI-MI. Bold font indicates the lower difference in each case. A statistical test did not reject the null hypothesis of no difference between the results of the two methods.

|  | ALVI-LR | ALVI-MI |
| --- | --- | --- |
| Noisy Data 5% | **1.97538** | 2.68019 |
| Replicate Data 5% | **0.24117** | 0.32537 |
| Noisy Data 20% | **1.35692** | 46.60082 |
| Replicate Data 20% | 0.8835 | **0.66838** |
| Synthetic Data 1 | **0.00021** | 0.00038 |
| Synthetic Data 2 | 0.20832 | **0.15534** |
| Synthetic Data 3 | 1.06975 | **0.0982** |
| Synthetic Data 4 | **0.19183** | 0.60795 |
| Synthetic Data 5 | 0.82403 | **0.36212** |
| Synthetic Data 6 | **0.18794** | 0.97794 |

Comparing the values in Tables 1 and 3 for noisy and replicate datasets, the latter presented smaller values. However, using the one-sided Wilcoxson rank test for the values in Table 1 with the alternative hypothesis—that the SSE values for the replicate dataset were smaller than those for the noisy datasets— did not yield a significant *p*-value (0.2734). For the values in Table 3, the test produced a *p*-value (0.0625) smaller than 0.1, suggesting that time series data are better than clustered data for parameter estimation in LV using ALVI.

## 4. Discussion

We have compared LV and MAR models using, as parameter estimation strategies, a recently introduced ALVI method for LV models (Voit et al., 2021) and the established MARSS method for MAR models (Holmes et al., 2012). Summary Table 1 renders it evident that ALVI usually outperforms MARSS, with often substantially lower SSE values, and that LV models in the vast majority of cases provide better representations than MAR models. Furthermore, Certain *et al*. (Certain et al., 2018) prescribed the length of the data series as at least 5 times greater than the number of nonzero interaction elements in order to recover interaction signs correctly. A caveat is that we presented the MAR model equations without noise, thereby ignoring an important part of these models that characterizes the stochastic structure of the data.

ODEs and MAR models are derived from different philosophies and comparing them fairly is not straightforward. Here we compared the two approaches from a point of view of someone who is more interested in capturing the dynamics of a phenomenon, which poses an immediate disadvantage for MAR models, because they were created for phenomena with random noise and for characterizing the structure of this noise. Consequently, we found much of the dynamics quantified as noise in the estimates. Thus, an overarching conclusion is that the two approaches are different tools that are adequate in different situations. MAR models are well suited for simulations with variables presenting low abundances and investigations where the quantification of noise is relevant. By contrast, LV models, and ODE models in general, are better suited to capture the dynamics of a phenomenon and for cases where the dependent variables have high abundances and a high signal-to-noise ratio. Nonetheless, if the characterization of noise is of interest, one might subtract the LV fit from the raw data and then assessing the remainder. Another advantage of LV models is that ALVI permits *a priori* tests for the adequacy of the LV structure for a given dataset (Voit et al., 2021).

Because the LV structure is continuous, we can evaluate it at any point or choose any interval between the points in the numerical solution. By contrast, MAR does not truly reveal a time resolution higher than its intrinsic interval between solution points. However, Holmes *et al*. (Holmes et al., 2012) demonstrated with the MARSS R function that it is feasible to interpolate any number of missing values between the known datapoints, and that this method can be used to decrease the time unit for stepping forward. While this step does not make the MAR model as densely time-resolved as an ODE model, it mitigates the apparent granularity disadvantage considerably.

ALVI allows a choice between linear regression and matrix inversion. The former is simpler, because it uses all points available, and faster due to the fact that no data sample needs to be chosen. In most cases tested, it also produces good fits and estimates. However, the latter usually produces slightly better results and works well even in cases where the ALVI-LR solution fails (Table 1). It also offers a natural approach to inferring whole ensembles of well-fitting model parameterizations.

Results obtained with MARSS or with ALVI-LR are rather robust if the data are noisy, whereas solutions with ALVI-MI may be sensitive to small alterations in the data. As an example, consider the synthetic MAR data presented in Figure 2, where we added noise as a normally distributed variable with mean zero and standard deviation of 0.2 of the dependent variable mean. For comparison, consider now an alternative sample (Figure S8) obtained by applying the same procedure, but with a different seed for noise creation. The alternative dataset is almost indistinguishable from the original dataset in Figure 2, but using the same point sample determined for the original dataset, ALVI-MI produces different parameter values. This means that if we calculate a new set of splines, we should also search for a new point sample. The conclusion is that, although the inferred fits are similarly good, the parameter values associated with the best fit for a noisy point sample may not be optimal for another noisy sample, which may not be surprising. In fact, we showed in a different example with noise that the inferred parameter values yielded a better SSE than even the true values (Section 3.1). The argument may also be turned around into a positive feature: Different noisy datasets or subsamples of these datasets can easily be used to create natural ensembles of models that characterize the underlying data in a robust manner and yield additional insights into the variability of the model parameters.

The MARSS software makes modeling with MAR models easy, although not entirely automatic, as many options must be tested to find the one that returns the best fit in each case. For example, one must decide whether to use estimated initial conditions or the initial datapoints and which variables should have the same noise level. By contrast, the ALVI method for LV models is novel (Voit et al., 2021), and while all steps are straightforward and code is available on GitHub [https://github.com/LBSA-VoitLab/Comparison-Between-LV-and-MAR-Models-of-Ecological-Interaction-Systems], no formally published software currently exists that encompasses all these steps in a streamlined manner. As a new tool, ALVI offers several avenues for further refinement. One important component is optimal data smoothing with splines, which requires the determination of a suitable number of degrees of freedom and may also employ weights for different variables within a dataset.

A comparison of MARSS results with or without log transformation of the dependent variable abundances did not yield clear results. If the MAR models are to be viewed as a multispecies competition models with Gompertz density dependence (Certain et al., 2018), the log transformation is required (*Supplements* Section 1.3). While inferences for LV models usually benefit from smoothing, the same is not true for MAR model, where smoothing in some cases, but certainly not always, led to improved data fits (Table 2).

MARSS uses a steepest decent optimization step, which ALVI presently does not. Although ALVI already performs better than MARSS (Table 1), it might be possible to improve its results even further by adding a refinement step based on steepest-descent optimization. Steepest-descent methods tend to get trapped in local minima if the initial guesses are poor, but ALVI would not likely encounter this problem, as the solutions are already very good and could be used directly as initial guesses for the refinement step.

Estimation and inference methods typically do not scale well. ALVI bucks this trend, at least to some degree, as both the smoothing and estimation of slopes occur one equation at a time. Thus, instead of scaling quadratically, the inference problem scales linearly. The ultimate matrix inversion or linear regression is essentially the same for all realistically sized models. Thus, the only time-consuming step within ALVI-MI is the choice of datapoints. An exhaustive test for all combinations grows quickly in complexity, but it is always possible to opt for a random search. The resulting solution is not necessarily the best possible, but can still provide an excellent fit or at least a valuable starting point for a steepest-decent refinement optimization. Importantly, many random solutions can also be collated to establish an ensemble of well-fitting models, which in most cases yields more insight than a single optimized solution.

## 5. Acknowledgements

This work was supported in part by the following grant: NIH-2P30ES019776-05 (PI: Carmen Marsit). The funding agency is not responsible for the content of this article.

## 6. Author Contributions

D.V.O., J.D.D., and E.O.V. conceived this project, performed the literature review, created the synthetic data examples and wrote of the manuscript. D.V.O. produced the code required for the study. All authors reviewed and edited the manuscript.

## 7. Data Accessibility

The R code for running the experiments is available on GitHub: https://github.com/LBSA-VoitLab/Comparison-Between-LV-and-MAR-Models-of-Ecological-Interaction-Systems

## 8. Additional Information

Competing financial interests

The authors declare no competing interests.

## SUPPLEMENTS

### 1. Materials and Methods

#### 1.1. Lotka-Volterra models

For a single variable, Lotka-Volterra (LV) models (Eq. [1] in the main text) reduce to the well-known logistic growth law

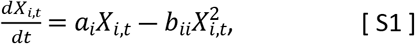

where the ratio *a*_*i*_/*b*_*ii*_ is called the “carrying capacity” of the system, which corresponds to the non-trivial steady state. If time-dependent environmental inputs are to be considered, one may add terms *γ*_*i*_*X*_*i*_*U*_*t*_, where *U*_*t*_ is an element of a vector of these inputs and the coefficients *γ*_*i*_ are weights that quantify the effects of the factors on species *X_i_* (Stein et al., 2013; Dam *et al*., 2016; Dam *et al*., 2020). The index *t* is usually omitted, and the left-hand side is often written as *X*_*i*_.

The LV system is a *canonical* model in the sense that its mathematical structure is immutable and scalable to any dimension (Voit, 2000). Such a canonical model may serve as a template to construct models of different systems that reasonably satisfy the following assumptions:

- encounters between and within species are representable by mass action kinetics;
- the environment does not change during the process, unless environmental variables are explicitly formulated as described above;
- the parameter values do not change during an experiment;
- the species respond to one another instantaneously;
- for very small population sizes, interactions are negligible and the rate of change (growth) of each population is initially proportional to its size, resulting in initial exponential growth;
- adaptations of species are absent or negligible.

Although the model structure and these assumptions might appear to be unduly rigid, LV models are extremely rich in the repertoire of their possible responses. In fact, the LV structure was shown to be capable of modeling any type of differentiable nonlinearities, including different kinds of oscillations and chaos (Vano et al., 2006), if sufficiently many auxiliary variables are permitted, some of which have mathematical, but no real biological meaning (Voit & Savageau, 1986; Peschel & Mende, 1986; Savageau & Voit, 1987). At the same time, the LV structure has severe limitations. For example, it is not well suited for metabolic pathway systems, because a simple conversion of a substrate *X*_1_ into a product *X*_2_ would require *X*_2_ to appear in its own synthesis term, although the generation of *X*_2_ in truth depends only on *X*_1_ and possibly some modulators (see (Voit, 2013) for this and other limitations).

LV models were initially used to describe the dynamics of predator and prey populations or of populations that compete for the same resources, but the same equations have also been used in entirely different contexts and fields, including physics (Nambu, 1986; Hacinliyan, Kusbeyzi, & Aybar, 2010), pollution assessment (Haas, 1981), economy (Zhou & Chen, 2006; Gandolfo, 2008), manufacturing (Chiang, 2012), and sales (Hung, Chiu, Huang, & Wu, 2017).

Beyond the fact that LV models can be formulated very easily, another significant advantage over other systems of nonlinear ODEs is the fact that LV models can be parameterized with linear regression methods if time series data are available (Voit & Chou, 2010). As an intriguing alternative, the linearity also permits us to select variable and slope values at *n*+1 time points and to obtain parameter inferences by solving a set of linear algebraic equations (see below). It is furthermore possible to estimate parameter values from sufficiently many profiles of species that initially coexist and ultimately survive under comparable conditions (Voit et al., 2021).

#### 1.2. Estimation of LV Parameters Based on Slopes of Time Courses

This section explains in detail an approach to parameter estimation that uses the Algebraic Lotka-Volterra Inference (ALVI) method. For a detailed explanation of the ALVI method itself please see (Voit et al., 2021).

##### 1.2.1. Smoothing

Even though one might consider smoothing a conceptually separate issue from the actual parameter inference, the two are so closely intertwined in our analysis that it appears useful to discuss smoothing options. The goal is two-fold. First, it is beneficial to reduce or even remove noise from the raw data, and second, this smoothing greatly aids the determination of slopes of the experimental time courses (see later).

A smoothed representation of a dataset implicitly integrates information that is not in the data. This implicit integration step is not entirely unbiased and requires prudent judgment, because it must answer the following questions, often without true knowledge of the system: Are the deviations between the data and the smoothing function due to (stochastic) noise or are they part of the true signal? For instance, are they the trace of true oscillations? Also, if very few data points deviate much more than all others from the smoothing function, are they true peaks or valleys or are they statistical outliers? It is difficult to answer these questions objectively, but two features of the data are of great benefit: First, if the variation in noise amplitude is much smaller than the range of signal values (high signal-to-noise-ratio), the distinction between signal and noise is relatively straightforward. Second, if the data come in replicates, they may support or refute the potential of true oscillations or peaks at certain time points in the data. Even if only one dataset is available, the biologist familiar with the phenomenon at hand usually has developed an expectation regarding signal and noise, and if there is no biological rationale for expecting oscillations or strong deviations from some simple trend, the smoothing strategies are flexible enough to allow the integration of the biologist’s knowledge and expectations. The result of the smoothing process therefore is a synthesis of all relevant information, constrained by external knowledge and reasonable expectations. Of course, it is also feasible to create alternative models with different thresholds between signal and noise and to analyze them side by side.

Independent of the options and intricacies of obtaining smoothed time courses of all variables, it is well known that the process of estimating slopes from data is more strongly affected by noise than the data themselves (Knowles & Renka, 2014). Expressed differently, if the noise is left unchecked, its effect on the estimated values of the slopes tends be higher than its effect on the values of the variables.

We explored a number of methods for smoothing the time course data and keeping the noise in check (Eilers, 2003; Vilela et al., 2007; Batista Júnior & Pires, 2014), cognizant of the fact that empirical raw data alone do not provide enough information of what is noise and what is relevant signal in the dynamics of the phenomenon under study.

One of the simplest approaches is the *three-point method*, where the slope at time point *t_k_* is taken as the average of the slopes at time points *t_k_*_-1_ and *t_k_*_+1_ (Burden, Faires, & Burden, 1993; Voit & Almeida, 2003). More sophisticated methods were reviewed in (Cleveland & Grosse, 1991; Eilers & Marx, 1996; Batista Júnior & Pires, 2014). For long, dense time series, moving average and collocation methods with or without roughness penalty (Ramsay et al., 2007) are often very effective. However, they tend to be unsuited for biological time series data because the measurements are usually quite sparse and obtained over a relatively short time horizon.

Smoothing splines and *local regression* methods like LOESS (locally estimated scatterplot smoothing) and LOWESS (locally weighted scatterplot smoothing) turned out to be particularly useful. A detailed description of these methods can be found in (Cleveland, 1981).

In a nutshell, splines are piecewise polynomial functions that: pass through all sample points, are continuous and have first and second derivatives that are continuous at junction points between adjacent intervals. In a smoothing spline, the first condition is substituted by a least-squares fit that is balanced with an additional criterion that penalizes splines with high second derivative values, which indicate local roughness (Cleveland, 1979; Garcia, 2010; Loader, 2012).

LOESS and LOWESS algorithms use locally-weighted polynomial regression. LOWESS is used for univariate smoothing and consists of computing a series of local linear regressions, with each local regression restricted to a window of x-values. Smoothness is achieved by using overlapping windows and by gradually down-weighing points in each regression according to their distance from the anchor point of the window. LOESS is for fitting a smooth surface to multivariate data and it is a generalization of LOWESS in that locally weighted univariate regressions are simply replaced by locally weighted multiple regressions. While LOESS is more versatile, LOWESS is faster and sometimes succeeds when LOESS fails (Cleveland, 1979; Cleveland & Devlin, 1988; Smyth, 2020). Locally-weighted polynomial regression methods have ‘span’ and splines have ‘degrees of freedom,’ which are parameters that control the degree of smoothing.

The main result of smoothing with splines is a reduction or even removal of what is believed to be noise in the data. The slope at each point can be computed directly from the smoothing spline, which after all is an explicit function. This step of slope determination offers two options: it allows us to estimate slopes only for the measured data points or to sample the smoothing function for any number of other points, which yields a larger set of numerical values for variables and slopes (Voit & Almeida, 2004).

If we select many points from the smoothing spline, we overcome the problem of data scarcity that is inherent in many datasets. In fact, sampling from the smoothing spline allows the subsequent parameter inference method to access a larger amount of information and thereby to mitigate noise amplification.

##### 1.2.2. Conversion of ODEs into systems of algebraic equations

If data are available as time series, it is mathematically feasible and beneficial to estimate slopes (for instance, from smoothing splines) and to convert the inference problem from one based on ODEs into one exclusively using algebraic functions (Voit & Savageau, 1982a, 1982b; Varah, 1982; Torres & Voit, 2002; Voit et al., 2005).

Suppose the growth and interaction parameters of an LV system are to be estimated from time series data of the dependent variables *X_i_*. The smoothing of these data facilitates the estimation of slopes *S_k_*(*X_i_*) of all variables at a set of time points *t_k_*, *k* = 1, …, *K*. These time points may or may not correspond to the measured data. In fact, the smoothing permits the computation of slopes at arbitrarily many time points within the observation interval. However many slopes are computed, they correspond to derivatives of the spline of *X_i_* at the given time points. Substituting numerical values of all variables and slopes from the smoothing splines into Eq. (1) yields a system of *n* × *K* linear algebraic equations containing all system parameters:

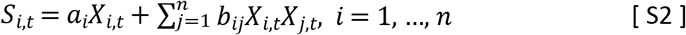

If environmental inputs *γ*_*i*_*X*_*i*,*t*_*U*_*t*_ are to be considered as well, they are added to the equations and either substituted with numerical values, if known, or estimated with the parameters *a*_*i*_ and *b*_*ij*_.

A caveat of this conversion of ODEs into algebraic equation is a possible time warp (see end of chapter 5 of (Voit, 2017)). The reason is that time is explicitly eliminated from the procedure. Nonetheless, the estimates usually provide good results, or at least good initial guesses for other optimization approaches such as traditional gradient methods.

Suppose the dependent variables are not zero within the dataset obtained from smoothing. If so, we can divide both sides of the *K* equations for *X_i_* in expression [S2] by the value of the dependent variable at the appropriate time point. This step is not mandatory but linearizes the equations. The case of variables with values of zero is typically not very interesting or can be handled by eliminating the variable or parts of the time series.

#### 1.2.3. Parameter inference

Once all differentials are replaced with estimated slopes, the inference of parameter values from LV-models offers two options: because the system of algebraic equations is linear, we may optimize its parameter values through simple multivariate linear regression (ALVI-LR), where we may use data points or iterate the regression with subsets of points, which naturally leads to an ensemble of well-fitting models.

An interesting alternative is to use just *n*+1 of the data points and slopes, if *n* is the number of variables, which results in a system of linear equations that can be solved with simple algebraic methods (ALVI-MI). Choosing different data points naturally creates ensembles of solutions. These can be further analyzed, for instance, with respect to model robustness and identifiability. They can also be used to determine to what degree the LV format is adequate for the available data (Voit et al., 2021).

##### 1.2.4. Example of parameter estimation with ALVI

To explain the parameter estimation procedure with ALVI, we use the sparse noisy dataset presented in *Supplements* Section 2 and also in Table S1.2. First, we smooth the data with a spline or LOESS. For this example, we use 5, 8, 11 and 5DF-splines for *X*_1_, *X*_2_, *X*_3_ and *X*_4_ respectively and compute the first derivative of the splines to estimate the slopes. At this point we discard the data and only use the spline values. We may use the original data, especially if we think they characterize the studied phenomenon well, but using the spline usually produces better results in the case of noisy data.

As an example, consider the first differential equation and the first datapoint, at *t* = 0:

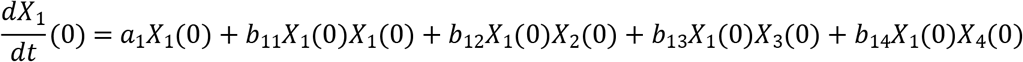

We substitute numerical values for the slope and for all variables on the system equations,

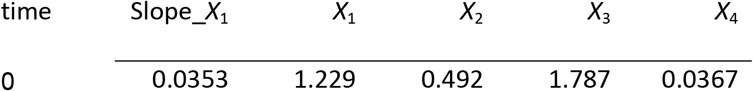

which yields

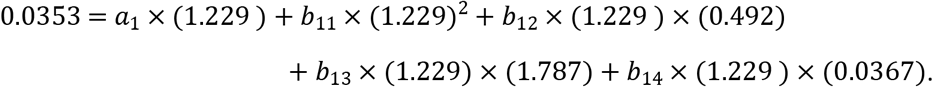

We may divide the equation by the numerical value of the dependent variable, but this step is not mandatory.

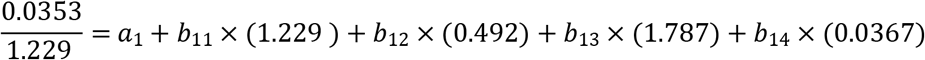

The same steps are performed for every equation and every chosen time point. The result is a system of linear equations with as many equations as chosen time points; each equation has *n*+1 unknown parameters, where *n* corresponds to the number of dependent variables.

Now we have two options: We may use linear regression (ALVI-LR) or matrix inversion (ALVI-MI). ALVI-LR uses every equation and every chosen time point and performs linear regression to produce estimates for the parameters. For the alternative, ALVI-MI, we choose a sample of data points that, when combined with the equations, generates a number of equations equal to the number of parameters to be estimated. If these equations are linearly independent, the system is solvable and the solution is unique, allowing us to obtain estimates for the parameters by simple matrix inversion.

#### 1.3. Multivariate Autoregressive (MAR) models

Multivariate Autoregressive (MAR) models are discrete recursive models. Their format is shown in eqn. 2 and 3 of the main text and conveys that the state of the system at time *t*+1 depends on the state at *t* and possibly on environmental or stochastic input. As an alternative to this modeling structure with “memory 1,” it is possible to extend MAR models to depend also on states farther in the past, such as *X_t_*_- 1_, *X_t_*_-2_, and *X_t_*_-3_, in addition to *X_t_*. However, the commonly used models depend only on the immediately prior state and are sometimes called MAR(1). Here, we only consider MAR(1) models and refer to them simply as MAR models.

MAR models can be interpreted in two distinct ways. In generic mathematical terms, MAR models are stochastic, linear approximations of nonlinear dynamic systems that evolve over time in the vicinity of a fixpoint (steady state). According to this interpretation (Holmes et al., 2013), *x_t_* is a vector of the realization of random variables at time *t*. Noise captures natural variations in environmental conditions and is modeled by a multivariate normal distribution with mean zero and variance-covariance matrix ***δ***. If stochasticity is omitted, MAR models are quite similar to LV models close to the steady state (see *Supplements* Section 1.5).

One may also interpret MAR models in an ecological context, where they can be viewed as multispecies competition models with Gompertz density dependence and an instantaneous growth rate that decreases linearly over time as the population sizes increase (Ives, 1995; Certain, Barraquand, & Gårdmark, 2018). In this context, *x_t_* is a vector of the log-abundances of dependent variables at time *t*.

MAR models may be augmented with state variables that simulate the observation process and these models are called Multivariate Autoregressive(1) State-Space (MARSS) (Holmes et al., 2012; Certain et al., 2018); we will not analyze these for the sake of simplicity. For the comparisons in this study, we are not considering the influence of environmental variables, so the ***γu***_*t*_ term will be omitted henceforth.

It is considered an advantage in ecology if models explicitly take the influence of environmental factors into account (for details, see (Certain et al., 2018) (Hytti et al., 2006)), which is the case for MAR. The availability of estimation software like MARSS (Holmes et al., 2012, 2020) has greatly increased the appeal of MAR models.

#### 1.4. MARSS

MARSS is a software package for analyzing MAR models with or without log transformation of the dependent variables. Its use requires several steps.

1. Specify key MARSS settings:

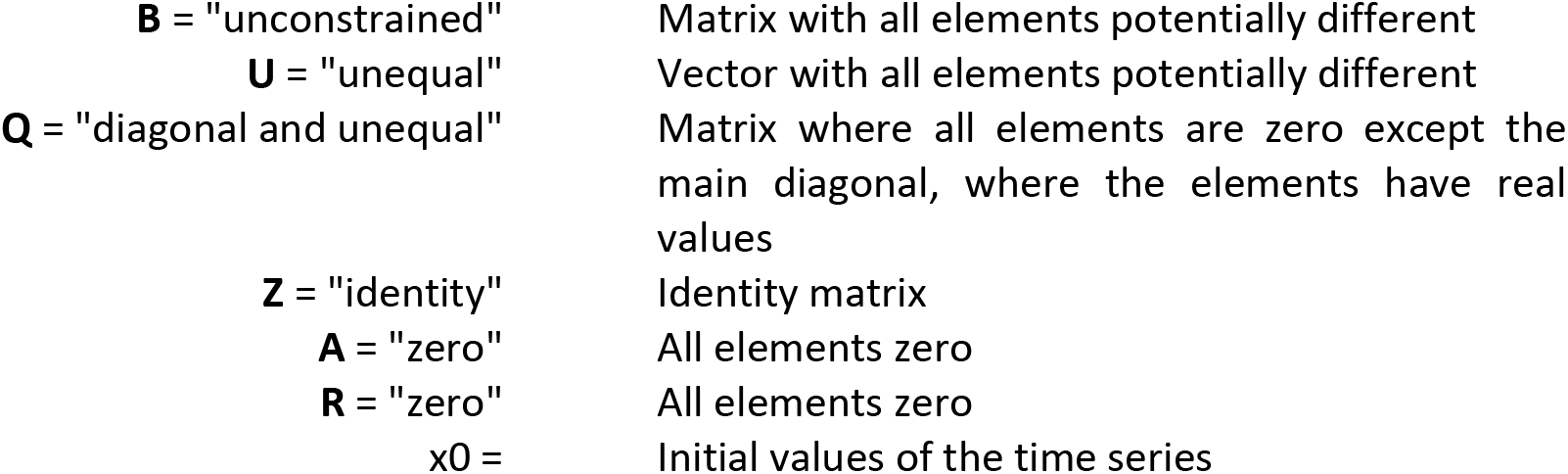 **A** and **R** correspond to the “observation variables,” which simulate the observation process of the system variables. For our comparisons, these are set to zero because observation variables are not considered.
2. In the MARSS function, the data must be formatted with variables in rows and observations in columns. If the data points are not equally distributed in time, they must be augmented by “NA” to force the interval between any two consecutive data entries to be of the same length. This is necessary to ensure the correct time structure of the data for the estimator.
3. With this regularization, MARSS finds estimates for **B**, **U** and **Z**, that correspond to *α*, *β* and *δ* in eqn 3 of the main text.

If MARSS does not converge, it is advisable to increase the max number of iterations. This step solves the problem but is different from the suggestion offered by Holmes and colleagues, namely, that the model assumptions should be checked (see p. 57 in (Holmes et al., 2020)).

Our setup is exactly equal to that used by Holmes *et al*. (Holmes et al., 2012, 2020) for the Isle Royale dataset, which the authors used to exemplify the inference of species interaction parameters with and without covariates. Some of the illustration examples were modeled differently in the literature but for purpose of comparisons with LV models, this model structure was used. For example, the ‘gray whales’ dataset was modeled by Holmes and colleagues with ***β***, the species interaction matrix, set to zero, whereas **R**, the matrix that captures the noise from the observational process, was estimated from the data. Because we are interested in the interactions between species, we do not focus on observational noise, and Holmes’ original setup was replaced with the one discussed above.

#### 1.5. Structural Similarities between Modeling Formats

The two modeling formats appear to be very different mathematically. Nonetheless, they can be compared in terms of their mathematical representations and also with respect to practical considerations. These comparisons demonstrate that the two models can actually behave quite similarly if the community of populations operates relatively close to a stable steady state. By contrast, if this assumption is violated, the two models often show strongly diverging results, as the linearity of the MAR model can deviate considerably from the nonlinearities of the LV model.

Purely considered on mathematical grounds, MAR is defined recursively in eqn 2 and 3 of the main text. By omitting environmental variables, we directly obtain

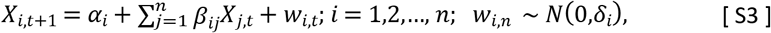

which can be interpreted as a multispecies competition model with Gompertz density dependence, if the dependent variables represent logarithmic abundancies (Certain et al., 2018).

Suppose that the MAR model indeed uses log-abundances. To explain similarities between the MAR and LV formats, we rewrite this multispecies log-abundance model equivalently in Cartesian form, which yields the following:

For *i* = 1,2,…, *n*; *w_i,n_* ∼ *N*(0,*d*_*i*_):

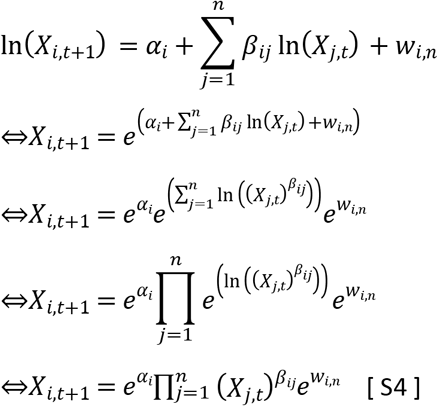

In this form, MAR is similar to a discrete multivariate power-law function.

The similarity of this result to the LV model can be seen if we use Euler’s method for determining the numerical solution. Euler’s method is an approximation of more sophisticated methods and its simplicity makes it preferable for the comparisons between time series and ODEs.

Formulating the typical Euler step for the LV model transforms the ODE into a series of discrete steps of the type

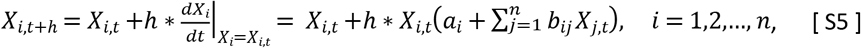

where *h* is the step size of Euler’s method and *dX_i_/dt* is the left-hand side of the differential equations in eqn 1, evaluated at time *t*.

A comparison of eqn [ S4 ] and [ S5 ] suggests that the MAR and LV models seem to be very different. Whereas the LV model captures nonlinear dynamic behaviors without variable transformations, the MAR model uses linearity in log space. Nonetheless, there are similarities between the two formats. To see these, we compare eqn [ S3 ] and [ S5 ] instead of [ S4 ] and [ S5 ].

Furthermore, we suppose that the dynamics is near the steady state of the differential equations, so that *Xi*,*t*+1 − *Xi*,*t* ≈ 0 for any given *t*. Using this approximate equality in eqn [ S3 ], we obtain

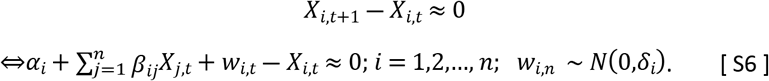

Using this approximate equality in format [ S5 ] for LV yields, for *i* = 1,2,…, *n*:

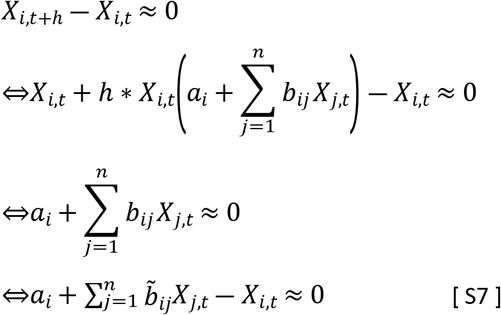

Here *b*_*ij*_ equals *b_ij_* for aIl *i* ≠ *j* and *b_ij_* + 1 Ior *i* = *j*.

If we disregard the Gaussian noise, *w_ij_*, in the MAR model, the two sets of near-steady-state eqn [ S6 ] and [ S7 ] are the same. They are both linear, although the dynamic LV model itself is non-linear. Thus, if *α_i_ = a_i_*, *β_ij_ = b_ij_* for all *i ≠ j* and *β_ij_ = b_ij_* + 1 for *i* = *j*, the MAR and LV models are mathematically equivalent at the steady state and similar close to it. As long as the nonlinearity are close to linear or close to power-law functions, MAR without or with log-transformation, respectively, may be expected to lead to acceptable fits.

### 2. Case study 1: Synthetic LV data

As a representative example, we use the four-variable LV system

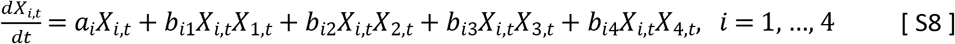

The parameters are presented in Figure S1. For a first analysis, we use this system to create one set of synthetic time courses, consisting of 100 time points, which is presented in Table S1. The dynamics is shown in Figures 1 and S1 as circles.

If we use these noise-free data, the inferences are close to perfect with respect to the trajectories and parameter values (Figure S9).

To mimic a more realistic scenario, we created a noisy dataset, visualized in Figure S1a, which was constructed by randomly choosing forty of the one hundred original datapoints obtained from the synthetic system and adding to the chosen points a normal random variable with mean 0 and a standard deviation of 20% of the mean of each variable. This *noisy dataset* is shown in Table S1.2.

A second realistic dataset (Figure S1b) was constructed by first choosing eleven points from the data that characterize the dynamic (including extremes values). Next, each of the chosen points was multiplied by a random normal variable with mean 1 and standard deviation of 0.2. This process was iterated to create five replicates per chosen point. This *replicate dataset* is shown in Table S1.3.

Variable *X*_4_ was designed as a (decoupled) logistic function. It is unaffected by the other variables and does not affect them either. It was included to explore to what degree the methods to be tested can detect this detachment.

The smoothing and slope estimation steps followed directly the procedures described in *Supplements* Section 1.2. The first derivative of the smoothing function was used for estimates for the slopes.

To infer numerical values for the parameters of a given equation, we have the choice between linear regression (ALVI-LR) and algebraic matrix inversion (ALVI-MI). For ALVI-MI, we choose points from the sample and use the corresponding slope estimates to create a system of equations with the same number of equations and unknowns. As we have 4 variables and 20 parameters, we need 20 independent equations and thus 5 time points. For each time point we obtain the value for each of the 4 dependent variables and use these to populate the equations.

As an illustration for the noisy dataset, we choose 5, 8, 11 and 5DF-splines and time points *t* = 8, 13, 17, 19 and 79 for the noisy dataset. For the replicate dataset, we use 5, 6, 8 and 8DF-splines, and the ALVI-MI solution was calculated with spline points at times 11, 18, 26, 33 and 60. The time point selection for the ALVI-MI solution can be automated using a random or exhaustive search of the possibilities.

The noisy dataset (Figures 1a, S1a, S3a and S4a) is representative of a study where each time point sample corresponds to a single observation taken when it is possible or convenient. By contrast, the replicate dataset (Figures 1b, S1b, S3b and S4b) simulates a series of experimental replicates where the observations were conducted multiple times, but at fewer time points, which the researchers suspect would contain valuable information.

As an illustration of how the smoothing techniques work, we used splines and LOESS with different degrees of smoothing. The results for variable X1 are shown in Figure S2. Choosing the optimal degree of smoothing is not a trivial matter. Too much smoothing ignores important details in the variable dynamics, while too little incorporates noise. In the programing language R, the function “loess.as” allows the calculation of the optimum value for the spam, which controls the smoothing. The user still must decide the degree of the polynomials to be used and choose from two criteria for automatic smoothing parameter selection: a bias-corrected Akaike information criterion (AICC) and generalized cross-validation (GCV). This choice is not always leading to the optimal solution, but it should be used to challenge our assumptions.

Figure S3 presents the same treatment as presented in Figure 1 but for datasets with 5% noise. It is clear and not surprising that the models produce better quality fits when the signal-do-noise ratio is higher.

#### 2.1.1. Application of ALVI to synthetic data

An alternative to using the algebraic parameter inference method with matrix inversion (ALVI-MI) is the linear regression method (ALVI-LR); fits for 20% noise are shown in Figure S4. As in ALVI-MI, the parameter values are close to the true values and the fit is acceptable. However, the dynamics of the system is slightly different. One reason is that the dynamic solutions are sometimes quite sensitive to the chosen initial values. As a remedy, it is often beneficial to initiate the solution somewhere inside the overall time interval, typically close to the midpoints of the variable ranges, and solve forward and backward.

ALVI also works for more complicated dynamics, as can be seen in Figure S5. Here we are interested in finding out if the methods can recover the dynamics, which in some cases turns out to be challenging for sparse data even without the introduction of noise. Thus, we used the synthetic data unaltered. Specifically, data for early time points (*t* = [1, 100]) were fitted and then extrapolated for a total time horizon of (*t* = [1, 500]). In these examples, ALVI-MI is used with 100DF-splines. It uses data samples with points corresponding to timepoints *t* = 5, 10, 20, 30 and 50 for all cases except for the chaotic oscillations where we used *t* = 4, 6, 10, 15 and 35. For each case, we also present the MAR estimates. Of course, one must recall that the original data were produced with LV models. While the MAR model extrapolations are not always satisfactory, it is nevertheless comforting that the inference method returns good results for the time interval used for data fitting.

ALVI-MI generally performed very well but did not adequately capture the deterministic chaos (chaos 1). For this case only, we obtained a better fit using ALVI-LR, which may not be surprising given the extremely sensitive nature of chaotic systems to noise. Apart from this situation, results with ALVI-LR are very similar to ALVI-MI results and will not be displayed.

In Figure S5b, the MAR model performed well when log-abundances were used. In the remaining cases, it failed to replicate the oscillations or these exploded by reaching amplitudes far bigger than in the dataset. One also notes early discrepancies between the initial points used to create the estimates and the MAR estimates.

### 3. Supplemental Figures

**Figure S1:**
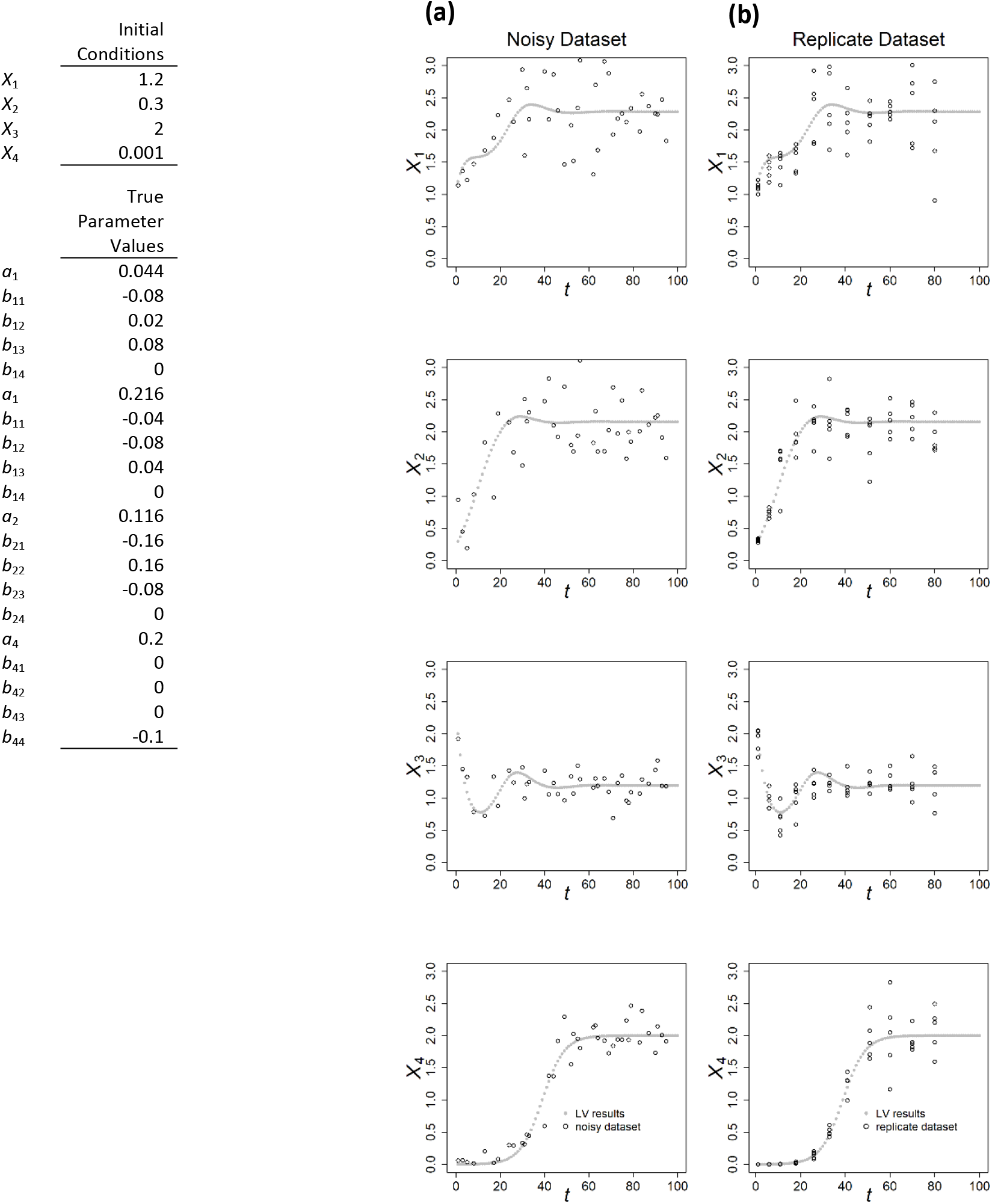
Time courses with superimposed noise. Initial conditions and parameter values for the synthetic LV example in equation S8 with four dependent variables. **Column a:** Noisy dataset Based on 40 points from the synthetic data with added random normal noise with mean 0 and standard deviation equal to 20% of each variable mean. **Column b:** Replicate dataset 11 points were chosen from the synthetic data and at each point five “observations” were created by multiplying the value of the variable by a random normal value of mean 1 and standard deviation of 0.2.

**Figure S2:**
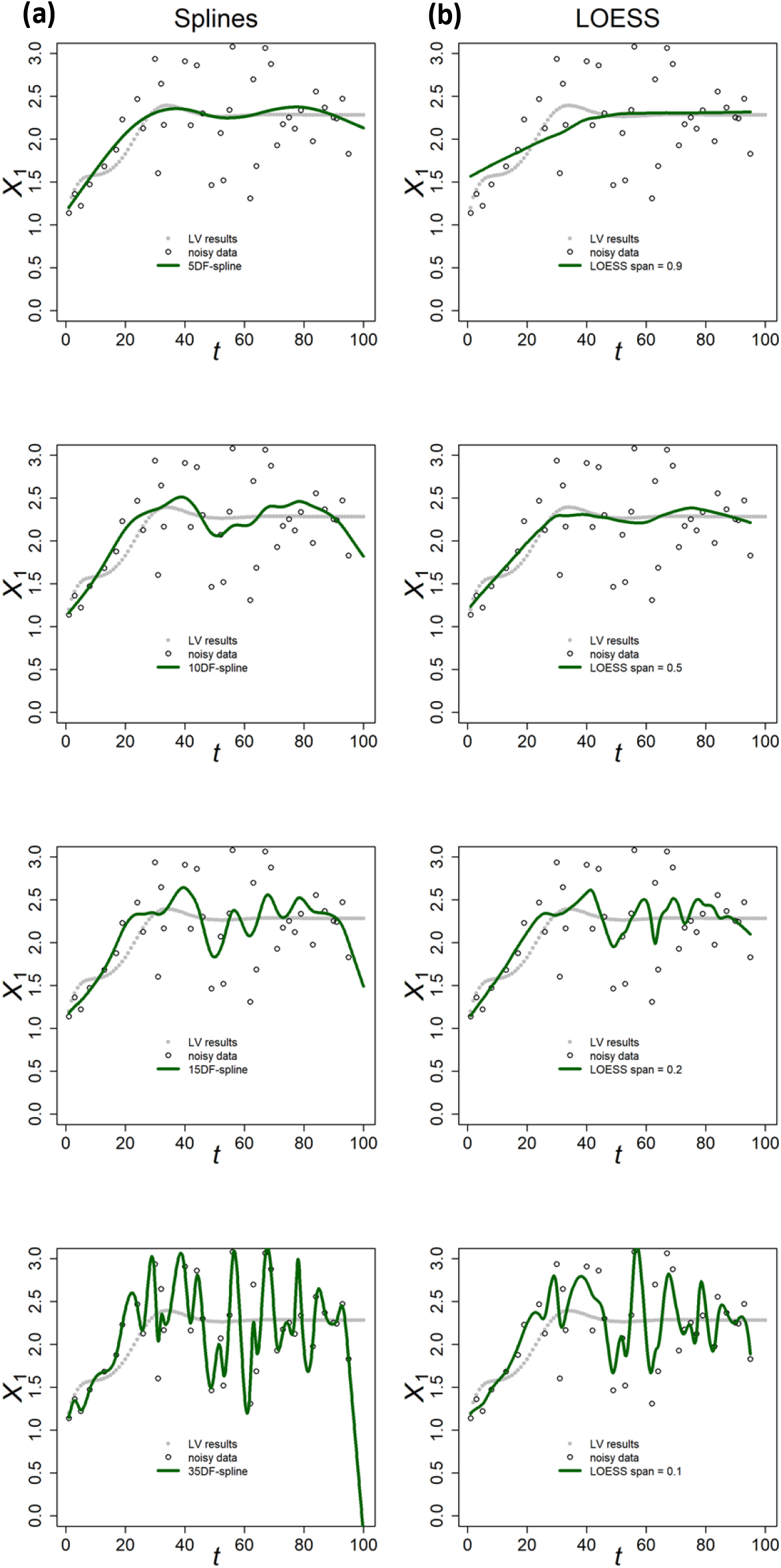
Smoothing noisy variable *X*_1_. **Column a:** Splines with different degrees of freedom. **Column b:** LOESS with different span levels.

**Figure S3:**
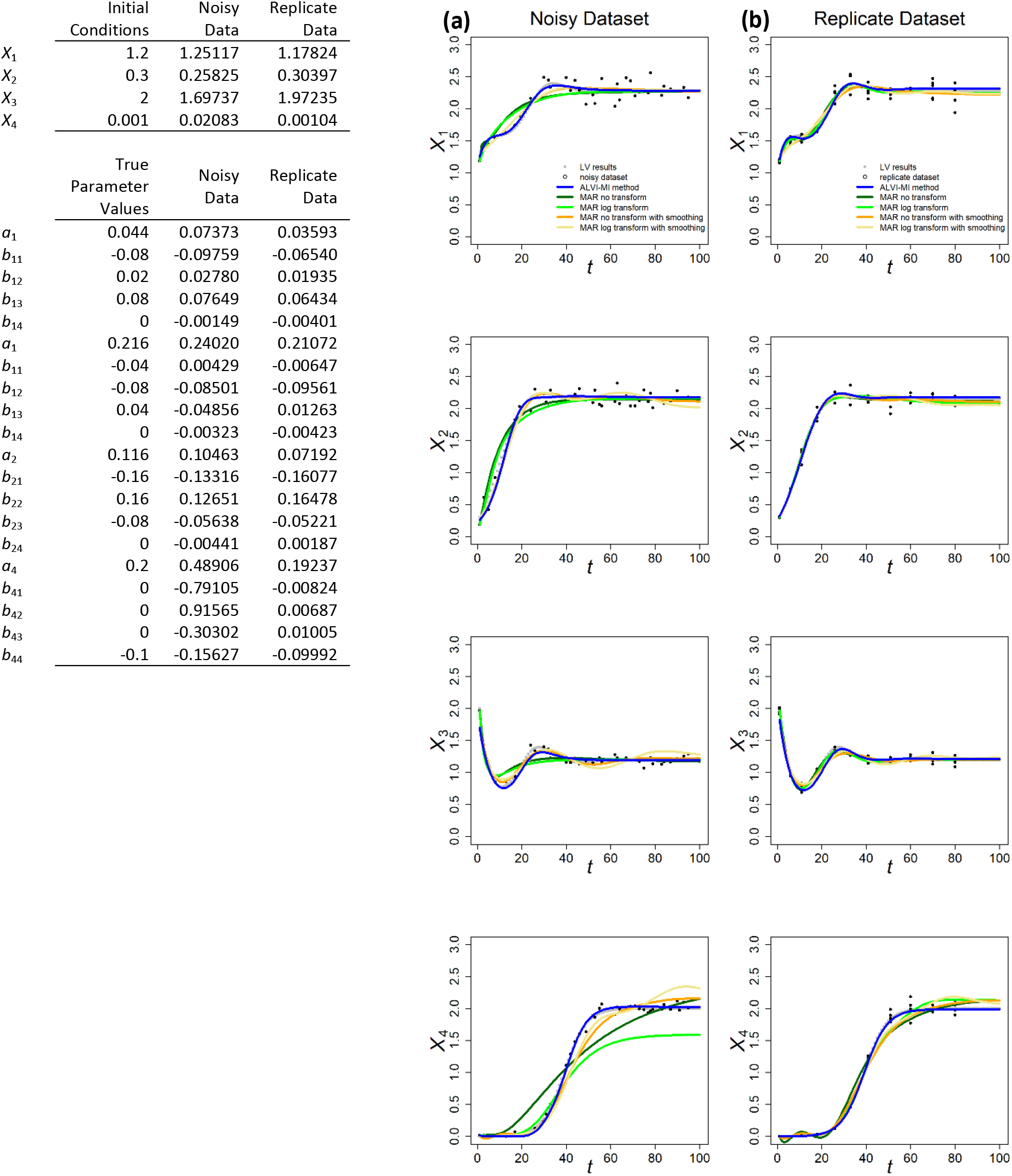
Results of ALVI-MI and MAR applied to the noisy and replicate datasets with 5% of noise. **Column a:** Noisy dataset. All variables were smoothed with 11DF-splines and the ALVI-MI solution was calculated with spline points at time points 8, 17, 30, 42 and 62. **Column b:** Replicate dataset. All variables were smoothed with an 8DF-spline and the ALVI-MI solution was calculated with spline points corresponding to time 6, 18, 26, 33 and 60. MAR estimates are presented in Tables S2.1 and S2.2.

**Figure S4:**
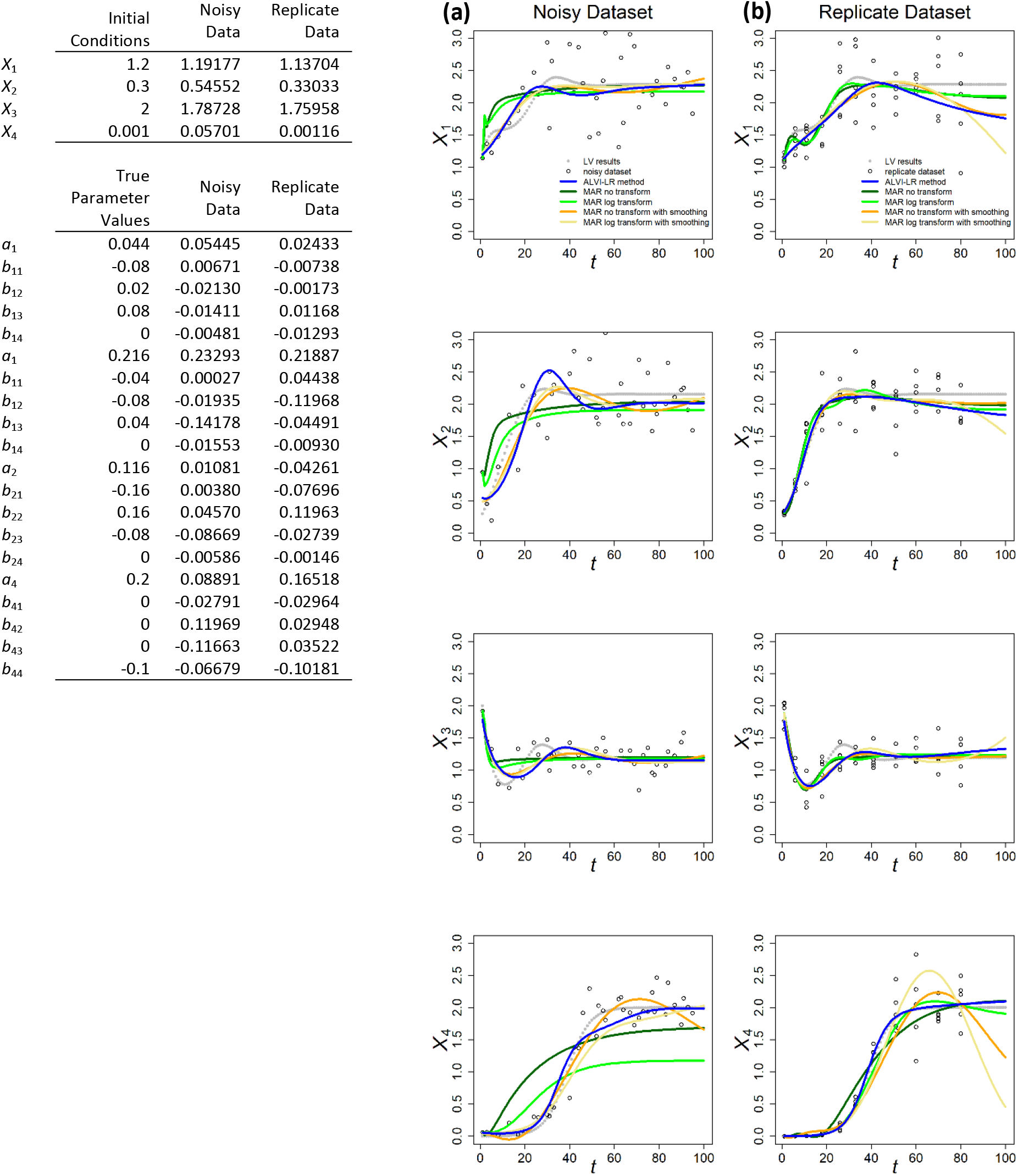
Results of ALVI-LR and MAR applied to the noisy and replicate datasets. **Column a:** Noisy dataset. Time courses of *X*_1_, *X*_2_, *X*_3_ and *X*_4_ were smoothed with 6, 11, 11 and 11DF-splines respectively. **Column b:** Replicate dataset. All variables were smoothed with 8DF-splines. MAR estimates are the same presented in Figure 1.

**Figure S5.**
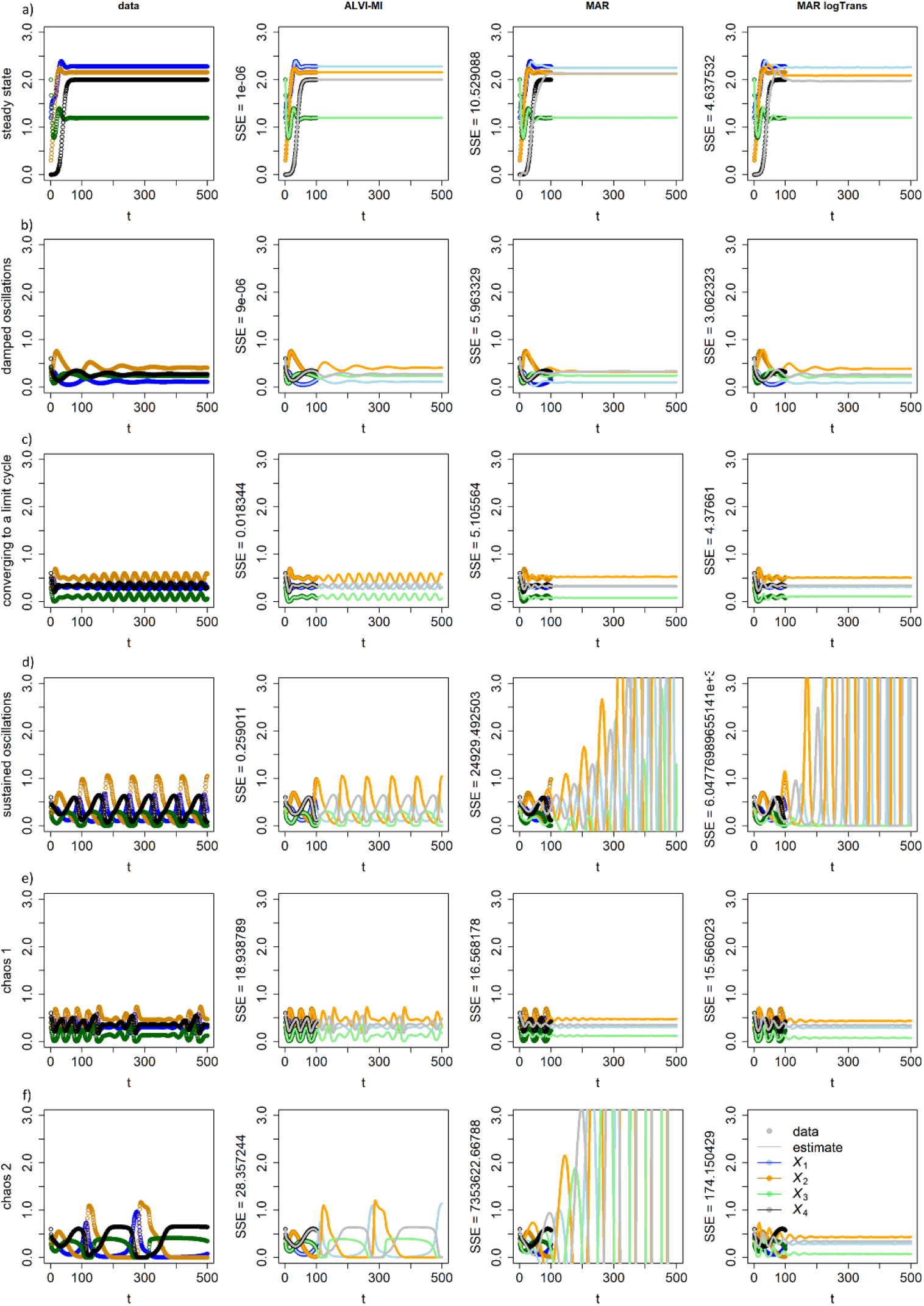
Data and results of inferences with ALVI-MI and MAR methods for LV systems with different dynamics. **Row a:** Data converging to a stable steady state; **Row b:** Damped oscillations; **Row c:** Initially erratic oscillations converging to a limit cycle; **Row d:** Sustained oscillations; **Row e:** Deterministic chaos, example 1; **Row f:** Deterministic chaos, example 2. Data, ALVI-MI and MARSS estimates are presented in Table S3. The SSEs concerning the differences between the data and estimates for *t* = [1, 500] are presented as labels to the Y-axis. No smoothing preceded MAR because the data are noise free.

**Figure S6.**
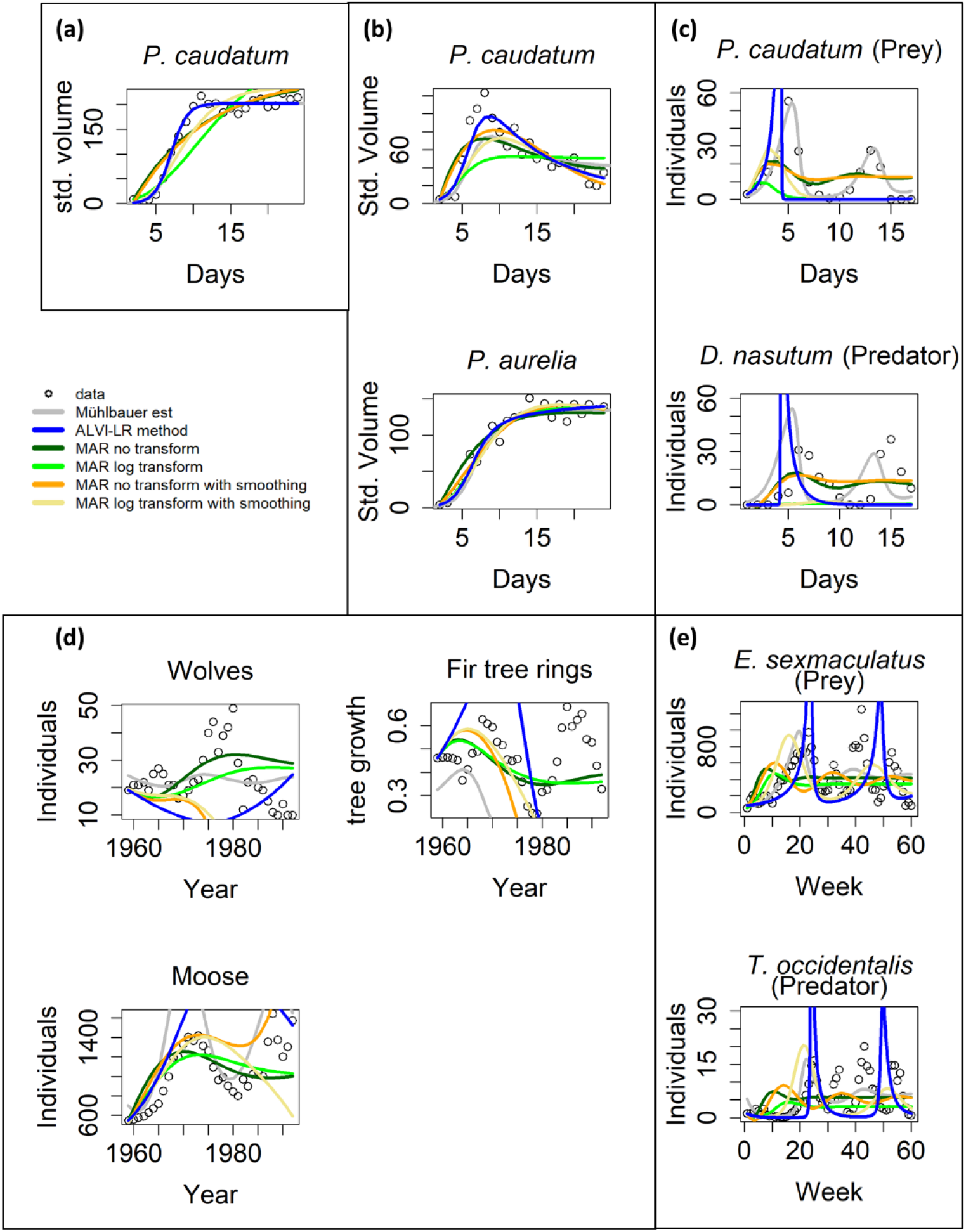
Examples of experimental data analyzed with ALVI-LR and MAR. Black lines are estimates from Mühlbauer *et al*. (Mühlbauer et al., 2020). ALVI-LR estimates are represented as blue lines; corresponding parameter values can be seen in Table S4.2. MAR estimates are presented in green, orange and yellow. Parameter estimates are presented in Table S4.3. **a:** Standardized volume of *Paramecium caudatum* grown in monoculture (Gause, 1934). **b:** Standardized volume of *Paramecium caudatum* and *Paramecium aurelia* grown in co-culture (Gause, 1934). **c:** Predator-prey interactions between *Didinium nasutum* and *Paramecium caudatum* grown in mixture (Gause, 1934). **d:** Multi-trophic dynamics for wolves, moose, and fir tree rings on Isle Royale from 1960 to 1994 (McLaren & Peterson, 1994). **e:** Predator-prey interactions between *Eotetranychus sexmaculatus* and *Typhlodromus occidentalis* in a spatially structured experiment (Huffaker, Shea, & Herman, 1963).

**Figure S7.**
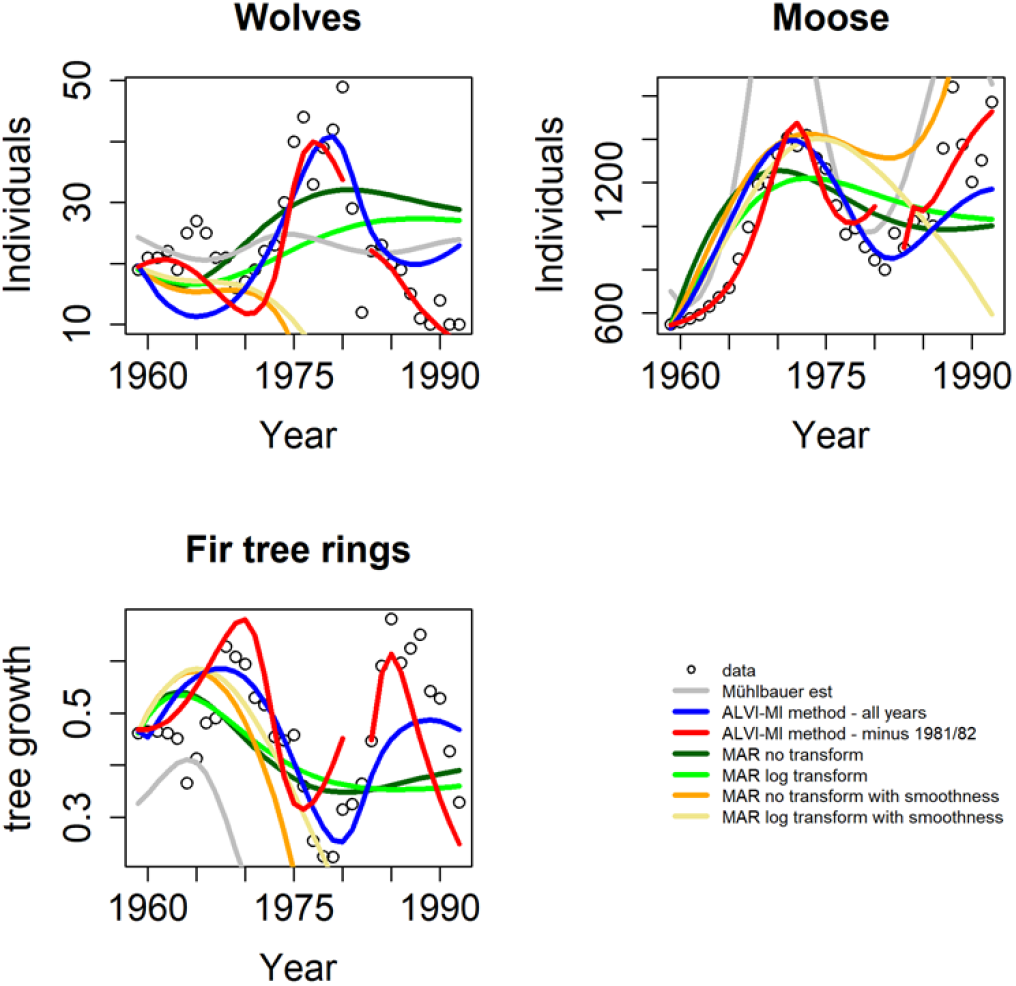
Multi-trophic dynamics for wolves, moose, and fir trees on Isle Royale from 1960 to 1994, from McLaren & Peterson (1994) (McLaren & Peterson, 1994). This panel is very similar to Figure 2 d) but contains additional information. ALVI-MI estimates using all data are represented as blue lines. MAR estimates are presented in green, orange and yellow. Red lines correspond to the estimates using ALVI-MI for two intervals, from 1959 to 1980 and form 1983 until the end of the series. This split was tested because around 1980 the wolves were exposed to a disease that drastically reduced their numbers, an event that dynamic models do not capture outside piecewise operation. MAR estimates are the same as in Figure 3.

**Figure S8:**
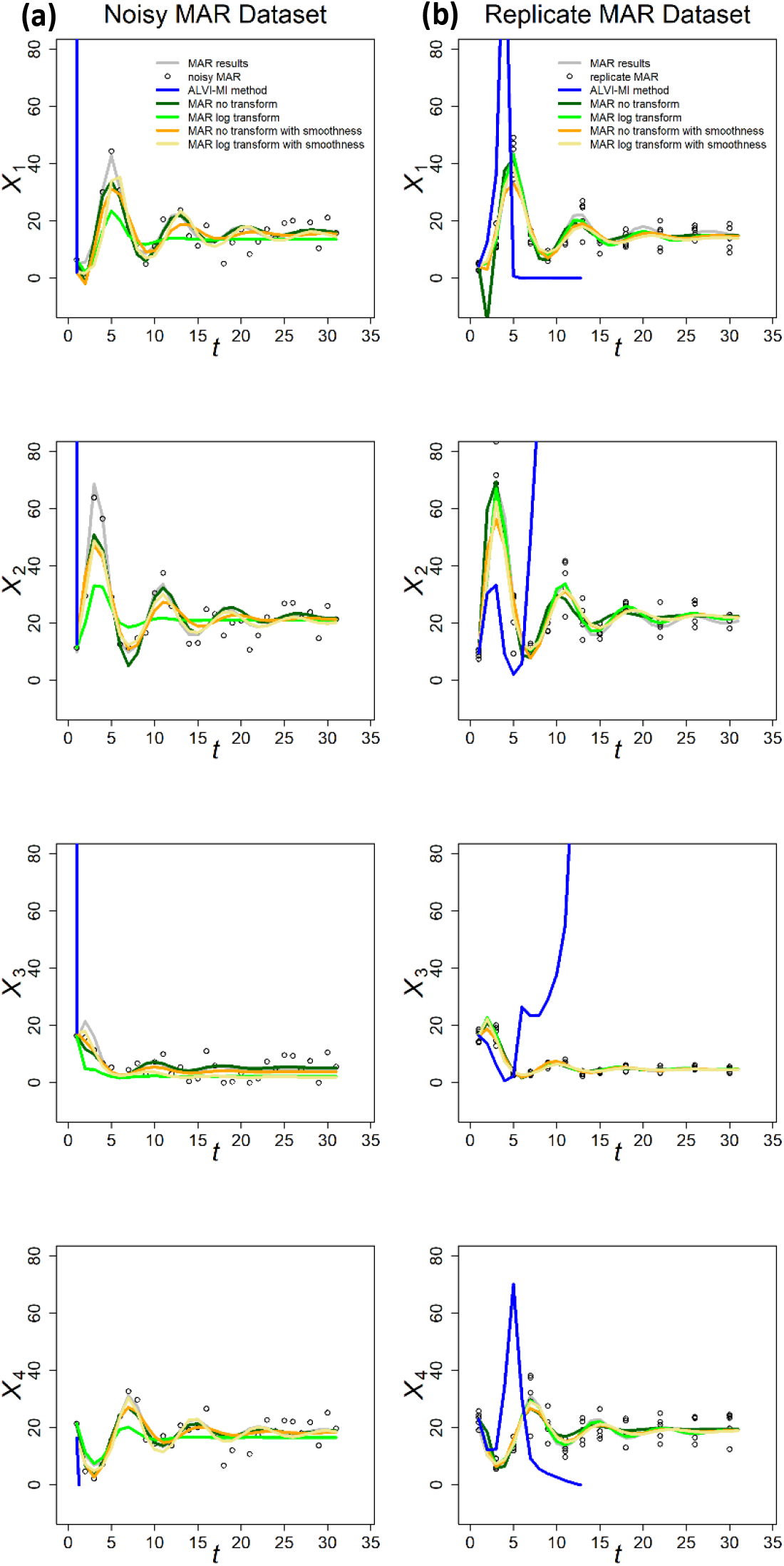
ALVI-MI and MAR applied to an alternative sample of the same data presented in Figure 4, but with slightly changed noise. Although the differences in noise are visually almost undetectable, very different results for the ALVI-MI fit are obtained if the same sample of spline points is used. However, if a new sample of spline points is determined, the fits are almost indistinguishable (not shown). See Text for further explanations.

**Figure S9:**
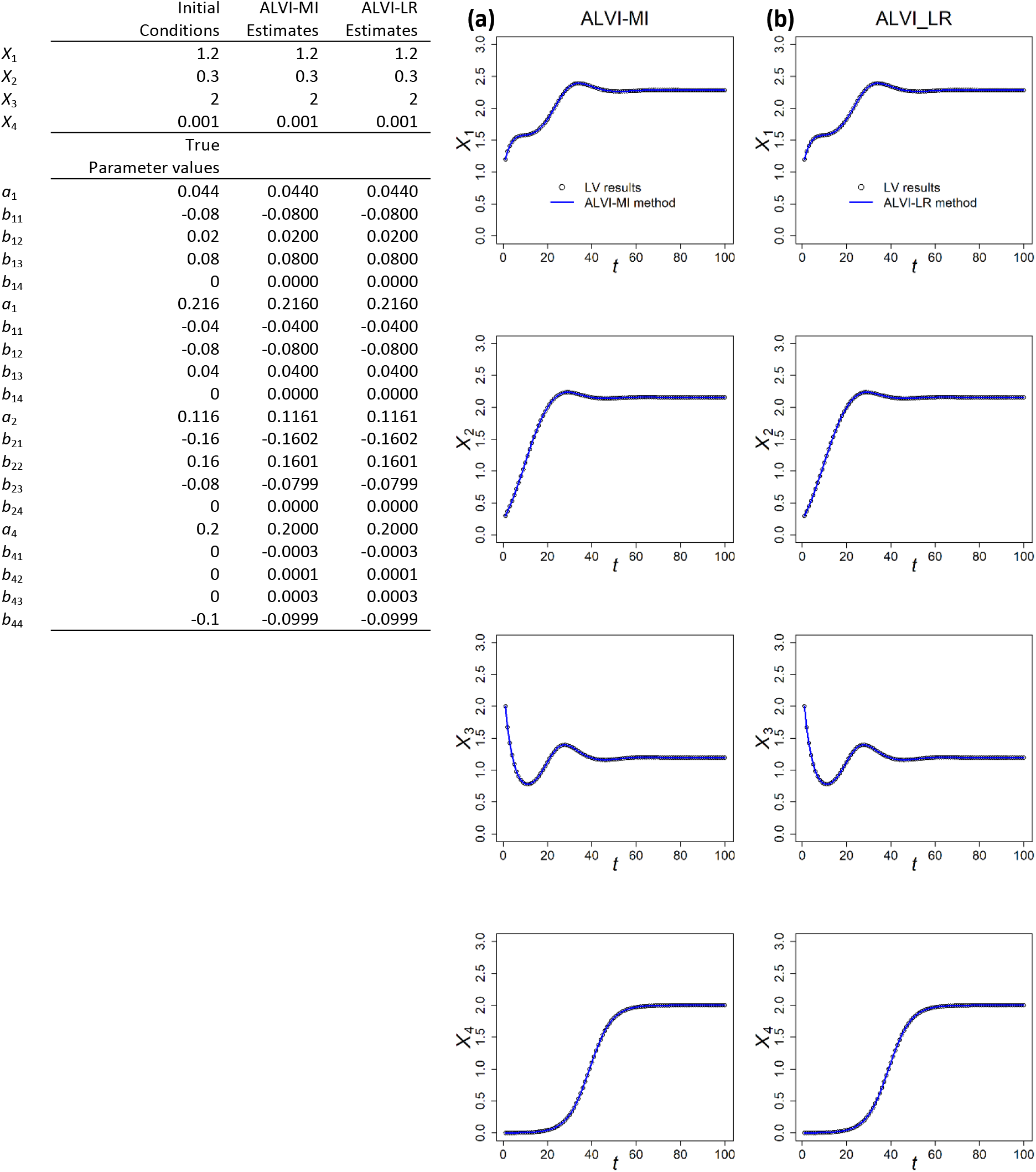
Estimates with alternative ALVI methods. **Column a:** ALVI-MI and **Column b:** ALVI-LR with original synthetic LV data. The fits are of high quality (ALVI-MI SSE = 1.162229e-05 and ALVI-LR SSE = 3.289283e-07) and the parameter estimates are very close to the true parameters.

### 4. Supplemental Tables

**Table S1.1.**
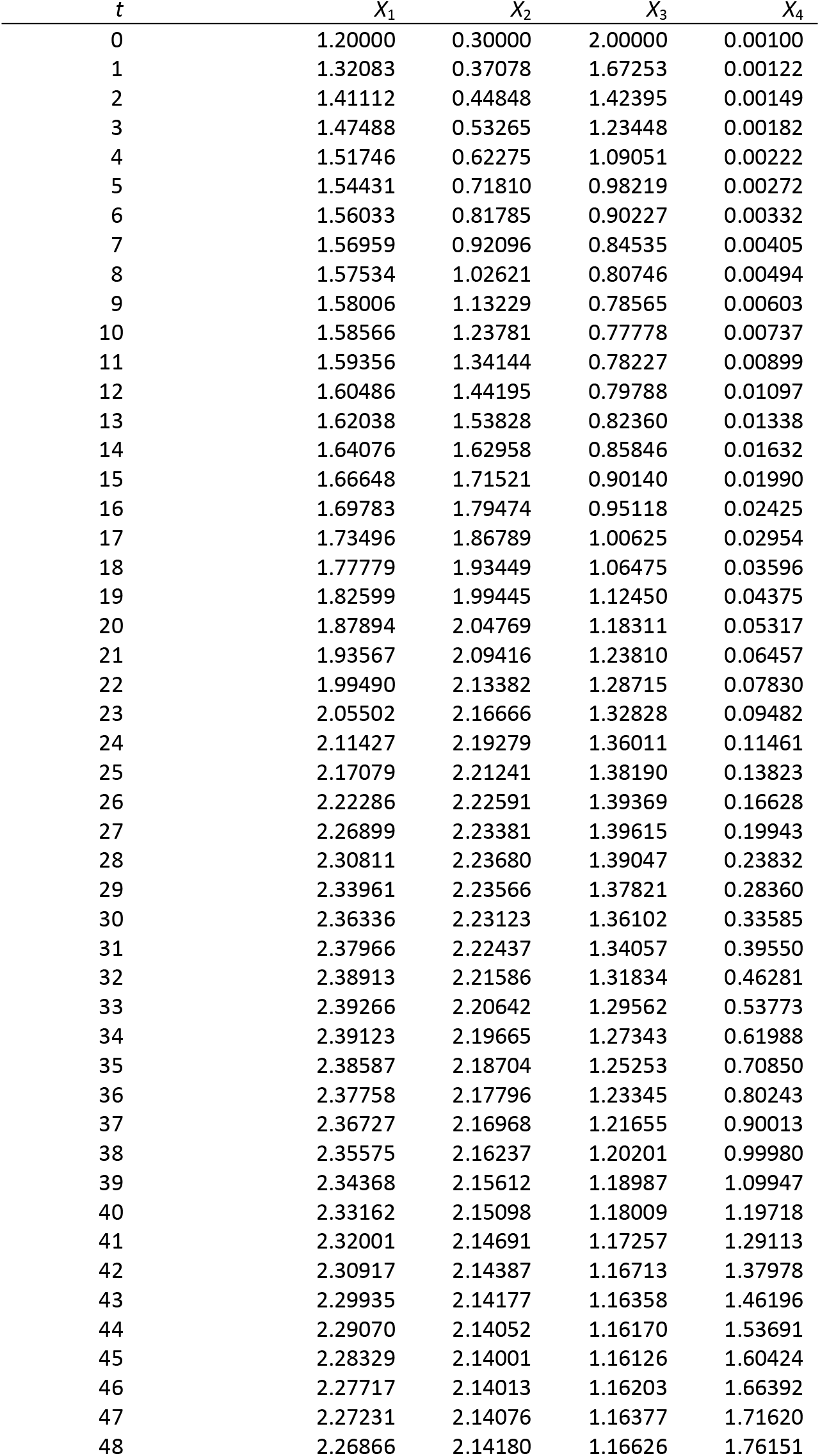
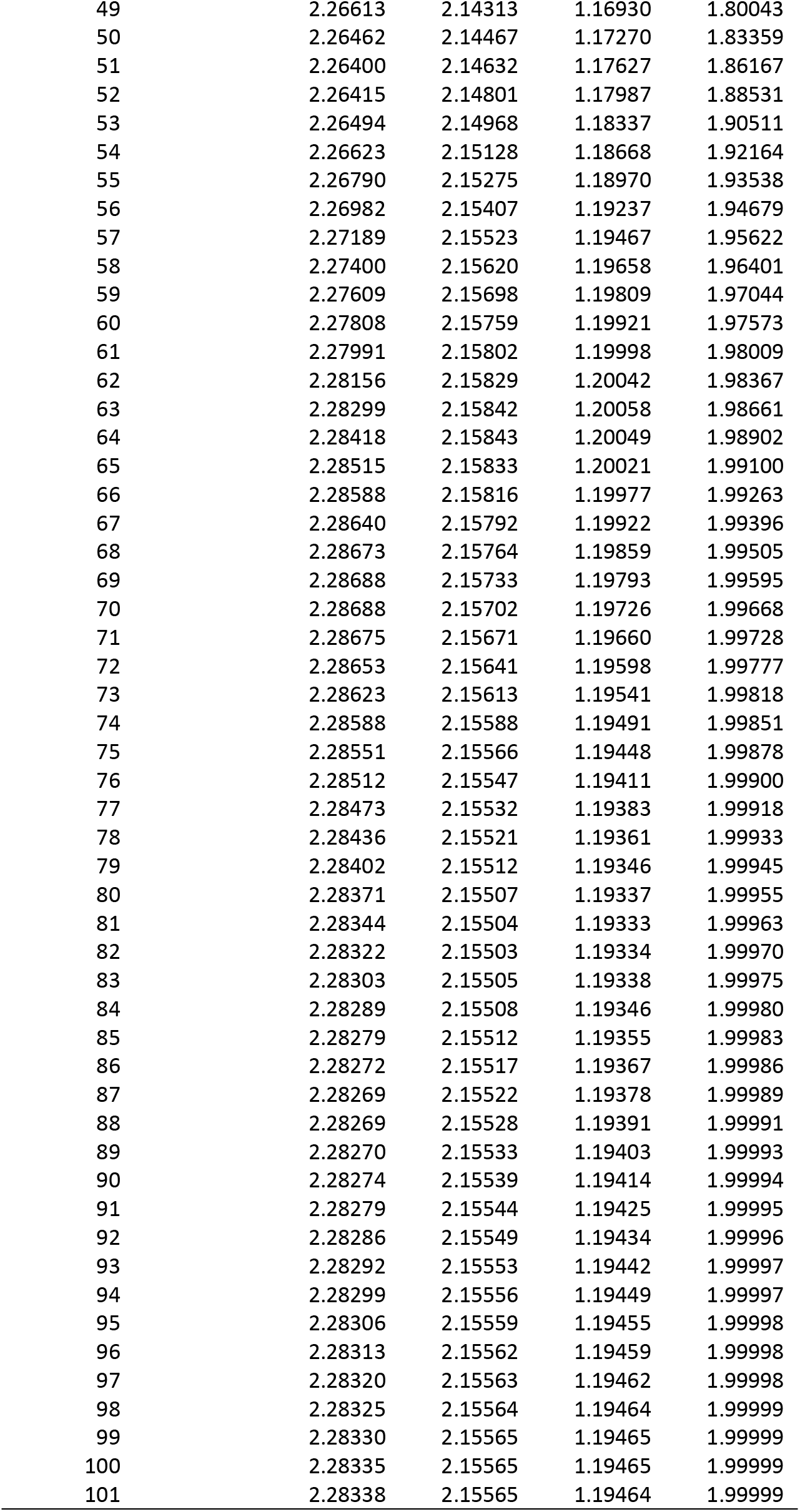
Synthetic LV data. The data were generated with an LV system with four dependent variables with parameter values presented in Figure S1.

**Table S1.2.**
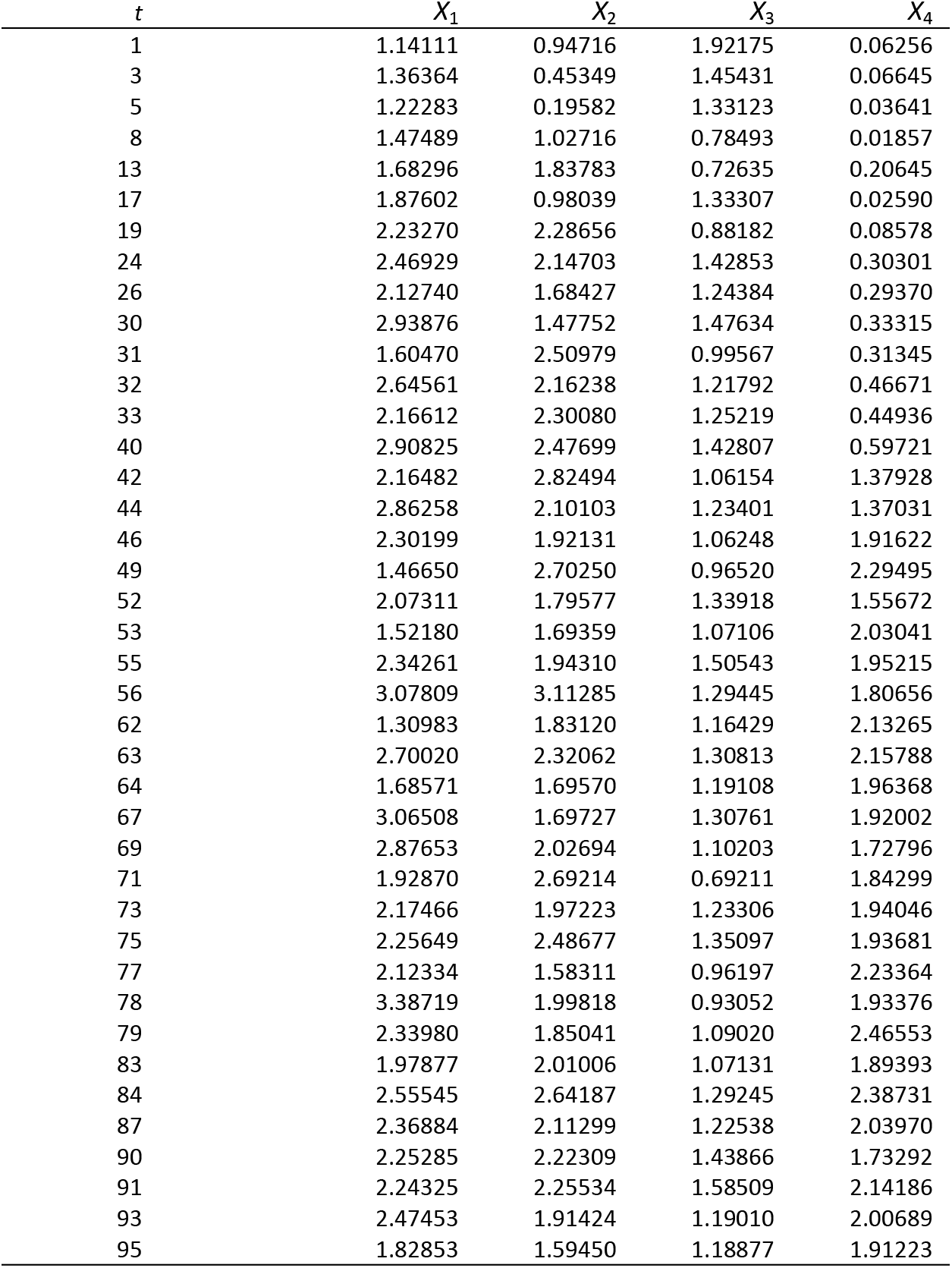
Noisy LV dataset. From the synthetic data, generated with the LV system in Table S1.1, forty values were selected and random normal noise was added with mean 0 and standard deviation equal to 20% of each variable mean.

| $t$ | $X_1$ | $X_2$ | $X_3$ | $X_4$ |
| --- | --- | --- | --- | --- |
| 1 | 1.14111 | 0.94716 | 1.92175 | 0.06256 |
| 3 | 1.36364 | 0.45349 | 1.45431 | 0.06645 |
| 5 | 1.22283 | 0.19582 | 1.33123 | 0.03641 |
| 8 | 1.47489 | 1.02716 | 0.78493 | 0.01857 |
| 13 | 1.68296 | 1.83783 | 0.72635 | 0.20645 |
| 17 | 1.87602 | 0.98039 | 1.33307 | 0.02590 |
| 19 | 2.23270 | 2.28656 | 0.88182 | 0.08578 |
| 24 | 2.46929 | 2.14703 | 1.42853 | 0.30301 |
| 26 | 2.12740 | 1.68427 | 1.24384 | 0.29370 |
| 30 | 2.93876 | 1.47752 | 1.47634 | 0.33315 |
| 31 | 1.60470 | 2.50979 | 0.99567 | 0.31345 |
| 32 | 2.64561 | 2.16238 | 1.21792 | 0.46671 |
| 33 | 2.16612 | 2.30080 | 1.25219 | 0.44936 |
| 40 | 2.90825 | 2.47699 | 1.42807 | 0.59721 |
| 42 | 2.16482 | 2.82494 | 1.06154 | 1.37928 |
| 44 | 2.86258 | 2.10103 | 1.23401 | 1.37031 |
| 46 | 2.30199 | 1.92131 | 1.06248 | 1.91622 |
| 49 | 1.46650 | 2.70250 | 0.96520 | 2.29495 |
| 52 | 2.07311 | 1.79577 | 1.33918 | 1.55672 |
| 53 | 1.52180 | 1.69359 | 1.07106 | 2.03041 |
| 55 | 2.34261 | 1.94310 | 1.50543 | 1.95215 |
| 56 | 3.07809 | 3.11285 | 1.29445 | 1.80656 |
| 62 | 1.30983 | 1.83120 | 1.16429 | 2.13265 |
| 63 | 2.70020 | 2.32062 | 1.30813 | 2.15788 |
| 64 | 1.68571 | 1.69570 | 1.19108 | 1.96368 |
| 67 | 3.06508 | 1.69727 | 1.30761 | 1.92002 |
| 69 | 2.87653 | 2.02694 | 1.10203 | 1.72796 |
| 71 | 1.92870 | 2.69214 | 0.69211 | 1.84299 |
| 73 | 2.17466 | 1.97223 | 1.23306 | 1.94046 |
| 75 | 2.25649 | 2.48677 | 1.35097 | 1.93681 |
| 77 | 2.12334 | 1.58311 | 0.96197 | 2.23364 |
| 78 | 3.38719 | 1.99818 | 0.93052 | 1.93376 |
| 79 | 2.33980 | 1.85041 | 1.09020 | 2.46553 |
| 83 | 1.97877 | 2.01006 | 1.07131 | 1.89393 |
| 84 | 2.55545 | 2.64187 | 1.29245 | 2.38731 |
| 87 | 2.36884 | 2.11299 | 1.22538 | 2.03970 |
| 90 | 2.25285 | 2.22309 | 1.43866 | 1.73292 |
| 91 | 2.24325 | 2.25534 | 1.58509 | 2.14186 |
| 93 | 2.47453 | 1.91424 | 1.19010 | 2.00689 |
| 95 | 1.82853 | 1.59450 | 1.18877 | 1.91223 |

**Table S1.3.** Replicate LV dataset. 11 points were selected from the synthetic data in Table S1.1. For each time point, five observations were created by multiplying the original value by a normal random variable with mean 1 and standard deviation 0.2.

| $t$ | $X_1$ | $X_2$ | $X_3$ | $X_4$ |
| --- | --- | --- | --- | --- |
| 1 | 1.079474 | 0.307892 | 1.968433 | 0.001177 |
| 1 | 1.228073 | 0.319118 | 1.767284 | 0.001143 |
| 1 | 1.001938 | 0.278408 | 2.035955 | 0.001019 |
| 1 | 1.151608 | 0.34439 | 2.049352 | 0.000994 |
| 1 | 1.106675 | 0.330651 | 1.634474 | 0.001462 |
| 6 | 1.409001 | 0.827832 | 1.033651 | 0.003136 |
| 6 | 1.292779 | 0.655128 | 0.840712 | 0.002841 |
| 6 | 1.186731 | 0.753583 | 0.964293 | 0.003671 |
| 6 | 1.501709 | 0.702128 | 0.846646 | 0.002595 |
| 6 | 1.600803 | 0.778034 | 1.191477 | 0.003243 |
| 11 | 1.553429 | 1.585188 | 0.501395 | 0.008283 |
| 11 | 1.420026 | 1.565142 | 0.721249 | 0.009309 |
| 11 | 1.599542 | 0.772725 | 0.708241 | 0.004804 |
| 11 | 1.642382 | 1.707548 | 0.424371 | 0.00881 |
| 11 | 1.142047 | 1.689577 | 0.992655 | 0.00613 |
| 18 | 1.644046 | 1.84217 | 0.929999 | 0.044789 |
| 18 | 1.780007 | 1.601518 | 1.134646 | 0.030728 |
| 18 | 1.710696 | 1.833336 | 1.096591 | 0.023249 |
| 18 | 1.331607 | 2.483739 | 0.591253 | 0.029612 |
| 18 | 1.357593 | 1.968956 | 1.2092 | 0.017282 |
| 26 | 2.560157 | 2.19029 | 1.010075 | 0.084837 |
| 26 | 2.478865 | 2.142542 | 1.441704 | 0.160821 |
| 26 | 2.920662 | 2.166496 | 1.227926 | 0.177711 |
| 26 | 1.783109 | 1.700208 | 1.23534 | 0.205836 |
| 26 | 1.809358 | 2.395388 | 1.056138 | 0.105769 |
| 33 | 2.230052 | 2.81995 | 1.194645 | 0.540826 |
| 33 | 1.692464 | 2.038453 | 1.113628 | 0.428625 |
| 33 | 2.981684 | 2.168246 | 1.363852 | 0.486374 |
| 33 | 2.095491 | 1.58247 | 1.231077 | 0.474691 |
| 33 | 2.875616 | 2.102594 | 1.238574 | 0.612312 |
| 41 | 1.970818 | 2.333379 | 1.04227 | 1.296557 |
| 41 | 1.611026 | 1.927812 | 1.114056 | 1.438404 |
| 41 | 2.112647 | 2.279129 | 1.081485 | 0.99357 |
| 41 | 2.652937 | 1.953001 | 1.498288 | 1.30327 |
| 41 | 2.261238 | 2.34695 | 1.170614 | 1.306394 |
| 51 | 2.079631 | 1.228253 | 1.209477 | 2.075646 |
| 51 | 1.819921 | 1.666988 | 1.070121 | 1.644326 |
| 51 | 2.454389 | 2.202211 | 1.41537 | 2.439964 |
| 51 | 2.256488 | 2.134285 | 1.231388 | 1.709964 |
| 51 | 2.213276 | 2.102253 | 1.234634 | 1.884561 |
| 60 | 2.165804 | 2.18245 | 1.155609 | 2.283614 |
| 60 | 2.279157 | 1.885294 | 1.137585 | 1.698353 |
| 60 | 2.36829 | 2.52211 | 1.349542 | 2.049815 |
| 60 | 2.234632 | 2.281866 | 1.184983 | 1.165772 |
| 60 | 2.439225 | 1.996245 | 1.501995 | 2.82506 |
| 70 | 1.719859 | 2.411846 | 1.227642 | 1.786889 |
| 70 | 2.570555 | 2.462909 | 1.1756 | 1.878108 |
| 70 | 1.790253 | 1.887747 | 1.142104 | 1.895824 |
| 70 | 2.723166 | 2.042576 | 1.65201 | 1.8243 |
| 70 | 3.007495 | 2.227207 | 0.93787 | 2.226255 |
| 80 | 2.296889 | 2.001374 | 1.396972 | 2.20474 |
| 80 | 2.749139 | 1.714839 | 1.059392 | 1.594539 |
| 80 | 0.9041 | 2.298373 | 1.489557 | 2.267917 |
| 80 | 1.676455 | 1.788501 | 0.766479 | 2.491924 |
| 80 | 2.131265 | 1.738182 | 1.403356 | 1.898047 |

**Table S1.4.** MAR estimates for the noisy LV dataset in Figs. 1 and S4.

|  | MAR without<br>transformation | MAR with log<br>transformation | MAR with<br>smoothing | MAR with log<br>transformation<br>and smoothing |
| --- | --- | --- | --- | --- |
| $\beta_{11}$ | -0.09089 | -0.03360 | 1.06000 | 1.02000 |
| $\beta_{21}$ | 0.27209 | 0.43560 | 0.13800 | 0.07940 |
| $\beta_{31}$ | -0.21012 | -0.43570 | 0.07490 | -0.10500 |
| $\beta_{41}$ | 0.12619 | 0.42840 | -0.33900 | -0.22100 |
| $\beta_{12}$ | 0.43295 | 0.24860 | -0.05360 | -0.01990 |
| $\beta_{22}$ | 0.47619 | 0.62610 | 0.91000 | 0.94700 |
| $\beta_{32}$ | 0.18150 | 0.17100 | 0.01730 | 0.12000 |
| $\beta_{42}$ | 0.00788 | 0.23190 | 0.22900 | 0.19400 |
| $\beta_{13}$ | -0.20530 | -0.01010 | -0.01980 | -0.01510 |
| $\beta_{23}$ | -0.28009 | -0.33410 | -0.07090 | -0.07300 |
| $\beta_{33}$ | 0.50995 | 0.57920 | 0.86700 | 0.90900 |
| $\beta_{43}$ | -0.00869 | -0.02950 | -0.03190 | 0.04240 |
| $\beta_{14}$ | 0.07772 | 0.00960 | -0.00954 | -0.00439 |
| $\beta_{24}$ | 0.06929 | 0.02720 | -0.02780 | -0.01270 |
| $\beta_{34}$ | 0.00957 | 0.01530 | -0.00911 | -0.00807 |
| $\beta_{44}$ | 0.93364 | 0.88740 | 0.98800 | 0.95600 |
| $\alpha_1$ | 1.70925 | 0.63980 | 0.01030 | 0.00614 |
| $\alpha_2$ | 0.67025 | -0.04340 | 0.01430 | 0.01390 |
| $\alpha_3$ | 0.67931 | 0.29480 | -0.00571 | -0.00386 |
| $\alpha_4$ | -0.17950 | -0.45920 | 0.02010 | 0.04190 |
| $\delta_1$ | 0.22184 | 0.05270 | 0.00005 | 0.00001 |
| $\delta_2$ | 0.19571 | 0.07520 | 0.00014 | 0.00003 |
| $\delta_3$ | 0.01924 | 0.01480 | 0.00049 | 0.00028 |
| $\delta_4$ | 0.04708 | 0.07570 | 0.00005 | 0.00010 |

**Table S1.5.** MAR estimates for the replicate LV dataset in Figs. 1 and S4.

|  | MAR without<br>transformation | MAR with log<br>transformation | MAR with<br>smoothing | MAR with log<br>transformation<br>and smoothing |
| --- | --- | --- | --- | --- |
| $\beta_{11}$ | 0.77800 | 0.76500 | 1.01000 | 1.07000 |
| $\beta_{21}$ | 0.07760 | 0.21000 | 0.00367 | 0.07720 |
| $\beta_{31}$ | -0.08920 | -0.31200 | -0.01630 | -0.10700 |
| $\beta_{41}$ | 0.09140 | 0.13400 | 0.23200 | 0.38400 |
| $\beta_{12}$ | 0.13500 | 0.11600 | -0.00781 | -0.00292 |
| $\beta_{22}$ | 0.87500 | 0.80500 | 0.93600 | 0.88300 |
| $\beta_{32}$ | 0.14900 | 0.22500 | 0.10400 | 0.14800 |
| $\beta_{42}$ | -0.00518 | 0.17400 | -0.08900 | 0.06510 |
| $\beta_{13}$ | 0.24800 | 0.22200 | -0.00982 | -0.00328 |
| $\beta_{23}$ | -0.20400 | -0.18300 | -0.10300 | -0.05940 |
| $\beta_{33}$ | 0.94400 | 1.13000 | 0.86500 | 0.98300 |
| $\beta_{43}$ | 0.01090 | 0.17500 | -0.02880 | 0.04240 |
| $\beta_{14}$ | -0.02600 | -0.01100 | -0.02170 | -0.00991 |
| $\beta_{24}$ | -0.00159 | -0.00250 | -0.00530 | -0.00583 |
| $\beta_{34}$ | -0.00022 | -0.00299 | 0.00004 | -0.00317 |
| $\beta_{44}$ | 0.97200 | 0.93000 | 0.96400 | 0.93000 |
| $\alpha_1$ | 0.02400 | 0.01470 | 0.00925 | 0.00610 |
| $\alpha_2$ | 0.02610 | 0.03690 | 0.01980 | 0.02190 |
| $\alpha_3$ | -0.02240 | -0.01470 | -0.00880 | -0.00573 |
| $\alpha_4$ | 0.02160 | 0.10700 | 0.02500 | 0.09300 |
| $\delta_1$ | 0.00557 | 0.00089 | 0.00008 | 0.00001 |
| $\delta_2$ | 0.00243 | 0.00073 | 0.00037 | 0.00009 |
| $\delta_3$ | 0.00001 | 0.00004 | 0.00024 | 0.00043 |
| $\delta_4$ | 0.00559 | 0.00267 | 0.00029 | 0.00022 |

**Table S2.1.** MAR estimates for the noisy dataset in Fig. S3.

|  | MAR without<br>transformation | MAR with log<br>transformation | MAR with<br>smoothing | MAR with log<br>transformation<br>and smoothing |
| --- | --- | --- | --- | --- |
| $\beta_{11}$ | 0.23009 | 0.01769 | 0.90100 | 0.93300 |
| $\beta_{21}$ | 0.11849 | 0.16822 | -0.18000 | -0.10300 |
| $\beta_{31}$ | -0.18449 | -0.35186 | -0.26400 | -0.61400 |
| $\beta_{41}$ | 0.14318 | 2.18894 | 0.40800 | 1.05000 |
| $\beta_{12}$ | 0.34796 | 0.23560 | 0.03870 | 0.01540 |
| $\beta_{22}$ | 0.84182 | 0.75617 | 1.05000 | 0.94800 |
| $\beta_{32}$ | 0.19214 | 0.17605 | 0.18600 | 0.20600 |
| $\beta_{42}$ | -0.03130 | 0.01950 | -0.15300 | -0.06350 |
| $\beta_{13}$ | 0.21488 | 0.18478 | -0.00110 | 0.00908 |
| $\beta_{23}$ | -0.04121 | 0.02412 | -0.10500 | -0.04810 |
| $\beta_{33}$ | 0.78353 | 0.91531 | 0.87900 | 1.02000 |
| $\beta_{43}$ | 0.01040 | 1.00320 | -0.02870 | 0.05100 |
| $\beta_{14}$ | 0.03493 | 0.01754 | -0.00471 | -0.00229 |
| $\beta_{24}$ | -0.00776 | 0.00115 | -0.00050 | -0.00170 |
| $\beta_{34}$ | -0.00548 | 0.00382 | 0.01160 | 0.01240 |
| $\beta_{44}$ | 0.97027 | 0.81890 | 0.95100 | 0.91800 |
| $\alpha_1$ | -0.00061 | 0.00073 | 0.01020 | 0.00586 |
| $\alpha_2$ | 0.02045 | 0.03086 | 0.02000 | 0.02130 |
| $\alpha_3$ | -0.01764 | -0.01187 | -0.00431 | -0.00279 |
| $\alpha_4$ | 0.02229 | 0.04416 | 0.02100 | 0.06150 |
| $\delta_1$ | 0.01953 | 0.00318 | 0.00001 | 0.00000 |
| $\delta_2$ | 0.01746 | 0.01414 | 0.00002 | 0.00000 |
| $\delta_3$ | 0.00158 | 0.00121 | 0.00002 | 0.00004 |
| $\delta_4$ | 0.00363 | 0.39522 | 0.00038 | 0.00007 |

**Table S2.2.** MAR estimates for the replicate dataset in Fig. S3.

|  | MAR without<br>transformation | MAR with log<br>transformation | MAR with<br>smoothing | MAR with log<br>transformation<br>and smoothing |
| --- | --- | --- | --- | --- |
| $\beta_{11}$ | 0.83000 | 0.81000 | 0.88600 | 0.87400 |
| $\beta_{21}$ | -0.06680 | 0.02220 | -0.05350 | 0.04420 |
| $\beta_{31}$ | -0.12600 | -0.35400 | -0.09890 | -0.39000 |
| $\beta_{41}$ | 0.23600 | 0.37800 | 0.23500 | 0.41700 |
| $\beta_{12}$ | 0.07710 | 0.04860 | 0.05470 | 0.04280 |
| $\beta_{22}$ | 0.97800 | 0.89700 | 0.97600 | 0.90000 |
| $\beta_{32}$ | 0.16200 | 0.19000 | 0.13100 | 0.19100 |
| $\beta_{42}$ | -0.07360 | 0.10500 | -0.08250 | 0.08650 |
| $\beta_{13}$ | 0.12900 | 0.11100 | 0.05310 | 0.06250 |
| $\beta_{23}$ | -0.06320 | -0.03200 | -0.09610 | -0.06310 |
| $\beta_{33}$ | 0.87400 | 1.02000 | 0.89000 | 1.06000 |
| $\beta_{43}$ | -0.04980 | 0.06360 | -0.02580 | 0.08010 |
| $\beta_{14}$ | -0.00609 | -0.00121 | -0.00871 | -0.00304 |
| $\beta_{24}$ | -0.00592 | -0.00445 | -0.00621 | -0.00488 |
| $\beta_{34}$ | -0.00154 | 0.00406 | -0.00073 | 0.00508 |
| $\beta_{44}$ | 0.96100 | 0.92600 | 0.96300 | 0.92500 |
| $\alpha_1$ | 0.02250 | 0.01380 | 0.01240 | 0.00749 |
| $\alpha_2$ | 0.03120 | 0.04080 | 0.02220 | 0.02350 |
| $\alpha_3$ | -0.02690 | -0.01750 | -0.00777 | -0.00512 |
| $\alpha_4$ | 0.01850 | 0.10700 | 0.02520 | 0.09450 |
| $\delta_1$ | 0.00052 | 0.00004 | 0.00007 | 0.00001 |
| $\delta_2$ | 0.00029 | 0.00007 | 0.00004 | 0.00001 |
| $\delta_3$ | 0.00002 | 0.00000 | 0.00002 | 0.00001 |
| $\delta_4$ | 0.00219 | 0.00114 | 0.00039 | 0.00014 |

**Table S3.1.**
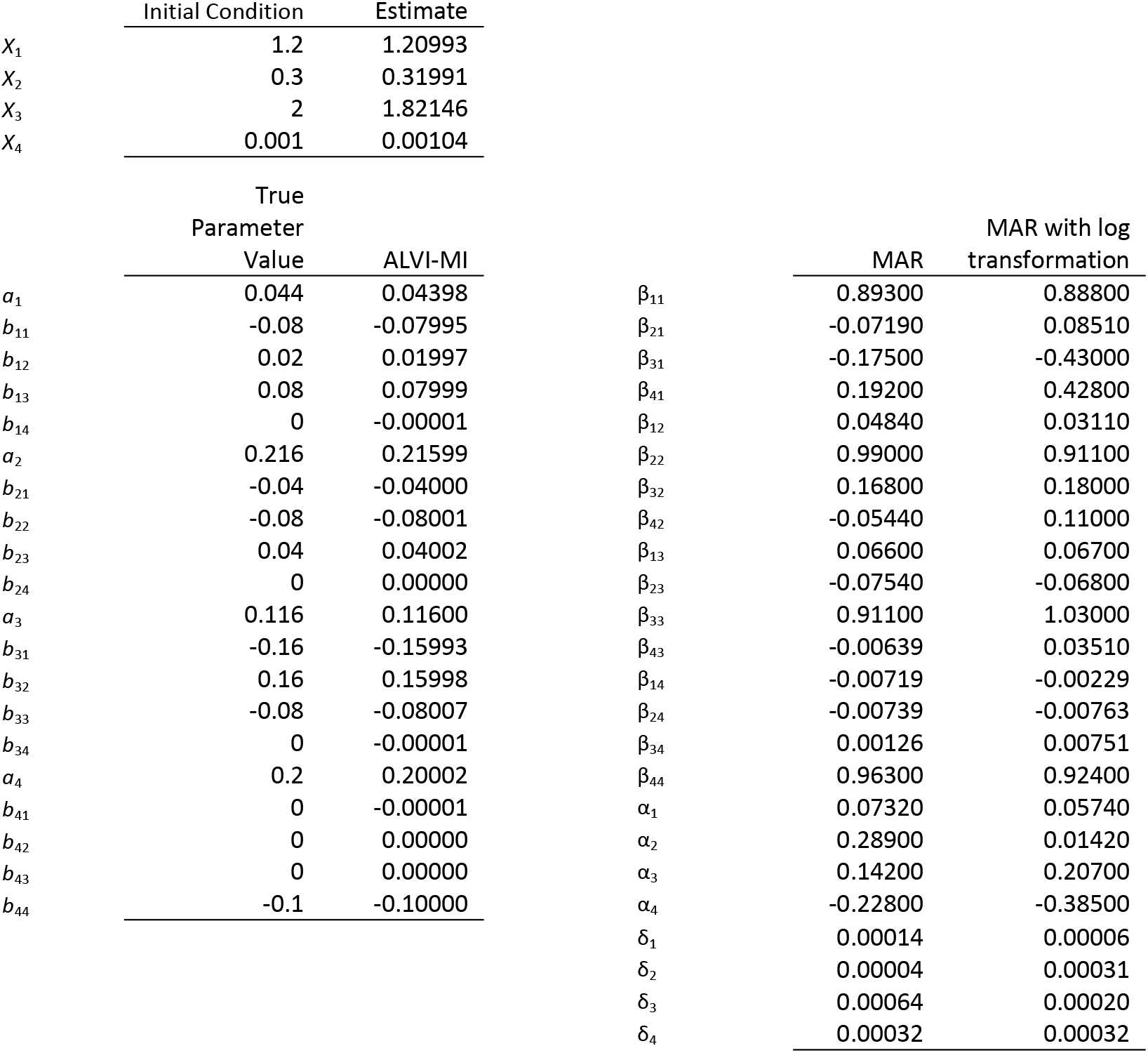
Initial conditions, parameter values and estimates for a four-variable LV system that converges to a stable steady state, as presented in Fig. S5.

|  | Initial Condition | Estimate |  |  |  |
| --- | --- | --- | --- | --- | --- |
| $X_1$ | 1.2 | 1.20993 | | | |
| $X_2$ | 0.3 | 0.31991 | | | |
| $X_3$ | 2 | 1.82146 | | | |
| $X_4$ | 0.001 | 0.00104 | | | |
|  | True |  |  |  |  |
|  | Parameter |  |  |  |  |
|  | Value | ALVI-MI |  | MAR | MAR with log transformation |
| $a_1$ | 0.044 | 0.04398 | $\beta_{11}$ | 0.89300 | 0.88800 |
| $b_{11}$ | -0.08 | -0.07995 | $\beta_{21}$ | -0.07190 | 0.08510 |
| $b_{12}$ | 0.02 | 0.01997 | $\beta_{31}$ | -0.17500 | -0.43000 |
| $b_{13}$ | 0.08 | 0.07999 | $\beta_{41}$ | 0.19200 | 0.42800 |
| $b_{14}$ | 0 | -0.00001 | $\beta_{12}$ | 0.04840 | 0.03110 |
| $a_2$ | 0.216 | 0.21599 | $\beta_{22}$ | 0.99000 | 0.91100 |
| $b_{21}$ | -0.04 | -0.04000 | $\beta_{32}$ | 0.16800 | 0.18000 |
| $b_{22}$ | -0.08 | -0.08001 | $\beta_{42}$ | -0.05440 | 0.11000 |
| $b_{23}$ | 0.04 | 0.04002 | $\beta_{13}$ | 0.06600 | 0.06700 |
| $b_{24}$ | 0 | 0.00000 | $\beta_{23}$ | -0.07540 | -0.06800 |
| $a_3$ | 0.116 | 0.11600 | $\beta_{33}$ | 0.91100 | 1.03000 |
| $b_{31}$ | -0.16 | -0.15993 | $\beta_{43}$ | -0.00639 | 0.03510 |
| $b_{32}$ | 0.16 | 0.15998 | $\beta_{14}$ | -0.00719 | -0.00229 |
| $b_{33}$ | -0.08 | -0.08007 | $\beta_{24}$ | -0.00739 | -0.00763 |
| $b_{34}$ | 0 | -0.00001 | $\beta_{34}$ | 0.00126 | 0.00751 |
| $a_4$ | 0.2 | 0.20002 | $\beta_{44}$ | 0.96300 | 0.92400 |
| $b_{41}$ | 0 | -0.00001 | $\alpha_1$ | 0.07320 | 0.05740 |
| $b_{42}$ | 0 | 0.00000 | $\alpha_2$ | 0.28900 | 0.01420 |
| $b_{43}$ | 0 | 0.00000 | $\alpha_3$ | 0.14200 | 0.20700 |
| $b_{44}$ | -0.1 | -0.10000 | $\alpha_4$ | -0.22800 | -0.38500 |
| | | | $\delta_1$ | 0.00014 | 0.00006 |
| | | | $\delta_2$ | 0.00004 | 0.00031 |
| | | | $\delta_3$ | 0.00064 | 0.00020 |
| | | | $\delta_4$ | 0.00032 | 0.00032 |

**Table S3.2.** Initial conditions, parameter values and estimates for a four-variable LV system exhibiting damped oscillations as presented in Fig. S5.

|  | Initial Condition | Estimate |  |  |  |
| --- | --- | --- | --- | --- | --- |
| $X_1$ | 0.3 | 0.3 | | | |
| $X_2$ | 0.3 | 0.3 | | | |
| $X_3$ | 0.4 | 0.4 | | | |
| $X_4$ | 0.6 | 0.6 | | | |
|  | True |  |  |  |  |
|  | Parameter |  |  |  |  |
|  | Value | ALVI-MI |  | MAR | MAR with log transformation |
| $a_1$ | 0.3 | 0.30245 | $\beta_{11}$ | 1.05000 | 0.98800 |
| $b_{11}$ | -0.3 | -0.30328 | $\beta_{21}$ | 0.14700 | 0.01610 |
| $b_{12}$ | -0.27 | -0.27098 | $\beta_{31}$ | -0.25300 | -0.07880 |
| $b_{13}$ | -0.6 | -0.60467 | $\beta_{41}$ | -0.23100 | -0.05360 |
| $b_{14}$ | -0.045 | -0.04708 | $\beta_{12}$ | -0.08350 | -0.11600 |
| $a_2$ | 0.4 | 0.40299 | $\beta_{22}$ | 0.88800 | 0.92200 |
| $b_{21}$ | 0.2 | 0.19592 | $\beta_{32}$ | 0.04880 | -0.03410 |
| $b_{22}$ | -0.4 | -0.40118 | $\beta_{42}$ | -0.00228 | -0.07260 |
| $b_{23}$ | -0.4 | -0.40578 | $\beta_{13}$ | 0.03660 | -0.04410 |
| $b_{24}$ | -0.6 | -0.60250 | $\beta_{23}$ | -0.07340 | -0.06540 |
| $a_3$ | 0.7 | 0.71519 | $\beta_{33}$ | 0.71200 | 0.89700 |
| $b_{31}$ | -2.38 | -2.39901 | $\beta_{43}$ | -0.20500 | -0.02050 |
| $b_{32}$ | 0.35 | 0.34344 | $\beta_{14}$ | -0.10400 | -0.01220 |
| $b_{33}$ | -2.8 | -2.82806 | $\beta_{24}$ | -0.12800 | -0.03300 |
| $b_{34}$ | 0.35 | 0.33650 | $\beta_{34}$ | 0.03690 | -0.14100 |
| $a_4$ | 0.6 | 0.60791 | $\beta_{44}$ | 0.96100 | 0.88100 |
| $b_{41}$ | -0.96 | -0.97003 | $\alpha_1$ | 0.04710 | -0.22200 |
| $b_{42}$ | -0.24 | -0.24338 | $\alpha_2$ | 0.08150 | -0.17700 |
| $b_{43}$ | -0.96 | -0.97466 | $\alpha_3$ | 0.06750 | -0.56600 |
| $b_{44}$ | -0.6 | -0.60704 | $\alpha_4$ | 0.08660 | -0.38800 |
| | | | $\delta_1$ | 0.00000 | 0.00004 |
| | | | $\delta_2$ | 0.00002 | 0.00016 |
| | | | $\delta_3$ | 0.00015 | 0.00271 |
| | | | $\delta_4$ | 0.00015 | 0.00065 |

**Table S3.3.** Initial conditions, parameter values and estimates for a four-variable LV system displaying initially erratic oscillations, but converging to a limit cycle as presented in Fig. S5.

|  | Initial Condition | Estimate |  |  |  |
| --- | --- | --- | --- | --- | --- |
| $X_1$ | 0.3 | 0.3 | | | |
| $X_2$ | 0.3 | 0.3 | | | |
| $X_3$ | 0.4 | 0.4 | | | |
| $X_4$ | 0.6 | 0.6 | | | |
|  | True<br>Parameter<br>Value | ALVI-MI |  | MAR | MAR with log<br>transformation |
| $a_1$ | 1 | 0.98743 | $\beta_{11}$ | 0.87600 | 0.94673 |
| $b_{11}$ | -1 | -0.99101 | $\beta_{21}$ | 0.26700 | 0.19269 |
| $b_{12}$ | -1.09 | -1.08083 | $\beta_{31}$ | -0.11100 | -0.77772 |
| $b_{13}$ | -1.52 | -1.51341 | $\beta_{41}$ | -0.17600 | -0.18648 |
| $b_{14}$ | 0 | 0.01346 | $\beta_{12}$ | -0.13600 | -0.11408 |
| $a_2$ | 0.72 | 0.72230 | $\beta_{22}$ | 0.90900 | 0.88847 |
| $b_{21}$ | 0 | -0.00274 | $\beta_{32}$ | 0.03980 | 0.59990 |
| $b_{22}$ | -0.72 | -0.72223 | $\beta_{42}$ | 0.06220 | 0.11789 |
| $b_{23}$ | -0.3168 | -0.31966 | $\beta_{13}$ | -0.44100 | 0.00528 |
| $b_{24}$ | -0.9792 | -0.97942 | $\beta_{23}$ | 0.07280 | 0.01411 |
| $a_3$ | 1.53 | 1.53466 | $\beta_{33}$ | 0.86500 | 0.93625 |
| $b_{31}$ | -3.672 | -3.67283 | $\beta_{43}$ | -0.03720 | -0.00153 |
| $b_{32}$ | 0 | -0.00071 | $\beta_{14}$ | 0.36500 | 0.00392 |
| $b_{33}$ | -1.53 | -1.52615 | $\beta_{24}$ | -0.19100 | -0.17211 |
| $b_{34}$ | -0.7191 | -0.73207 | $\beta_{34}$ | -0.02410 | 0.33444 |
| $a_4$ | 1.27 | 1.27361 | $\beta_{44}$ | 0.93800 | 0.97758 |
| $b_{41}$ | -1.5367 | -1.53884 | $\alpha_1$ | 0.02540 | -0.12418 |
| $b_{42}$ | -0.6477 | -0.64952 | $\alpha_2$ | 0.01980 | -0.00507 |
| $b_{43}$ | -0.4445 | -0.44519 | $\alpha_3$ | 0.03310 | -0.28038 |
| $b_{44}$ | -1.27 | -1.27581 | $\alpha_4$ | 0.04680 | -0.16530 |
| | | | $\delta_1$ | 0.00001 | 0.00057 |
| | | | $\delta_2$ | 0.00001 | 0.00012 |
| | | | $\delta_3$ | 0.00013 | 0.00112 |
| | | | $\delta_4$ | 0.00005 | 0.00022 |

**Table S3.4.**
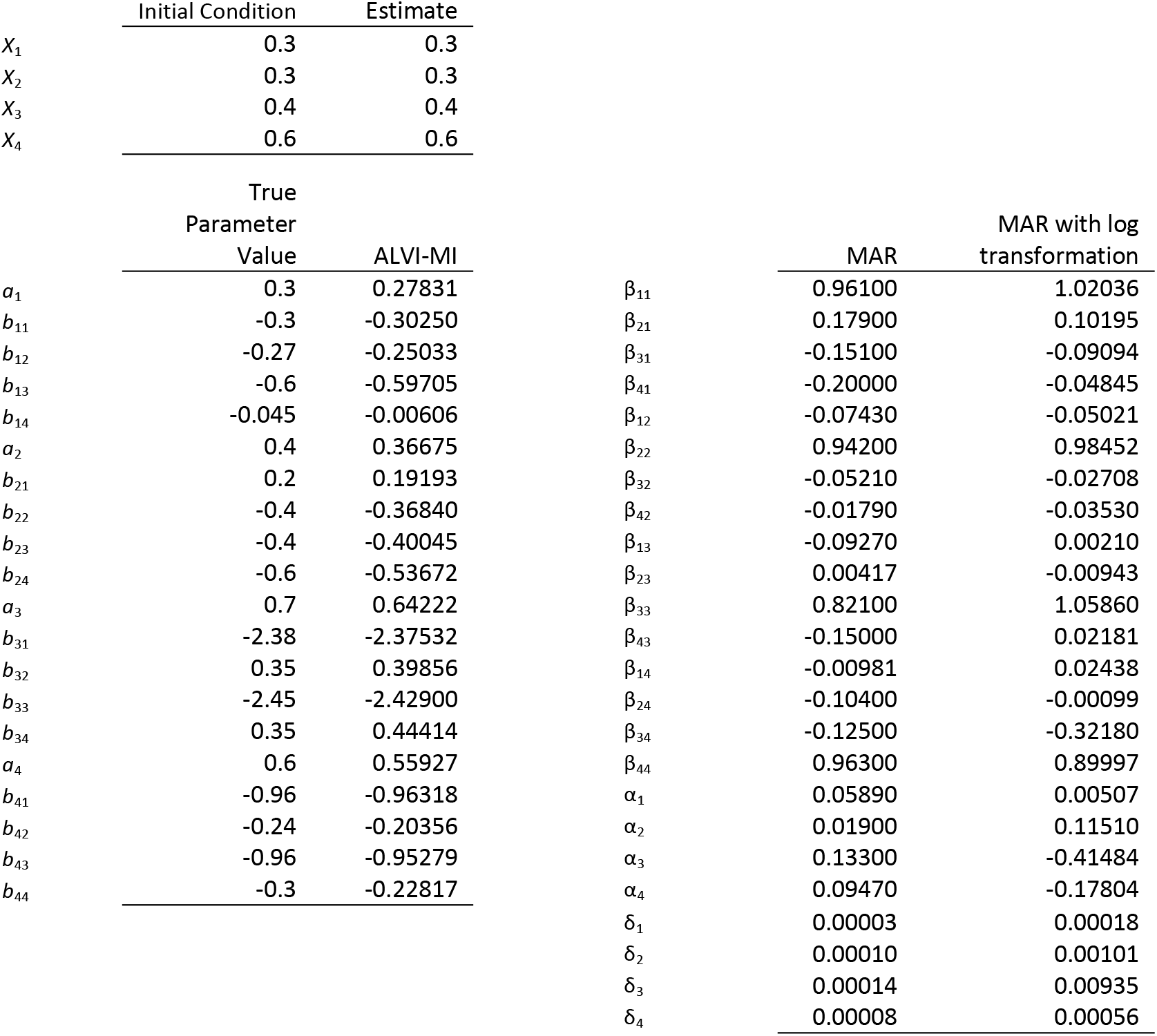
Initial conditions, parameter values and estimates for a four-variable LV system displaying damped oscillations as presented in Fig. S5.

|  | Initial Condition | Estimate |  |  |  |
| --- | --- | --- | --- | --- | --- |
| $X_1$ | 0.3 | 0.3 | | | |
| $X_2$ | 0.3 | 0.3 | | | |
| $X_3$ | 0.4 | 0.4 | | | |
| $X_4$ | 0.6 | 0.6 | | | |
|  | True |  |  |  |  |
|  | Parameter |  |  |  |  |
|  | Value | ALVI-MI |  | MAR | MAR with log transformation |
| $a_1$ | 0.3 | 0.27831 | $\beta_{11}$ | 0.96100 | 1.02036 |
| $b_{11}$ | -0.3 | -0.30250 | $\beta_{21}$ | 0.17900 | 0.10195 |
| $b_{12}$ | -0.27 | -0.25033 | $\beta_{31}$ | -0.15100 | -0.09094 |
| $b_{13}$ | -0.6 | -0.59705 | $\beta_{41}$ | -0.20000 | -0.04845 |
| $b_{14}$ | -0.045 | -0.00606 | $\beta_{12}$ | -0.07430 | -0.05021 |
| $a_2$ | 0.4 | 0.36675 | $\beta_{22}$ | 0.94200 | 0.98452 |
| $b_{21}$ | 0.2 | 0.19193 | $\beta_{32}$ | -0.05210 | -0.02708 |
| $b_{22}$ | -0.4 | -0.36840 | $\beta_{42}$ | -0.01790 | -0.03530 |
| $b_{23}$ | -0.4 | -0.40045 | $\beta_{13}$ | -0.09270 | 0.00210 |
| $b_{24}$ | -0.6 | -0.53672 | $\beta_{23}$ | 0.00417 | -0.00943 |
| $a_3$ | 0.7 | 0.64222 | $\beta_{33}$ | 0.82100 | 1.05860 |
| $b_{31}$ | -2.38 | -2.37532 | $\beta_{43}$ | -0.15000 | 0.02181 |
| $b_{32}$ | 0.35 | 0.39856 | $\beta_{14}$ | -0.00981 | 0.02438 |
| $b_{33}$ | -2.45 | -2.42900 | $\beta_{24}$ | -0.10400 | -0.00099 |
| $b_{34}$ | 0.35 | 0.44414 | $\beta_{34}$ | -0.12500 | -0.32180 |
| $a_4$ | 0.6 | 0.55927 | $\beta_{44}$ | 0.96300 | 0.89997 |
| $b_{41}$ | -0.96 | -0.96318 | $\alpha_1$ | 0.05890 | 0.00507 |
| $b_{42}$ | -0.24 | -0.20356 | $\alpha_2$ | 0.01900 | 0.11510 |
| $b_{43}$ | -0.96 | -0.95279 | $\alpha_3$ | 0.13300 | -0.41484 |
| $b_{44}$ | -0.3 | -0.22817 | $\alpha_4$ | 0.09470 | -0.17804 |
| | | | $\delta_1$ | 0.00003 | 0.00018 |
| | | | $\delta_2$ | 0.00010 | 0.00101 |
| | | | $\delta_3$ | 0.00014 | 0.00935 |
| | | | $\delta_4$ | 0.00008 | 0.00056 |

**Table S3.5.** Initial conditions, parameter values and estimates for a four-variable LV system displaying deterministic chaos (chaos 1) as presented in Fig. S5.

|  | Initial Condition | Estimate |  |  |  |
| --- | --- | --- | --- | --- | --- |
| $X_1$ | 0.3 | 0.3 | | | |
| $X_2$ | 0.3 | 0.3 | | | |
| $X_3$ | 0.4 | 0.4 | | | |
| $X_4$ | 0.6 | 0.6 | | | |
|  | True |  |  |  |  |
|  | Parameter |  |  |  |  |
|  | Value | ALVI-MI |  | MAR | MAR with log transformation |
| $a_1$ | 1 | 1.01561 | $\beta_{11}$ | 0.85600 | 1.00059 |
| $b_{11}$ | -1 | -1.01294 | $\beta_{21}$ | 0.26400 | 0.20355 |
| $b_{12}$ | -1.09 | -1.10110 | $\beta_{31}$ | -0.16900 | -0.79454 |
| $b_{13}$ | -1.52 | -1.52409 | $\beta_{41}$ | -0.21800 | -0.23777 |
| $b_{14}$ | 0 | -0.01741 | $\beta_{12}$ | -0.14300 | -0.10901 |
| $a_2$ | 0.72 | 0.72904 | $\beta_{22}$ | 0.88800 | 0.88053 |
| $b_{21}$ | 0 | -0.00777 | $\beta_{32}$ | 0.01640 | 0.59018 |
| $b_{22}$ | -0.72 | -0.72674 | $\beta_{42}$ | 0.02020 | 0.08015 |
| $b_{23}$ | -0.3168 | -0.32100 | $\beta_{13}$ | -0.47500 | -0.03240 |
| $b_{24}$ | -0.9792 | -0.98775 | $\beta_{23}$ | 0.10500 | 0.03330 |
| $a_3$ | 1.53 | 1.48293 | $\beta_{33}$ | 1.07000 | 0.76513 |
| $b_{31}$ | -3.5649 | -3.52631 | $\beta_{43}$ | 0.06420 | -0.03995 |
| $b_{32}$ | 0 | 0.03573 | $\beta_{14}$ | 0.38200 | 0.17765 |
| $b_{33}$ | -1.53 | -1.50898 | $\beta_{24}$ | -0.24000 | -0.23158 |
| $b_{34}$ | -0.7191 | -0.67335 | $\beta_{34}$ | -0.21500 | 0.90744 |
| $a_4$ | 1.27 | 1.25074 | $\beta_{44}$ | 0.80900 | 1.08027 |
| $b_{41}$ | -1.5367 | -1.52127 | $\alpha_1$ | 0.03510 | 0.01644 |
| $b_{42}$ | -0.6477 | -0.63310 | $\alpha_2$ | 0.04320 | -0.01131 |
| $b_{43}$ | -0.4445 | -0.43667 | $\alpha_3$ | 0.11000 | -0.11896 |
| $b_{44}$ | -1.27 | -1.25061 | $\alpha_4$ | 0.11500 | -0.24402 |
| | | | $\delta_1$ | 0.00002 | 0.00114 |
| | | | $\delta_2$ | 0.00002 | 0.00015 |
| | | | $\delta_3$ | 0.00022 | 0.00191 |
| | | | $\delta_4$ | 0.00006 | 0.00024 |

**Table S3.6.** Initial conditions, parameter values and estimates for a four-variable LV system displaying deterministic chaos (chaos 2) as presented in Fig. S5.

|  | Initial Condition | Estimate |  |  |  |
| --- | --- | --- | --- | --- | --- |
| $X_1$ | 0.3 | 0.3 | | | |
| $X_2$ | 0.3 | 0.3 | | | |
| $X_3$ | 0.4 | 0.4 | | | |
| $X_4$ | 0.6 | 0.6 | | | |
|  | True |  |  |  |  |
|  | Parameter |  |  |  |  |
|  | Value | ALVI-MI |  | MAR | MAR with log transformation |
| $a_1$ | 0.3 | 0.29277 | $\beta_{11}$ | 1.05000 | 0.97700 |
| $b_{11}$ | -0.3 | -0.25379 | $\beta_{21}$ | 0.00560 | 0.02030 |
| $b_{12}$ | -0.27 | -0.28051 | $\beta_{31}$ | -0.31600 | -0.08880 |
| $b_{13}$ | -0.6 | -0.55795 | $\beta_{41}$ | -0.23000 | -0.06400 |
| $b_{14}$ | -0.045 | -0.07045 | $\beta_{12}$ | -0.09830 | -0.01780 |
| $a_2$ | 0.4 | 0.39048 | $\beta_{22}$ | 0.99000 | 1.00000 |
| $b_{21}$ | 0.2 | 0.26661 | $\beta_{32}$ | 0.08310 | -0.02620 |
| $b_{22}$ | -0.4 | -0.41601 | $\beta_{42}$ | -0.01940 | -0.01610 |
| $b_{23}$ | -0.4 | -0.33988 | $\beta_{13}$ | -0.00541 | -0.06900 |
| $b_{24}$ | -0.6 | -0.63818 | $\beta_{23}$ | -0.12100 | -0.07290 |
| $a_3$ | 0.8 | 0.77659 | $\beta_{33}$ | 0.70200 | 0.87400 |
| $b_{31}$ | -2.38 | -2.23922 | $\beta_{43}$ | -0.18000 | -0.05050 |
| $b_{32}$ | 0.35 | 0.31928 | $\beta_{14}$ | -0.08040 | 0.05960 |
| $b_{33}$ | -2.45 | -2.32117 | $\beta_{24}$ | 0.00941 | -0.02020 |
| $b_{34}$ | 0.35 | 0.27470 | $\beta_{34}$ | 0.07400 | -0.14500 |
| $a_4$ | 0.6 | 0.58578 | $\beta_{44}$ | 0.97300 | 0.94100 |
| $b_{41}$ | -0.96 | -0.86962 | $\alpha_1$ | 0.05270 | -0.11500 |
| $b_{42}$ | -0.24 | -0.26048 | $\alpha_2$ | 0.02820 | -0.09870 |
| $b_{43}$ | -0.96000 | -0.87774 | $\alpha_3$ | 0.07760 | -0.53700 |
| $b_{44}$ | -0.30000 | -0.34969 | $\alpha_4$ | 0.10200 | -0.28000 |
| | | | $\delta_1$ | 0.00000 | 0.00006 |
| | | | $\delta_2$ | 0.00002 | 0.00023 |
| | | | $\delta_3$ | 0.00009 | 0.00115 |
| | | | $\delta_4$ | 0.00007 | 0.00025 |

**Table S4.1.**
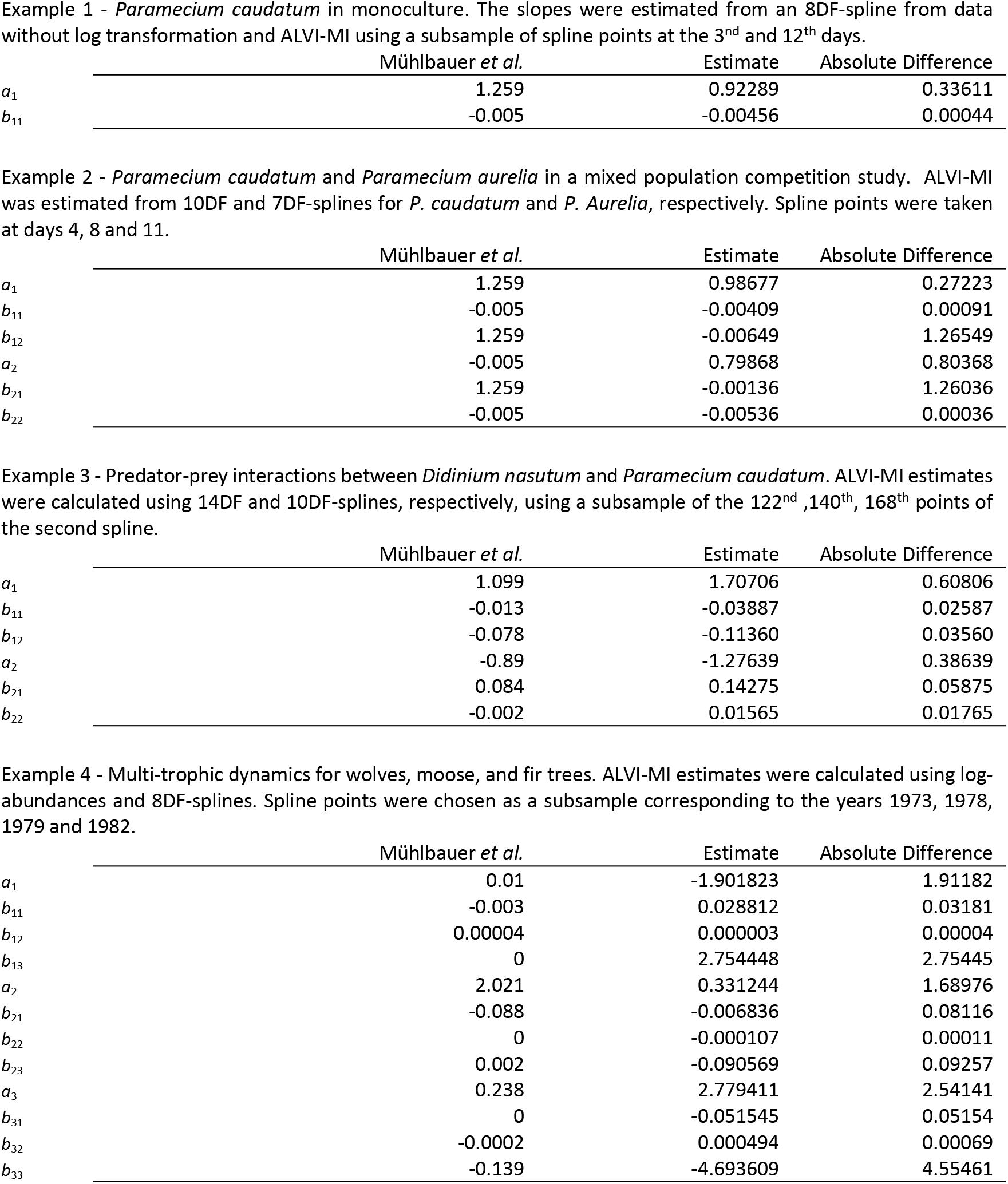
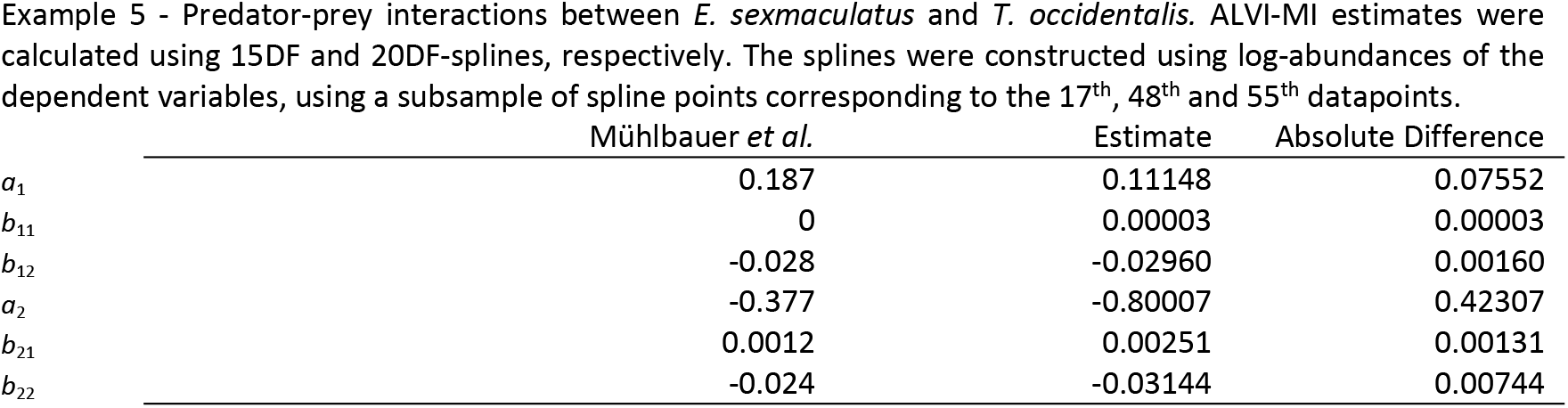
ALVI-MI estimates for five experimental data examples from (Mühlbauer et al., 2020). Data came from experiments described in (Gause, 1934), (McLaren & Peterson, 1994) and (Huffaker et al., 1963). See R package gauseR (Mühlbauer et al., 2020) for datasets “gause_1934_science_f02_03”, “gause_1934_book_f32”, “mclaren_1994_f03” and “huffaker_1963” for details on observations. Parameter estimates from Mühlbauer *et al*. can also be found in Table 2 in their paper.

**Table S4.2.** ALVI-LR estimates for five experimental data sets from (Mühlbauer et al., 2020). Data came from (Gause, 1934), (McLaren & Peterson, 1994) and (Huffaker et al., 1963) experiments. See R package gauseR (Mühlbauer et al., 2020) datasets “gause_1934_science_f02_03”, “gause_1934_book_f32”, “mclaren_1994_f03” and “huffaker_1963” for details on observations. Parameter estimates from Mühlbauer *et al*. can also be found in Table 2 of their paper.

|  | Mühlbauer *et al.* | Estimate | Absolute Difference |
| --- | --- | --- | --- |
| $a_1$ | 1.259 | 0.93948 | 0.31952 |
| $b_{11}$ | -0.005 | -0.00465 | 0.00035 |

Example 2 - *Paramecium caudatum* and *Paramecium aurelia* in mixed population, analyzed with 10DF and 7DF-splines
|  | Mühlbauer *et al.* | Estimate | Absolute Difference |
| --- | --- | --- | --- |
| $a_1$ | 1.259 | 0.85524 | 0.40376 |
| $b_{11}$ | -0.005 | -0.00289 | 0.00211 |
| $b_{12}$ | -0.008 | -0.00580 | 0.00220 |
| $a_2$ | 1.026 | 0.84423 | 0.18177 |
| $b_{21}$ | -0.002 | -0.00187 | 0.00013 |
| $b_{22}$ | -0.007 | -0.00553 | 0.00147 |

|  | Mühlbauer *et al.* | Estimate | Absolute Difference |
| --- | --- | --- | --- |
| $a_1$ | 1.099 | 0.45652 | 0.64248 |
| $b_{11}$ | -0.013 | 0.02117 | 0.03417 |
| $b_{12}$ | -0.078 | -0.11495 | 0.03695 |
| $a_2$ | -0.89 | -0.98922 | 0.09922 |
| $b_{21}$ | 0.084 | 0.16549 | 0.08149 |
| $b_{22}$ | -0.002 | -0.01146 | 0.00946 |

Example 4 - Multi-trophic dynamics for wolves, moose, and fir trees, analyzed with 28DF, 24DF and 28DF-splines
|  | Mühlbauer *et al.* | Estimate | Absolute Difference |
| --- | --- | --- | --- |
| $a_1$ | 0.01 | -0.06509 | 0.07509 |
| $b_{11}$ | -0.003 | 0.00164 | 0.00464 |
| $b_{12}$ | 0.00004 | 0.00007 | 0.00003 |
| $b_{13}$ | 0 | -0.13411 | 0.13411 |
| $a_2$ | 2.021 | 0.20754 | 1.81346 |
| $b_{21}$ | -0.088 | -0.00483 | 0.08317 |
| $b_{22}$ | 0 | -0.00009 | 0.00009 |
| $b_{23}$ | 0.002 | 0.06010 | 0.05810 |
| $a_3$ | 0.238 | -0.08580 | 0.32380 |
| $b_{31}$ | 0 | 0.00343 | 0.00343 |
| $b_{32}$ | -0.0002 | -0.00020 | 0.00000 |
| $b_{33}$ | -0.139 | 0.43352 | 0.57252 |

|  | Mühlbauer *et al.* | Estimate | Absolute Difference |
| --- | --- | --- | --- |
| $a_1$ | 0.344 | 0.03525 | 0.30875 |
| $b_{11}$ | 0 | 0.00038 | 0.00038 |
| $b_{12}$ | -0.059 | -0.03619 | 0.02281 |
| $a_2$ | -0.236 | -0.44687 | 0.21087 |
| $b_{21}$ | 0.0005 | 0.00159 | 0.00109 |
| $b_{22}$ | 0 | -0.03540 | 0.03540 |

**Table S4.3.** MAR estimates for five experimental data sets from (Mühlbauer et al., 2020). Data came from (Gause, 1934), (McLaren & Peterson, 1994) and (Huffaker et al., 1963) experiments. See R package gauseR (Mühlbauer et al., 2020) for datasets “gause_1934_science_f02_03”, “gause_1934_book_f32”, “mclaren_1994_f03” and “huffaker_1963” for details on observations. Parameter estimates from Mühlbauer *et al*. can also be found in Table 2 on their paper.

|  | MAR | MAR log transformation | MAR with data smoothing | MAR log transformation with smoothing |
| --- | --- | --- | --- | --- |
| $\beta_1$ | 0.90000 | 0.85550 | 0.90400 | 0.74718 |
| $\alpha_1$ | 9.93000 | 0.15590 | 10.22600 | 0.21106 |
| $\delta_1$ | 287.57000 | 0.06920 | 126.15300 | 0.00525 |

Example 2 - *Paramecium caudatum* and *Paramecium aurelia* in coculture
|  | MAR | MAR log transformation | MAR with data smoothing | MAR log transformation with smoothing |
| --- | --- | --- | --- | --- |
| $\beta_{11}$ | 0.81800 | 0.73590 | 0.91500 | 0.98309 |
| $\beta_{21}$ | 0.07170 | 0.03240 | 0.11600 | 0.03625 |
| $\beta_{12}$ | -0.16030 | -0.07020 | -0.16900 | -0.26372 |
| $\beta_{22}$ | 0.82140 | 0.71450 | 0.87700 | 0.73153 |
| $\alpha_1$ | 27.54890 | 1.38260 | 0.91000 | 0.07810 |
| $\alpha_2$ | 20.30740 | 1.28150 | 5.82700 | 0.15430 |
| $\delta_1$ | 262.75870 | 0.17190 | 113.99000 | 0.03530 |
| $\delta_2$ | 253.59620 | 0.03470 | 11.12100 | 0.00202 |

Example 3 - Predator-prey interactions between *Didinium nasutum* and *Paramecium caudatum*
|  | MAR | MAR log transformation | MAR with data smoothing | MAR log transformation with smoothing |
| --- | --- | --- | --- | --- |
| $\beta_{11}$ | 0.52400 | 0.80250 | 0.37800 | 0.86900 |
| $\beta_{21}$ | 0.57700 | 0.10510 | 0.52100 | 0.18700 |
| $\beta_{12}$ | -0.57200 | -0.21950 | -0.45100 | -0.25400 |
| $\beta_{22}$ | 0.63300 | 0.59490 | 0.69100 | 0.76500 |
| $\alpha_1$ | 12.93300 | -1.11100 | 0.40300 | -0.75400 |
| $\alpha_2$ | -2.76100 | 0.06450 | 0.23900 | 0.70000 |
| $\delta_1$ | 94.22700 | 10.97630 | 140.72600 | 7.15700 |
| $\delta_2$ | 27.95400 | 23.00840 | 83.62600 | 9.82300 |

Example 4 - Multi-trophic dynamics for wolves, moose, and fir trees
|  | MAR | MAR log<br>transformation | MAR with data<br>smoothing | MAR log<br>transformation with<br>smoothing |
| --- | --- | --- | --- | --- |
| $\beta_{11}$ | 0.62900 | 0.69390 | 1.13000 | 1.17360 |
| $\beta_{21}$ | -4.15000 | -0.04540 | -3.48000 | -0.04426 |
| $\beta_{31}$ | -0.00083 | -0.02210 | 0.00137 | 0.07236 |
| $\beta_{12}$ | 0.00517 | 0.07570 | 0.00148 | 0.01665 |
| $\beta_{22}$ | 0.84800 | 0.86270 | 0.90300 | 0.89080 |
| $\beta_{32}$ | -0.00007 | -0.12840 | -0.00009 | -0.13234 |
| $\beta_{13}$ | -27.30000 | -0.34650 | 10.70000 | 0.22128 |
| $\beta_{23}$ | 96.60000 | 0.08020 | 245.00000 | 0.13054 |
| $\beta_{33}$ | 0.89000 | 0.89160 | 1.11000 | 1.06497 |
| $\alpha_1$ | 15.90000 | 0.12600 | -0.35700 | -0.02613 |
| $\alpha_2$ | 241.00000 | 1.18230 | 27.80000 | 0.02986 |
| $\alpha_3$ | 0.14200 | 0.85670 | -0.00485 | -0.01065 |
| $\delta_1$ | 32.30000 | 0.06210 | 3.88000 | 0.00553 |
| $\delta_2$ | 15000.00000 | 0.01080 | 1290.00000 | 0.00127 |
| $\delta_3$ | 0.00382 | 0.02160 | 0.00098 | 0.00591 |

Example 5 - Predator-prey interactions between *E. sexmaculatus* and *T. occidentalis*
|  | MAR | MAR log<br>transformation | MAR with data<br>smoothing | MAR log<br>transformation with<br>smoothing |
| --- | --- | --- | --- | --- |
| $\beta_{11}$ | 1.01500 | 0.88120 | 1.11000 | 1.05163 |
| $\beta_{21}$ | 0.00900 | 0.53850 | 0.00718 | 0.47337 |
| $\beta_{12}$ | -17.79700 | -0.09960 | -16.00000 | -0.10626 |
| $\beta_{22}$ | 0.56200 | 0.75520 | 0.73200 | 0.86330 |
| $\alpha_1$ | 94.24600 | 0.80680 | 0.47700 | -0.00088 |
| $\alpha_2$ | -1.28500 | -2.86420 | -0.02580 | -0.01771 |
| $\delta_1$ | 15440.27300 | 0.13470 | 1360.00000 | 0.02078 |
| $\delta_2$ | 5.55100 | 0.33790 | 1.93000 | 0.08104 |

**Table S5.1.** Synthetic MAR data.

| $t$ | $X_1$ | $X_2$ | $X_3$ | $X_4$ |
| --- | --- | --- | --- | --- |
| 1 | 5 | 10 | 15 | 20 |
| 2 | 5.62418 | 35.2103 | 21.4125 | 10.3696 |
| 3 | 13.077 | 68.6603 | 16.2175 | 6.98986 |
| 4 | 30.8946 | 57.2944 | 7.75681 | 8.13234 |
| 5 | 42.7475 | 27.4855 | 3.71568 | 13.9927 |
| 6 | 31.3421 | 13.2593 | 2.63367 | 24.3619 |
| 7 | 16.5916 | 9.92497 | 3.02139 | 31.2622 |
| 8 | 9.868 | 12.4469 | 4.56419 | 27.4996 |
| 9 | 8.72248 | 20.3998 | 6.5998 | 19.5587 |
| 10 | 11.2768 | 30.657 | 7.33895 | 14.4998 |
| 11 | 16.7965 | 33.724 | 6.20514 | 13.314 |
| 12 | 21.9882 | 27.4413 | 4.66674 | 15.2345 |
| 13 | 22.1248 | 19.9474 | 3.79027 | 19.1227 |
| 14 | 18.0405 | 16.0646 | 3.69825 | 22.5386 |
| 15 | 14.0932 | 15.9557 | 4.22409 | 22.9275 |
| 16 | 12.3388 | 18.7495 | 5.03802 | 20.5772 |
| 17 | 12.7869 | 22.8183 | 5.59201 | 17.8653 |
| 18 | 14.8002 | 25.3774 | 5.51118 | 16.4484 |
| 19 | 17.1129 | 24.6804 | 4.99159 | 16.6605 |
| 20 | 18.1251 | 21.9731 | 4.48974 | 18.0462 |
| 21 | 17.3171 | 19.5701 | 4.27918 | 19.6396 |
| 22 | 15.6847 | 18.6862 | 4.39198 | 20.3941 |
| 23 | 14.456 | 19.3693 | 4.70288 | 19.9587 |
| 24 | 14.1822 | 20.9554 | 4.99681 | 18.8896 |
| 25 | 14.7787 | 22.3651 | 5.08888 | 17.9987 |
| 26 | 15.7517 | 22.7146 | 4.95911 | 17.7463 |
| 27 | 16.4482 | 21.9861 | 4.74141 | 18.1246 |
| 28 | 16.4701 | 20.8982 | 4.58774 | 18.7896 |
| 29 | 15.9468 | 20.1878 | 4.56995 | 19.2878 |
| 30 | 15.3325 | 20.1628 | 4.6678 | 19.3417 |
| 31 | 15.0101 | 20.6857 | 4.80099 | 19.0122 |

**Table S5.2.** Synthetic MAR data with added noise (noisy MAR)

| $t$ | $X_1$ | $X_2$ | $X_3$ | $X_4$ |
| --- | --- | --- | --- | --- |
| 1 | 0.7537 | 5.7537 | 10.7537 | 15.7537 |
| 2 | 3.6789 | 33.265 | 19.4672 | 8.42431 |
| 3 | 22.0484 | 77.6318 | 25.189 | 15.9613 |
| 4 | 27.8408 | 54.24056 | 4.70298 | 5.0785 |
| 5 | 44.2644 | 29.0024 | 5.23259 | 15.5096 |
| 6 | 27.0183 | 8.93558 | 0.005 | 20.0381 |
| 7 | 12.285 | 5.61827 | 0.005 | 26.9555 |
| 8 | 8.64674 | 11.2257 | 3.34294 | 26.2784 |
| 9 | 13.7228 | 25.4001 | 11.6001 | 24.559 |
| 10 | 9.55584 | 28.9361 | 5.61798 | 12.7788 |
| 11 | 19.8884 | 36.8159 | 9.29705 | 16.406 |
| 12 | 16.6399 | 22.093 | 0.005 | 9.88615 |
| 13 | 20.6563 | 18.479 | 2.32184 | 17.6543 |
| 14 | 15.1923 | 13.2165 | 0.85012 | 19.6905 |
| 15 | 12.7385 | 14.601 | 2.86941 | 21.5728 |
| 16 | 16.8878 | 23.2986 | 9.58707 | 25.1263 |
| 17 | 12.3928 | 22.4243 | 5.19791 | 17.4712 |
| 18 | 15.4333 | 26.0105 | 6.14431 | 17.0815 |
| 19 | 18.0468 | 25.6143 | 5.92554 | 17.5945 |
| 20 | 15.8709 | 19.7188 | 2.23545 | 15.7919 |
| 21 | 12.0743 | 14.3274 | 0.005 | 14.3969 |
| 22 | 14.4706 | 17.472 | 3.17787 | 19.18 |
| 23 | 14.9269 | 19.8403 | 5.17384 | 20.4297 |
| 24 | 17.9169 | 24.6901 | 8.73157 | 22.6243 |
| 25 | 13.8412 | 21.4276 | 4.15137 | 17.0612 |
| 26 | 14.6419 | 21.6048 | 3.8493 | 16.6365 |
| 27 | 22.3732 | 27.9111 | 10.6664 | 24.0496 |
| 28 | 13.6319 | 18.06 | 1.74954 | 15.9514 |
| 29 | 17.5022 | 21.7432 | 6.12531 | 20.8432 |
| 30 | 13.1904 | 18.0207 | 2.52572 | 17.1996 |
| 31 | 16.5325 | 22.2082 | 6.32346 | 20.5347 |

**Table S5.3.**
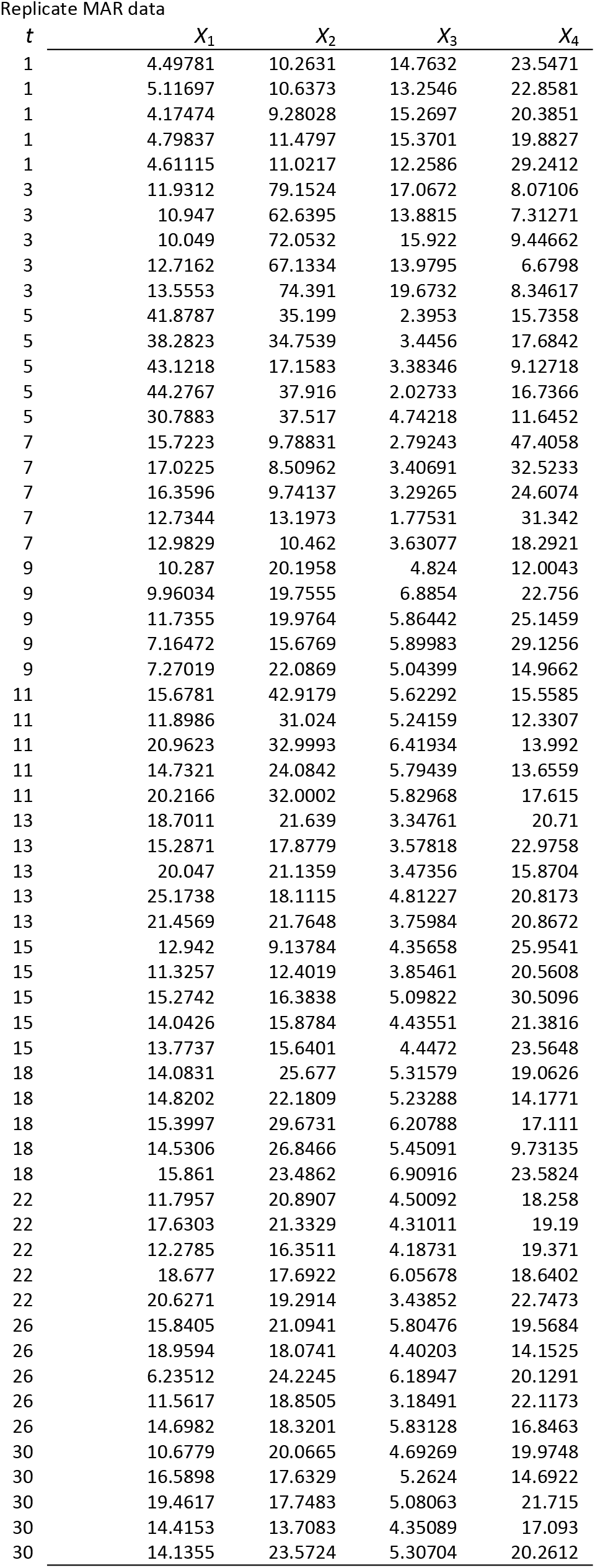
Synthetic MAR data with added noise replicates (replicate MAR)

| Replicate MAR data |  |  |  |  |
| --- | --- | --- | --- | --- |
| $t$ | $X_1$ | $X_2$ | $X_3$ | $X_4$ |
| 1 | 4.49781 | 10.2631 | 14.7632 | 23.5471 |
| 1 | 5.11697 | 10.6373 | 13.2546 | 22.8581 |
| 1 | 4.17474 | 9.28028 | 15.2697 | 20.3851 |
| 1 | 4.79837 | 11.4797 | 15.3701 | 19.8827 |
| 1 | 4.61115 | 11.0217 | 12.2586 | 29.2412 |
| 3 | 11.9312 | 79.1524 | 17.0672 | 8.07106 |
| 3 | 10.947 | 62.6395 | 13.8815 | 7.31271 |
| 3 | 10.049 | 72.0532 | 15.922 | 9.44662 |
| 3 | 12.7162 | 67.1334 | 13.9795 | 6.6798 |
| 3 | 13.5553 | 74.391 | 19.6732 | 8.34617 |
| 5 | 41.8787 | 35.199 | 2.3953 | 15.7358 |
| 5 | 38.2823 | 34.7539 | 3.4456 | 17.6842 |
| 5 | 43.1218 | 17.1583 | 3.38346 | 9.12718 |
| 5 | 44.2767 | 37.916 | 2.02733 | 16.7366 |
| 5 | 30.7883 | 37.517 | 4.74218 | 11.6452 |
| 7 | 15.7223 | 9.78831 | 2.79243 | 47.4058 |
| 7 | 17.0225 | 8.50962 | 3.40691 | 32.5233 |
| 7 | 16.3596 | 9.74137 | 3.29265 | 24.6074 |
| 7 | 12.7344 | 13.1973 | 1.77531 | 31.342 |
| 7 | 12.9829 | 10.462 | 3.63077 | 18.2921 |
| 9 | 10.287 | 20.1958 | 4.824 | 12.0043 |
| 9 | 9.96034 | 19.7555 | 6.8854 | 22.756 |
| 9 | 11.7355 | 19.9764 | 5.86442 | 25.1459 |
| 9 | 7.16472 | 15.6769 | 5.89983 | 29.1256 |
| 9 | 7.27019 | 22.0869 | 5.04399 | 14.9662 |
| 11 | 15.6781 | 42.9179 | 5.62292 | 15.5585 |
| 11 | 11.8986 | 31.024 | 5.24159 | 12.3307 |
| 11 | 20.9623 | 32.9993 | 6.41934 | 13.992 |
| 11 | 14.7321 | 24.0842 | 5.79439 | 13.6559 |
| 11 | 20.2166 | 32.0002 | 5.82968 | 17.615 |
| 13 | 18.7011 | 21.639 | 3.34761 | 20.71 |
| 13 | 15.2871 | 17.8779 | 3.57818 | 22.9758 |
| 13 | 20.047 | 21.1359 | 3.47356 | 15.8704 |
| 13 | 25.1738 | 18.1115 | 4.81227 | 20.8173 |
| 13 | 21.4569 | 21.7648 | 3.75984 | 20.8672 |
| 15 | 12.942 | 9.13784 | 4.35658 | 25.9541 |
| 15 | 11.3257 | 12.4019 | 3.85461 | 20.5608 |
| 15 | 15.2742 | 16.3838 | 5.09822 | 30.5096 |
| 15 | 14.0426 | 15.8784 | 4.43551 | 21.3816 |
| 15 | 13.7737 | 15.6401 | 4.4472 | 23.5648 |
| 18 | 14.0831 | 25.677 | 5.31579 | 19.0626 |
| 18 | 14.8202 | 22.1809 | 5.23288 | 14.1771 |
| 18 | 15.3997 | 29.6731 | 6.20788 | 17.111 |
| 18 | 14.5306 | 26.8466 | 5.45091 | 9.73135 |
| 18 | 15.861 | 23.4862 | 6.90916 | 23.5824 |
| 22 | 11.7957 | 20.8907 | 4.50092 | 18.258 |
| 22 | 17.6303 | 21.3329 | 4.31011 | 19.19 |
| 22 | 12.2785 | 16.3511 | 4.18731 | 19.371 |
| 22 | 18.677 | 17.6922 | 6.05678 | 18.6402 |
| 22 | 20.6271 | 19.2914 | 3.43852 | 22.7473 |
| 26 | 15.8405 | 21.0941 | 5.80476 | 19.5684 |
| 26 | 18.9594 | 18.0741 | 4.40203 | 14.1525 |
| 26 | 6.23512 | 24.2245 | 6.18947 | 20.1291 |
| 26 | 11.5617 | 18.8505 | 3.18491 | 22.1173 |
| 26 | 14.6982 | 18.3201 | 5.83128 | 16.8463 |
| 30 | 10.6779 | 20.0665 | 4.69269 | 19.9748 |
| 30 | 16.5898 | 17.6329 | 5.2624 | 14.6922 |
| 30 | 19.4617 | 17.7483 | 5.08063 | 21.715 |
| 30 | 14.4153 | 13.7083 | 4.35089 | 17.093 |
| 30 | 14.1355 | 23.5724 | 5.30704 | 20.2612 |

**Table S5.4.** Initial conditions and ALVI-MI parameters estimates for the synthetic MAR data.

|  | Noise MAR | Replicate MAR |
| --- | --- | --- |
| Spline | 15DF-spline | 15DF-spline |
| Point subsample | $t = 2, 6, 15, 18, 26$ | $t = 1, 3, 11, 13, 15$ |
|  | Initial condition | Initial condition |
| $X_1$ | 0.814535 | 4.639806 |
| $X_2$ | 7.333485 | 10.536398 |
| $X_3$ | 12.89041 | 14.183246 |
| $X_4$ | 14.301336 | 23.182858 |
| Parameter | Estimate | Estimate |
| $a_1$ | 51.4374203 | 1.986927 |
| $b_{11}$ | -0.7981952 | -0.029770 |
| $b_{12}$ | -0.1668495 | -0.008283 |
| $b_{13}$ | -1.2191651 | 0.012518 |
| $b_{14}$ | -1.5223832 | -0.068446 |
| $a_2$ | 34.0728264 | 2.071364 |
| $b_{21}$ | -0.5711864 | -0.054005 |
| $b_{22}$ | -0.1142499 | -0.025029 |
| $b_{23}$ | -0.8100883 | 0.065384 |
| $b_{24}$ | -0.9677829 | -0.051774 |
| $a_3$ | 103.292766 | 0.327628 |
| $b_{31}$ | -1.7809655 | -0.022297 |
| $b_{32}$ | -0.3029228 | -0.008557 |
| $b_{33}$ | -2.6887155 | -0.002260 |
| $b_{34}$ | -2.8840097 | 0.010410 |
| $a_4$ | 30.8332588 | -1.318063 |
| $b_{41}$ | -0.4422159 | 0.036978 |
| $b_{42}$ | -0.1141606 | 0.014019 |
| $b_{43}$ | -0.770332 | -0.029955 |
| $b_{44}$ | -0.9159577 | 0.029689 |

**Table S6.1.** Parameter values and initial conditions estimated for the ‘grey whales’ dataset (Gerber, Demaster, & Kareiva, 1999)

Results generated with ALVI-MI with 3DF-spline and a data sample composed of spline values in 1959 and 1966.
| Parameter | True value | Estimate |
| --- | --- | --- |
| **ALVI-MI** |  |  |
| $a_1$ | | 0.0948 |
| $b_{11}$ | | -3.79E-06 |
| $X_1(0)$ | 2894 | 3663.9550 |
| **MAR** |  |  |
| $\alpha_1$ | | 1260 |
| $\beta_{11}$ | | 0.943 |
| $\delta_1$ | | 7240000 |
| $X_1(0)$ | 2894 | |
| **MAR with log transformation** |  |  |
| $\alpha_1$ | | 1.0368 |
| $\beta_{11}$ | | 0.9512 |
| $\delta_1$ | | 0.0327 |
| $X_1(0)$ | 2894 | |
| **MAR with smoothing** |  |  |
| $\alpha_1$ | | 597 |
| $\beta_{11}$ | | 0.9930 |
| $\delta_1$ | | 199000 |
| $X_1(0)$ | 2894 | |
| **MAR with log transformation and smoothing** |  |  |
| $\alpha_1$ | | 0.4902 |
| $\beta_{11}$ | | 0.9535 |
| $\delta_1$ | | 0.0014 |
| $X_1(0)$ | 2894 | |

**Table S6.2.** Parameters and initial conditions estimated for the ‘Wolves and Moose’ dataset (Vucetich, 2021)

**ALVI-MI**
|  | Initial<br>condition | Estimate |
| --- | --- | --- |
| $X_1$ | 20 | 21.6545 |
| $X_2$ | 538 | 560.5340 |
| Parameter |  | Estimate |
| $a_1$ | | -0.2732 |
| $b_{11}$ | | 0.0073 |
| $b_{12}$ | | 0.0001 |
| $a_2$ | | 0.8431 |
| $b_{21}$ | | -0.0380 |
| $b_{22}$ | | -0.0001 |

**MAR with log transformation**
|  | Initial<br>condition | Estimate |
| --- | --- | --- |
| X1 | 20 | 22.0000 |
| X2 | 538 | 564.0000 |
| Parameter |  | Estimate |
| $\beta_{11}$ | | 0.7670 |
| $\beta_{21}$ | | -0.1788 |
| $\beta_{12}$ | | 0.0783 |
| $\beta_{22}$ | | 0.8277 |
| $\delta_1$ | | 0.4485 |
| $\delta_2$ | | 0.1758 |

## References

Certain, G., Barraquand, F., & Gårdmark, A. (2018). How do MAR(1) models cope with hidden nonlinearities in ecological dynamics? Methods in Ecology and Evolution, 9(9), 1975–1995. doi:10.1111/2041-210X.13021

Chou, I. C., & Voit, E. O. (2009). Recent developments in parameter estimation and structure identification of biochemical and genomic systems. Mathematical Biosciences, 219(2), 57–83. doi:10.1016/j.mbs.2009.03.002

Cleveland, W. S. (1981). LOWESS: A program for smoothing scatterplots by robust locally weighted regression. The American Statistician, 35(1), 54. doi:10.2307/2683591

Dam, P., Fonseca, L. L., Konstantinidis, K. T., & Voit, E. O. (2016). Dynamic models of the complex microbial metapopulation of lake mendota. Npj Systems Biology and Applications, 2(1), 16007. doi:10.1038/npjsba.2016.7

Dam, P., Rodriguez-R, L. M., Luo, C., Hatt, J., Tsementzi, D., Konstantinidis, K. T., & Voit, E. O. (2020). Model-based comparisons of the abundance dynamics of dacterial communities in two lakes. Scientific Reports, 10(1), 1–12. doi:10.1038/s41598-020-58769-y

Dennis, B., & Taper, M. L. (1994). Density Dependence in Time Series Observations of Natural Populations: Estimation and Testing. Ecological Monographs, 64(2), 205–224. doi:10.2307/2937041

Gause, G. F. (1934). Experiemental analysis of Vito Volterra’s mathematical theory of the struggle for existence. Science, 79(2036), 16–17. doi:10.1126/science.79.2036.16-a

Gavin, C., Pokrovskii, A., Prentice, M., & Sobolev, V. (2006). Dynamics of a Lotka-Volterra type model with applications to marine phage population dynamics. Journal of Physics: Conference Series, 55(1), 80–93. doi:10.1088/1742-6596/55/1/008

Gerber, L. R., Demaster, D. P., & Kareiva, P. M. (1999). Gray whales and the value of monitoring data in implementing the U.S. endangered species act. Conservation Biology, 13(5), 1215–1219. doi:10.1046/j.1523-1739.1999.98466.x

Holmes, E. E., Ward, E. J., & Scheuerell, M. D. (2020). Analysis of multivariate timeseries using the MARSS package. Seattle: Northwest Fisheries Science Center, NOAA. Retrieved from https://cran.r-project.org/package=MARSS/vignettes/UserGuide.pdf

Holmes, E. E., Ward, E. J., & Wills, K. (2012). MARSS: multivariate autoregressive state-space models for analyzing time-series data. The R Journal, 4(1), 11. doi:10.32614/RJ-2012-002

Huffaker, C. B., Shea, K. B., & Herman, S. G. (1963). Experimental studies on predation: dispersion factors and predator-prey oscillations. Hilgardia, 34, 305–330. Retrieved from http://hilgardia.ucanr.edu/fileaccess.cfm?article=152594&p=ZPTIMD

Ives, A. R. (1995). Predicting the response of populations to environmental change. Ecology, 76(3), 926– 941. doi:10.2307/1939357

Knowles, I., & Renka, R. J. (2014). Methods for Numerical Differentiation of Noisy Data. Electronic Journal of Differential Equations, 21, 235–246. Retrieved from https://ejde.math.txstate.edu/conf-proc/21/k3/knowles.pdf

Lotka, A. J. (1925). Elements of Physical Biology. Baltimor: Williams & Wilkins Company. Retrieved from https://archive.org/details/elementsofphysic017171mbp

May, R. M. (2001). Stability and complexity in model ecosystems. Princeton University Press. doi:10.1515/9780691206912

McLaren, B. E., & Peterson, R. O. (1994). Wolves, moose, and tree Rings on Isle Royale. Science, 266(5190), 1555–1558. doi:10.1126/science.266.5190.1555

Mendes, P., & Kell, D. (1998). Non-linear optimization of biochemical pathways: applications to metabolic engineering and parameter estimation. Bioinformatics, 14(10), 869–883. doi:10.1093/bioinformatics/14.10.869

Mühlbauer, L. K., Schulze, M., Harpole, W. S., & Clark, A. T. (2020). gauseR: Simple methods for fitting Lotka-Volterra models describing Gause’s “Struggle for Existence”. Ecology and Evolution, 10(23), 13275–13283. doi:10.1002/ece3.6926

Park Service, N. (2021). Why relocate wolves to Isle Royale? Retrieved from https://home.nps.gov/isro/learn/why-relocate-wolves-to-isle-royale.htm

Rykiel, E. J. (1996). Testing ecological models: the meaning of validation. Ecological Modelling, 90(3), 229–244. doi:10.1016/0304-3800(95)00152-2

Sachs, A., & Goetze, A. (1945). Mathematical cuneiform texts: 29 (American Oriental). (O. Neugebauer, Ed.). New Haven: American Oriental Society. Retrieved from https://www.jstor.org/stable/1359232?seq=1

Savageau, M. A. (1979). Allometric morphogenesis of complex systems: Derivation of the basic equations from first principles. Proceedings of the National Academy of Sciences, 76(12), 6023– 6025. doi:10.1073/pnas.76.12.6023

Shenhav, L., Furman, O., Briscoe, L., Thompson, M., Silverman, J. D., Mizrahi, I., & Halperin, E. (2019). Modeling the temporal dynamics of the gut microbial community in adults and infants. PLOS Computational Biology, 15(6), e1006960. doi:10.1371/journal.pcbi.1006960

Sims, C. A. (1980). Macroeconomics and reality. Econometrica, 48(1), 1. doi:10.2307/1912017

Stein, R. R., Bucci, V., Toussaint, N. C., Buffie, C. G., Rätsch, G., Pamer, E. G., … Xavier, J. B. (2013). Ecological modeling from time-series inference: insight into dynamics and stability of intestinal microbiota. PLoS Computational Biology, 9(12), e1003388. doi:10.1371/journal.pcbi.1003388

Varah, J. M. (1982). A spline least squares method for numerical parameter estimation in differential equations. SIAM Journal on Scientific and Statistical Computing, 3(1), 28–46. doi:10.1137/0903003

Voit, E. O., & Almeida, J. (2004). Decoupling dynamical systems for pathway identification from metabolic profiles. Bioinformatics, 20(11), 1670–1681. doi:10.1093/bioinformatics/bth140

Voit, E. O., & Chou, I.-C. (2010). Parameter estimation in canonical biological systems models. International Journal of Systems and Synthetic Biology, 1(June), 1–19.

Voit, E. O., Davis, J. D., & Olivença, D. V. (2021). Inference and validation of the structure of Lotka-Volterra models. BioRxiv. doi:10.1101/2021.08.14.456346

Voit, E. O., & Savageau, M. A. (1982). Power-law approach to modeling biological systems; II. Application to ethanol production. J. Ferment. Technol., 60(3), 229–232.

Volterra, V. (1926). Variazioni flultuazioni del numero d’invididui in specie convirenti. Men Acad Lincei. Vucetich, J. A. (2021). Wolves and moose of Isle Royale. Retrieved from https://isleroyalewolf.org/

Wedelin, D., & Gennemark, P. (2007). Efficient algorithms for ordinary differential equation model identification of biological systems. IET Systems Biology, 1(2), 120–129. doi:10.1049/iet-syb:20050098

## References

Batista Júnior, A. B., & Pires, P. S. M. (2014). An Approach to Outlier Detection and Smoothing Applied to a Trajectography Radar Data. Journal of Aerospace Technology and Management, 6(3), 237–248. doi:10.5028/jatm.v6i3.325

Burden, R. L., Faires, J. D., & Burden, A. M. (1993). Numerical Analysis (Fifth edit). Boston, MA: PWS Publishing Co.

Chiang, S.-Y. (2012). An application of Lotka–Volterra model to Taiwan’s transition from 200mm to 300mm silicon wafers. Technological Forecasting and Social Change, 79(2), 383–392. doi:10.1016/j.techfore.2011.05.007

Cleveland, W. S. (1979). Robust Locally Weighted Regression and Smoothing Scatterplots. Journal of the American Statistical Association, 74(368), 829–836. doi:10.1080/01621459.1979.10481038

Cleveland, W. S. (1981). LOWESS: A Program for Smoothing Scatterplots by Robust Locally Weighted Regression. The American Statistician, 35(1), 54. doi:10.2307/2683591

Cleveland, W. S., & Devlin, S. J. (1988). Locally Weighted Regression: An Approach to Regression Analysis by Local Fitting. Journal of the American Statistical Association, 83(403), 596–610. doi:10.1080/01621459.1988.10478639

Cleveland, W. S., & Grosse, E. (1991). Computational methods for local regression. Statistics and Computing, 1(1), 47–62. doi:10.1007/BF01890836

Eilers, P. H. C. (2003). A Perfect Smoother. Analytical Chemistry, 75(14), 3631–3636. doi:10.1021/ac034173t

Eilers, P. H. C., & Marx, B. D. (1996). Flexible smoothing with B-splines and penalties. Statistical Science, 11(2), 89–121. doi:10.1214/ss/1038425655

Gandolfo, G. (2008). Giuseppe Palomba and the Lotka-Volterra equations. RENDICONTI LINCEI, 19(4), 347–357. doi:10.1007/s12210-008-0023-7

Garcia, D. (2010). Robust smoothing of gridded data in one and higher dimensions with missing values. Computational Statistics & Data Analysis, 54(4), 1167–1178. doi:10.1016/j.csda.2009.09.020

Gause, G. F. (1934). Experiemental Analysis of Vito Volterra’s mathematical theory of the struggle for existence. Science, 79(2036), 16–17. doi:10.1126/science.79.2036.16-a

Haas, C. N. (1981). Application of predator-prey models to disinfection. Journal of the Water Pollution Control Federation, 53(3 I), 378–386. doi:10.2307/25041087

Hacinliyan, A. S., Kusbeyzi, I., & Aybar, O. O. (2010). Approximate solutions of Maxwell Bloch equations and possible Lotka Volterra type behavior. Nonlinear Dynamics, 62(1–2), 17–26. doi:10.1007/s11071-010-9695-5

Holmes, E. E., Ward, E. J., & Scheuerell, M. D. (2020). Analysis of multivariate timeseries using the MARSS package, version 3.11.3.

Hung, H.-C., Chiu, Y.-C., Huang, H.-C., & Wu, M.-C. (2017). An enhanced application of Lotka–Volterra model to forecast the sales of two competing retail formats. Computers & Industrial Engineering, 109, 325–334. doi:10.1016/j.cie.2017.05.022

Hytti, H., Takalo, R., & Ihalainen, H. (2006). Tutorial on Multivariate Autoregressive Modelling. Journal of Clinical Monitoring and Computing, 20(2), 101–108. doi:10.1007/s10877-006-9013-4

Loader, C. (2012). Smoothing: Local Regression Techniques. In Handbook of Computational Statistics (pp. 571–596). Berlin, Heidelberg: Springer Berlin Heidelberg. doi:10.1007/978-3-642-21551-3_20

Nambu, M. (1986). Plasma-maser effects in plasma astrophysics. Space Science Reviews, 44(3–4), 357– 391. doi:10.1007/BF00200820

Peschel, M., & Mende, W. (1986). The Predator-Prey Model: Do we Live in a Volterra World? Berlin: Akademie-Verlag.

Ramsay, J. O., Hooker, G., Campbell, D., & Cao, J. (2007). Parameter estimation for differential equations: a generalized smoothing approach. Journal of the Royal Statistical Society: Series B (Statistical Methodology), 69(5), 741–796. doi:10.1111/j.1467-9868.2007.00610.x

Savageau, M. A., & Voit, E. O. (1987). Recasting nonlinear differential equations as S-systems: a canonical nonlinear form. Mathematical Biosciences, 87(1), 83–115. doi:10.1016/0025-5564(87)90035-6

Smyth, G. (2020). Difference between LOESS and LOWESS. Retrieved 3 September 2021, from https://stats.stackexchange.com/questions/161069/difference-between-loess-and-lowess

Torres, N. V., & Voit, E. O. (2002). Pathway Analysis and Optimization in Metabolic Engineering. Cambridge, U.K.: Cambridge University Press. doi:10.1017/CBO9780511546334

Vano, J. A., Wildenberg, J. C., Anderson, M. B., Noel, J. K., & Sprott, J. C. (2006). Chaos in low-dimensional Lotka–Volterra models of competition. Nonlinearity, 19(10), 2391–2404. doi:10.1088/0951-7715/19/10/006

Vilela, M., Borges, C. C. H., Vinga, S., Vasconcelos, A. T. R., Santos, H., Voit, E. O., & Almeida, J. S. (2007). Automated smoother for the numerical decoupling of dynamics models. BMC Bioinformatics, 8(1), 305. doi:10.1186/1471-2105-8-305

Voit, E. O. (2000). Canonical Modeling: Review of Concepts with Emphasis on Environmental Health. Environmental Health Perspectives, 108(s5), 895–909. doi:10.1289/ehp.00108s5895

Voit, E. O. (2013). Biochemical systems theory: a review. ISRN Biomathematics, 2013, 1–53. doi:10.1155/2013/897658

Voit, E. O. (2017). *A First Course in Systems Biology* (Second edi). Garland Science. doi:10.1201/9780203702260

Voit, E. O., & Almeida, J. (2003). Dynamic Profiling and Canonical Modeling. In Metabolic Profiling: Its Role in Biomarker Discovery and Gene Function Analysis (pp. 257–276). Boston, MA: Springer US. doi:10.1007/978-1-4615-0333-0_14

Voit, E. O., Marino, S., & Lall, R. (2005). Challenges for the Identification of Biological Systems from in vivo Time Series Data, 5(December 2004), 83–92.

Voit, E. O., & Savageau, M. A. (1982a). Power-law approach to modeling biological systems; II. Application to ethanol production. J. Ferment. Technol., 60(3), 229–232.

Voit, E. O., & Savageau, M. A. (1982b). Power-law approach to modeling biological systems; III. Methods of analysis. J. Ferment. Technol., 60(3), 233–241.

Voit, E. O., & Savageau, M. A. (1986). Equivalence between S-systems and Volterra systems. Mathematical Biosciences, 78(1), 47–55. doi:10.1016/0025-5564(86)90030-1

Vucetich, J. A. (2021). Wolves and moose of Isle Royale. Retrieved from https://isleroyalewolf.org/

Zhou, Y., & Chen, B. (2006). Analysis of multi-ISPs game based on lotka-volterra model. In CIMCA 2006: International Conference on Computational Intelligence for Modelling, Control and Automation, Jointly with IAWTIC 2006: International Conference on Intelligent Agents Web Technologies … IEEE Computer Society. doi:10.1109/CIMCA.2006.44

